# Dual effects of *ARX* poly-alanine mutations in human cortical and interneuron development

**DOI:** 10.1101/2024.01.25.577271

**Authors:** Vanesa Nieto-Estevez, Parul Varma, Sara Mirsadeghi, Jimena Caballero, Sergio Gamero-Alameda, Ali Hosseini, Marc J. Silvosa, Drew M. Thodeson, Zane R. Lybrand, Michele Giugliano, Christopher Navara, Jenny Hsieh

## Abstract

Infantile spasms, with an incidence of 1.6 to 4.5 per 10,000 live births, are a relentless and devastating childhood epilepsy marked by severe seizures but also leads to lifelong intellectual disability. Alarmingly, up to 5% of males with this condition carry a mutation in the *Aristaless-related homeobox* (*ARX*) gene. Our current lack of human-specific models for developmental epilepsy, coupled with discrepancies between animal studies and human data, underscores the gap in knowledge and urgent need for innovative human models, organoids being one of the best available. Here, we used human neural organoid models, cortical organoids (CO) and ganglionic eminences organoids (GEO) which mimic cortical and interneuron development respectively, to study the consequences of PAE mutations, one of the most prevalent mutation in *ARX*. ARX^PAE^ produces a decrease expression of *ARX* in GEOs, and an enhancement in interneuron migration. That accelerated migration is cell autonomously driven, and it can be rescued by inhibiting CXCR4. We also found that PAE mutations result in an early increase in radial glia cells and intermediate progenitor cells, followed by a subsequent loss of cortical neurons at later timepoints. Moreover, *ARX* expression is upregulated in COs derived from patients at 30 DIV and is associated with alterations in the expression of *CDKN1C*. Furthermore, ARX^PAE^ assembloids had hyperactivity which were evident at early stages of development. With effective treatments for infantile spasms and developmental epilepsies still elusive, delving into the role of ARX^PAE^ mutations in human brain organoids represents a pivotal step toward uncovering groundbreaking therapeutic strategies.

## INTRODUCTION

Infantile spasms are a severe and unyielding form of childhood epilepsy characterized by intense seizures and lasting intellectual disability (ID), impacting 1.6-4.5 per 10,000 live births. As many as 5% of affected males have a mutation in the *Aristaless-related homeobox* (*ARX*) gene (*1, 2*)*. ARX* gene is a paired-like homeodomain (HD)-containing transcription factor located on the X-chromosome, capable of functioning as either an activator or a repressor (*3–6*). In addition to the HD, ARX contains a conserved aristaless domain, an octapeptide domain, three nuclear localization sequences, a central acidic domain and four poly-alanine tracts (*6*). Mutations in *ARX* are associated with a broad spectrum of phenotypes that can be divided into two groups. Firstly, mutations in the HD and mutations which produce truncated proteins resulting in severe ID, autism spectrum disorder (ASD), seizures, and brain malformations, notably lissencephaly and agenesis of the corpus callosum (*7–9*). Secondly, missense mutations and in-frame expansions of the first two poly-alanine tracts that are associated with ID, ASD, and epilepsy but no structural brain malformations (*8, 10, 11*). Disease-causing poly-alanine expansion (PAE) mutations have been identified in nine genes, with eight of them, including *ARX*, encoding transcription factors (*12, 13*). Unlike the more common and well-studied poly-glutamine repeats, PAEs are shorter, often less than 20 alanines. They result in developmental disorders similar to those seen with PAE mutations in *ARX*, suggesting similar molecular/genetic mechanisms underlying PAE-related disorders (*12, 13*).

During development, *Arx* is expressed in the nervous system, pancreas and testis. In adult mice, it has been found in brain, muscle, heart, and liver (*14, 15*). In the brain, *Arx* is expressed in both cortical progenitor cells as well as postmitotic cortical ɣ-aminobutyric acid-containing (GABAergic) interneurons. It has been shown to play critical roles during interneuron migration and cortical development (*14, 16, 17*). Studies using knock-out (KO) mouse models or mutations in the HD of *Arx* have recapitulated the phenotype of patients with the most severe cases of ARX-associated epilepsy. Although these mice show aberrant tangential migration and reduced GABAergic interneurons in the cortex and other brain regions (*18, 19*), perinatal lethality has precluded further analysis in adolescent or adult mice (*15*). To overcome this limitation, conditional KO mice lacking *Arx* expression selectively in interneuron progenitor cells were used. These mice showed a similar loss of interneurons (*20–22*), and they developed seizures (*20*).

Moreover, the loss of *Arx* in cortical projection neuron progenitor cells resulted in a proliferation defect and decreased the number of upper-layer cortical neurons (*16, 17, 23*). Interestingly, mice lacking *Arx* in cortical projection neuron precursors did not develop seizures, despite being hyperactive and having behavioral abnormalities (*24*). In contrast, mice with a PAE mutation knocked in showed fewer GABAergic interneurons in the cortex (*19, 25*) and experienced seizures (*26–29*). Although these mouse models have advanced our understanding the role ARX plays in several neurological conditions, they fail to fully recapitulate its complete role in human brain development, as mice lack key aspects of human development. This is particularly evident when mouse models are used to develop new therapies; unfortunately drugs that show effectiveness in animals often fail in clinical trials (*30*). Moreover, epilepsy in many patients with *ARX* mutations is resistant to existing pharmaceutical treatments. Thus, there is a clear need for new models for studying neurological diseases and for drug screening.

Postmortem tissues from patients with these rare neurological diseases are not often available for study, especially from pediatric patients. Neural organoids can serve as excellent *in vitro* models for studying human development because they mimic key aspects of human brain development, such as the presence of outer radial glial cells (oRGCs), expansion of cortical layers and interneuron migration (*31–35*). Importantly, this approach opens up opportunities to develop new treatments for human neurological disorders (*36–38*). To study the role of *ARX* PAE mutations in human brain development and epilepsy, we recruited three male patients with a PAE in the second tract of *ARX*, along with three healthy male controls. We focused on male patients because *ARX* is located on the X Chromosome, and females present with milder and more variable phenotypes (*20*). Next, we generated two types of human regionalized neural organoids from human induced pluripotent stem cells (hiPSCs): cortical organoids (CO) and ganglionic eminences organoids (GEO), using an existing protocol (*32, 35, 39*). COs resemble the cortical plate during development, with progenitor cells located around ventricular-zone-like structures, and cortical neurons differentiating in an inside-out manner (*35, 40*). On the other hand, GEOs contain ventral progenitor cells similar to interneuron progenitor cells in ganglionic eminences (GE) that migrate to the cortex and integrate into the cortical network (*32, 40, 41*). We used immunohistochemistry (IHC) and single-cell RNA sequencing (scRNA-seq) at different developmental timepoints in patient-derived CO and GEO to test the hypothesis that *ARX* PAE mutations would result in cortical and interneuron defects. Our results revealed that PAE decreased the number of ARX^+^ cells in GEO and GABAergic neurons and accelerated the migration of interneurons in the cortex, possibly due to overexpression of CXCR4/CXCL12. Interestingly, CXCR4 inhibition was sufficient to reduce interneuron migration to control levels. On the other hand, ARX was upregulated in CO derived from patients compared to healthy controls at 30 days *in vitro* (DIV). We also observed an increase in radial glial cells (RGCs) and intermediate progenitor cells (IPs) in patient-derived CO at 30 DIV. Gene-expression analysis of these RGCs revealed that they either increased the expression of genes such as *NESTIN* and *HES5*, while downregulated *HES1* and *PAX6*, all canonical markers of RGCs. Moreover, patient-derived CO at 30 DIV showed a decreased expression of *CDKN1C*, a cyclin dependent kinase inhibitor 1C, which negatively regulates cellular proliferation. This activation of RGCs likely led to increased cortical neurogenesis initially, exhausting the RGC population and eventually resulting in decreased cortical neurons later in development, as evidenced by the reduced expression of cortical neuron markers such as *NEUROD1*, *CTIP2* and *SATB2* at 120 DIV. Using multi-electrode array (MEA) recording to measure extracellular activity, we demonstrate these cellular abnormalities can alter neuronal circuitry, leading to hyperactivity. Our findings demonstrate a dual role of ARX during human brain development, offering the first comprehensive comparison of ARX^PAE^ mutations across both human cortical and ganglionic eminence cell types. Our human model with ARX^PAE^ mutations not only mirrors the hyperexcitability observed in human cases but also holds promise for advancing our understanding of the disorder’s pathogenesis and paving the way for the discovery of novel therapeutic strategies.

## RESULTS

### Generation and characterization of CO and GEO

ARX is a key transcription factor expressed during brain development that controls cortical progenitor cell differentiation as well as interneuron migration in animal models (*14, 16, 17*). Although most of what is known about ARX function comes from mouse models, it is unclear if these models fully recapitulate its role during human brain development, thus necessitating the need for human models to study. To address this challenge, we turned to CO and GEO *in vitro* models of human brain development (*32, 35, 41*) to elucidate the role(s) ARX plays in human brain development and in the pathogenesis of epilepsy. To confirm our organoids accurately reproduced the sequence of development observed by others, we used IHC and scRNA-seq. Neural differentiation was achieved through SMAD inhibition to generate COs from hiPSCs (*35*) (**Fig. 1A**). scRNA-seq analysis of 30 DIV COs confirmed the presence of the main cell types in the developing cortex at early stage, including RGCs, cycling progenitor cells and IPs, deep-layer neurons, a population of early Cajal-Retzius cells, and a small percentage of interneurons (**Fig. 1B**). In addition, we observed by IHC that COs at early time points (30 and 60 DIV) had ventricle-like structures, defined by PAX6^+^ (Paired Box 6), SOX2^+^ (SRY-box transcription factor 2) and VIMENTIN^+^ cells [expressed in dorsal cortical progenitor cells (*42, 43*)], surrounded by maturing neurons [labeled with CTIP2 and TBR1, expressed in deep-layer cortical neurons (*44–46*)] (**Fig. 1C-D** and **Fig. S1A**). Similar analysis of CO at 120 DIV revealed the presence of oRGCs marked by expression of *FAM107A* (family with sequence similarity 107 member A) and *HOPX*, key factors expressed in human oRGCs (*47, 48*). We also observed the expression of the cortical neuron markers FEZF2, CTIP2 and TBR1, as well as BRN2 (*POU3F2*), SATB2, CUX1, CUX2, LHX5 and REELIN in COs at 120 DIV by IHC and/or scRNA-seq (**Fig. 1E-F** and **Fig. S1B**). While FEZF2, CTIP2 and TBR1 labeled deep-layer neurons, BRN2 (*POU3F2*), SATB2, CUX1, CUX2, LHX5 and REELIN are expressed in upper-layer neurons (*45, 49–51*). We also generated GEOs resembling the GE by modulating WNT (Wingless-related integration site) and SHH (Sonic Hedgehog) pathways through addition of IWP-2 and SAG (Smoothened receptor agonist) (*32*). We found that GEO at 30 DIV contained KI67^+^ and NKX2.1^+^ progenitor cells (**Fig. 1G-H**). KI67 labels proliferative cells in the G1, S, G2, and M phases (*52*), whereas *NKX2.1* is expressed in progenitor cells in the medial GE and in the preoptic area (*53, 54*). Our analysis of GEOs at 120 DIV, a later developmental timepoint, revealed TUJ1^+^, GABA^+^ and CXCR4^+^ neurons. (**Fig. 1I-J**). CXCR4 (C-X-C motif chemokine receptor 4) has been shown to play an important role in interneuron migration (*55–57*). We also detected cells expressing calbindin (CB), calretinin (CR) and somatostatin (SST, **Fig. 1K**), three interneuron subtypes (*54*). In addition, we found a small percentage of cells expressing gene signatures of microglia and oligodendrocyte progenitor cells at 30 and 120 DIV, respectively. Heatmaps and featureplots showing the specific set of genes expressed by each cluster/cell type identified by scRNA-seq in CO and GEO at 30 and 120 DIV are shown in **Fig. S2-S6** and supplemental excel file. These results corroborate the generation of neural organoids resembling cortical and subpallium regions of human brain as described previously (*32, 35, 40*).

**Figure 1.**
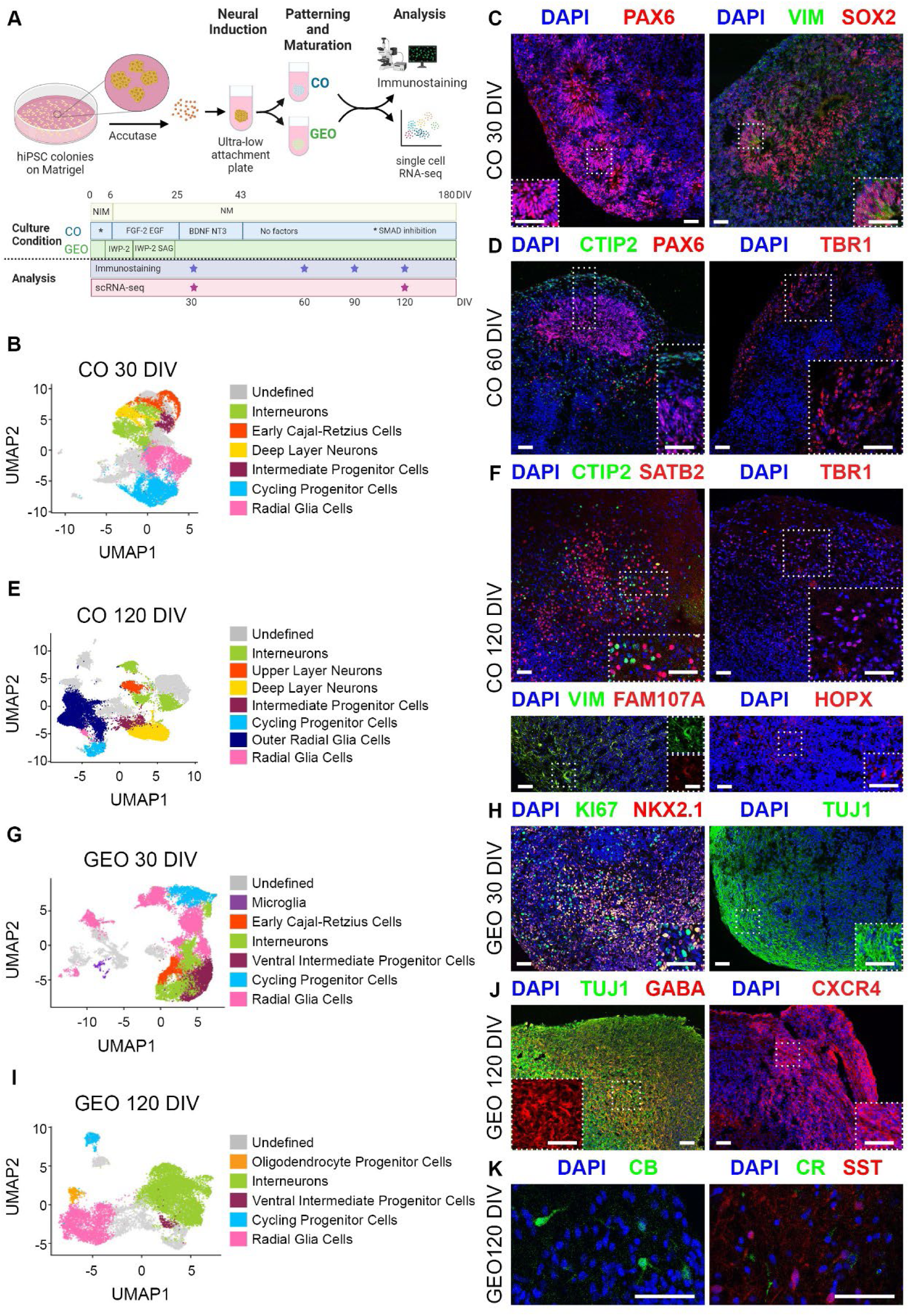
The generation and characterization of human organoids. **(A)** Experimental design and culture conditions to obtain COs and GEOs from hiPSCs. Organoids were analyzed by immunostaining at 30, 60, 90, and 120 DIV and by scRNA-sequencing at 30 and 120 DIV. **(B)** UMAP plot from control COs at 30 DIV. **(C)** The images show control COs at 30 DIV immunostained against PAX6, VIMENTIN and SOX2; and stained with DAPI. **(D)** The images show control COs at 60 DIV immunostained against PAX6, CTIP2 and TBR1; and stained with DAPI. **(E)** UMAP plot from control COs at 120 DIV. (**F)** The images show control COs at 120 DIV immunostained against CTIP2, SATB2, TBR1, VIMENTIN, FAM107A and HOPX; and stained with DAPI. **(G)** UMAP plot from control GEOs at 30 DIV. **(H)** The images show control GEOs at 30 DIV immunostained against KI67, NKX2.1 and TUJ1; and stain with DAPI. **(I)** UMAP plot from control GEOs at 120 DIV. **(J)** The images show control GEOs at 120 DIV immunostained against TUJ1, GABA and CXCR4; and stain with DAPI. **(K)** The images show control GEOs at 120 DIV immunostained against CB, CR and SST; and stain with DAPI. Scale bar = 50 µm. DIV = days *in vitro*, CB = Calbindin, CR = Calretinin, COs = cortical organoids, GEOs = ganglionic eminence organoids, NIM = Neural induction media, NM = Neural media, SST = Somatostatin, VIM = VIMENTIN. N= 18 organoids from 2 clones x 3 lines per condition.

### Expression of ARX and impact of PAE in CO and GEO

We first interrogated our scRNA-seq dataset and found that ARX is expressed in control COs and GEOs at 30 and 120 DIV (**Fig. 2A-B**). As predicted, a higher percentage of *ARX*-expressing cells was found in GEO when compared to CO. Furthermore, and consistent with known data, for the cells expressing *ARX* in CO, a high percentage was proliferative at the earlier time point (**Fig. 2A**) (*14*). We also observed cortical neurons expressing *ARX* in COs at both timepoints. In GEO, *ARX* was expressed in both progenitor cells and GABAergic neurons (**Fig. 2B**). We further confirmed the expression of ARX in progenitor cells (Ki67) and in neurons (doublecortin, DCX, and TUJ1) in COs and GEOs by IHC (**Fig. 2C-D**). However, the lack of compatible antibodies prevented us to analyze the co-localization of ARX and other specific markers (e.g. PAX6, SOX2, NKX2.1, GABA, etc.). These data were consistent with the expression of *ARX* in human brain tissue (**Fig. S7**). Thus, during embryonic development, *ARX* expression is higher in the GE/striatum than in the cortex, similar to data from mice (*14*). Furthermore, *ARX*-expressing cells can be found both in the cortex and the striatum throughout adult life, although the level of expression is lower than in embryonic tissues.

**Figure 2.**
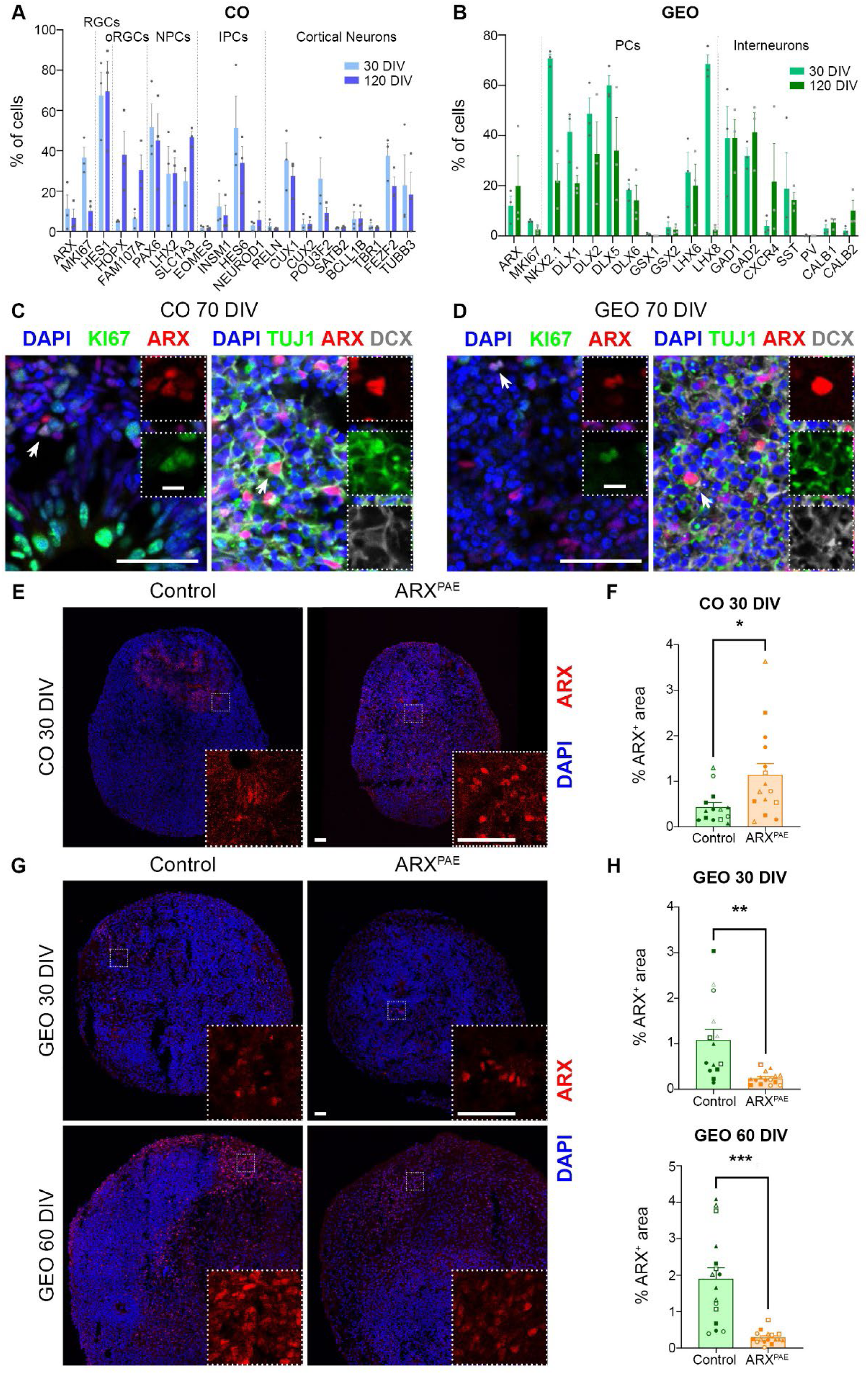
ARX expression in cortical and ganglionic eminence organoids. **(A)** Graph shows the percentage of ARX-expressing cells (number of ARX^+^ cells/total number of cells) and the percentage of cells expressing ARX (number of ARX^+^ cells expressing X marker/total number of ARX^+^ cells) in COs at 30 and 120 DIV using scRNA-seq. **(B)** Graph shows the percentage of ARX expressing cells (number of ARX^+^ cells/total number of cells) and the percentage of cells expressing ARX (number of ARX^+^ cells expressing X marker/total number of ARX^+^ cells) in GEOs at 30 and 120 DIV using scRNA-seq. **(C)** The images show control COs immunostained against ARX, KI67, DCX and TUJ1; and stained with DAPI. **(D)** The images show control GEOs immunostained against ARX, KI67, DCX and TUJ1; and stained with DAPI. The images show COs **(E)** and GEOs **(G)** derived from control and ARX^PAE^ hiPSCs immunostained against ARX and stained with DAPI. Graphs show the percentage of ARX^+^ area in COs **(F)** and GEOs **(H)**. Scale bar = 50 µm. Inserts (C-D) = 10 µm. DIV = days *in vitro*, COs = cortical organoids, GEOs = ganglionic eminence organoids, IPCs = intermediate progenitor cells, NPCs = neural progenitor cells, oRGCs = outer radial glial cells, PCs = progenitor cells, RGCs = radial glial cells. Two-tailed Student’s *t*-test, * = p < 0.05, ** = p < 0.01, *** = p < 0.001. The results are the mean ± SEM from N= 13-16 organoids from 2 clones x 3 lines per condition. Individual points represented data from individual organoid and each symbol represented the data from each clone/line (Line 1: Clone A ● Clone B ○; Line 2: Clone A ◼ Clone B ◻; Line 3: Clone A ▲ Clone B △).

To study the role of ARX during human development, we recruited three healthy controls and three patients with eight additional alanines in the second poly-alanine tract of *ARX* (**Table S1**). Six lines of hiPSCs were generated (one per person) and two clones per line were used in all experiments (or as otherwise indicated). All the lines expressed pluripotency markers (NANOG, SOX2, OCT4, LIN28, SSEA-4 and TRA-1-6), lost reprogramming factors (as shown by the lack of expression of the vectors used for reprograming) and maintained normal karyotype throughout the study (**Fig. S8**). The number of alanines was confirmed by PCR amplification followed by Sanger sequencing (data not shown). Then, we first assayed ARX expression in COs and GEOs derived from patients (**Fig. 2E-H**). We found a significant increase in ARX expression among cells in ARX^PAE^ COs at 30 DIV when compared to controls by IHC (P=0.0177, d=0.944, n=13-14; **Fig. 2E-F**) and scRNA-seq analysis (**Fig. S9**) which was consistent across all three lines (**Fig. S10**). No differences were found at later time points 60, 90 or 120 DIV COs (**Fig. S11A-C**). In contrast, ARX^+^ cells decreased in ARX^PAE^ GEOs at 30 and 60 DIV [30 DIV: P= 0.0033, d=-1.367, n=14-15; 60 DIV: P=0.0001, d=-1.823, n=16; **Fig. 2G-H**, **Fig. S12** (plots per condition) and **S13** (plots per line)], but we did not see any significant difference at 120 DIV (**Fig. S11D**). Although it has been reported that PAE produces nuclear inclusions and/or cytoplasmatic localization (*28, 58–60*), ARX was restricted to the nucleus without forming inclusions in COs and GEOs (see inserts). Our data demonstrated that ARX is expressed in cortical and subpallium tissues and that PAE affects *ARX* expression differently in COs and GEOs.

### PAE alters interneuron differentiation and migration

As we found higher ARX expression in GEOs than in COs, we studied the effect of PAE in GEO at different timepoints. We first analyzed the size of control and ARX^PAE^ GEOs over time and found that ARX^PAE^ GEOs were larger than controls at all time points (**Fig. S14**, 14DIV: P= 0.0084, 21DIV: P= 0.0004, 60DIV: P= 0.0059). No main differences were found in the number of progenitor cells between control and ARX^PAE^ GEOs at 30 DIV by scRNA-seq (**Fig. 3A**) with just a trend towards fewer KI67^+^ proliferative cells in ARX^PAE^ GEOs at 30 DIV (P= 0.069, d=-0.672, n=14-18; **Fig. 3B-C**), but no change in NKX2.1 (**Fig. 3B-C**). Although the interneuron cluster was decrease in ARX^PAE^ GEOs at 30 DIV (**Fig. 3A**), no differences were detected in DCX^+^ or TUJ1^+^ cells by IHC (**Fig. 3D**). To further explore the interneuron population, we examined neuronal markers (DCX, TUJ1 and GABA) at later time points. While we found a trend towards a decrease in DCX^+^ cells at 60 DIV in ARX^PAE^ GEOs (P= 0.0589, d=-0.672, n=17; **Fig. 3E**) and a significant decrease in GABA^+^ at 90 DIV (P= 0.0219, d=-1.206, n=10; **Fig. 3F**), the main differences were found at 120 DIV when ARX^PAE^ GEOs had fewer TUJ1^+^ and GABA^+^ neurons (TUJ1: P= 0.0154, d=-1.196, n=8-13; GABA: P=0.0641, d=-0.85, n=9-13; **Fig. 3G-H**). Similar reduction in interneurons was also observed in the scRNA-seq data at 120 DIV (**Fig. 3I**; and **Fig.S15**, plots per condition, and **Fig. S16**, plots per line). Although ARX^PAE^ GEOs had fewer interneurons, they exhibited higher expression levels of *CALB1*, *CALB2* and *SST* (**Fig. S15**, plots per condition; and **Fig. S16**, plots per line). Although the number of *PV* expressing cells was higher in ARX^PAE^ GEOs compared to controls, the number of cells was low in both conditions. This is expected as this subtype of interneurons is produced later during development as compared to the other subtypes (*54*). To determine if the reduction of interneurons in ARX^PAE^ GEOs were due to an increase of cell death, we analyzed the number of AC3^+^ cells by IHC. We did not observe any significant difference in AC3^+^ cells in ARX^PAE^ GEOs compared to controls at any of the time points analyzed, except for a decrease at 60 DIV (**Fig. S17**). Altogether, these data suggest PAE affects interneuron differentiation during development without affecting progenitor cell production.

**Figure 3.**
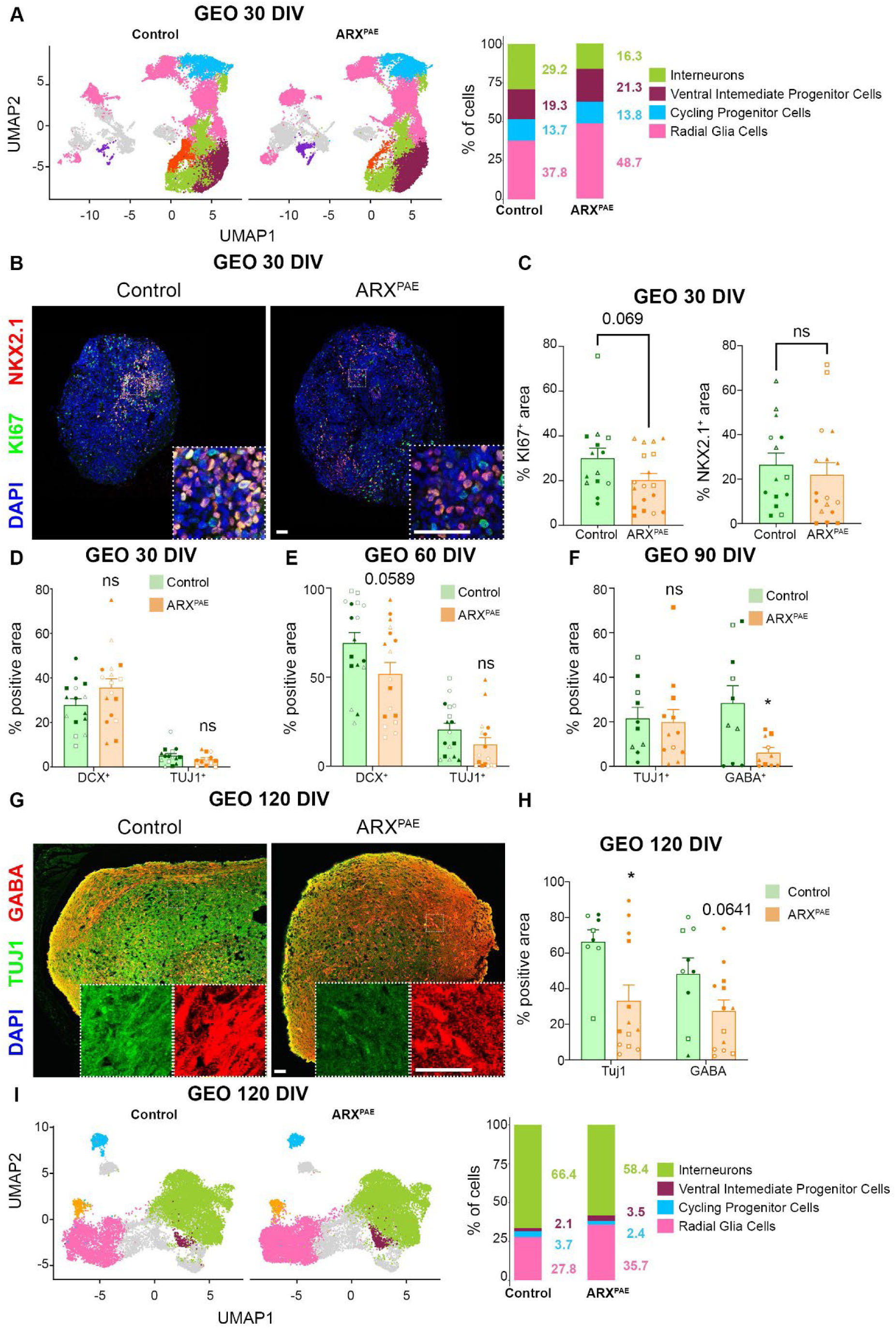
Poly-Alanine expansion affects GABAergic differentiation in ganglionic eminence organoids. **(A)** UMAP plots from control and ARX^PAE^ GEOs at 30 DIV. Bar graph shows the percentage of cells in each cluster. **(B)** The images show control and ARX^PAE^ GEOs at 30 DIV immunostained against KI67 and NKX2.1, and stained with DAPI. **(C)** Graphs show the percentage of KI67^+^ and NKX2.1^+^ area in control and ARX^PAE^ GEOs at 30 DIV. **(D)** Graph shows the percentage of DCX^+^ and TUJ1^+^ area in control and ARX^PAE^ GEOs at 30 DIV. **(E)** Graph shows the percentage of DCX^+^ and TUJ1^+^ area in control and ARX^PAE^ GEOs at 60 DIV. **(F)** Graph shows the percentage of TUJ1^+^ and GABA^+^ area in control and ARX^PAE^ GEOs at 90 DIV. **(G)** The images show control and ARX^PAE^ GEOs at 120 DIV immunostained against TUJ1 and GABA; and stained with DAPI. **(H)** Graphs show the percentage of TUJ1^+^ and GABA^+^ area in control and ARX^PAE^ GEO at 120 DIV. **(I)** UMAP plots from control and ARX^PAE^ GEOs at 120 DIV. Bar graph shows the percentage of cells in each cluster. Scale bar = 50 µm. DIV = days *in vitro*, GEOs = ganglionic eminence organoids. Two-tailed Student’s *t*-test, * = p < 0.05, ns = not significant. The results are the mean ± SEM from N= 8-14 organoids from 2 clones x 3 lines per condition. Individual points represented data from individual organoid and each symbol represented the data from each clone/line (Line 1: Clone A ● Clone B ○; Line 2: Clone A ◼ Clone B ◻; Line 3: Clone A ▲ Clone B △).

During development, interneurons are formed in the GE and migrate along circuitous routes to the cortex to integrate with projection neurons and form the cortical neuronal circuits (*54, 61*). Organoid models can recapitulate interneuron migration by fusing one GEO and one CO (referred as assembloids). We labeled interneuron progenitor cells in GEOs at 50 DIV using a DLX1/2b GFP-expressing lentivirus (**Fig. 4A**). DLX1 and DLX2 are expressed in ventral progenitor cells and promote interneuron differentiation and migration (*54, 62*). We also labeled COs using an AAV-synapsin-mCherry virus; these mCherry^+^ cells did not migrate into GEOs (**Fig. S18A**). After 10 DIV, one GEO was placed in close contact with one CO to allow their fusion and the migration of the DLX1/2-GFP^+^ cells from the GEO to the CO. Migration into the CO was assayed at 30 days after fusion by time-lapse imaging. We observed a similar percentage of cells moving in ARX^PAE^ assembloids (condition 2) compared to controls (condition 1, **Fig. S18B**). However, DLX1/2-GFP^+^ cells migrated further and faster in ARX^PAE^ assembloids (Distance: P= 0.0110, d=0.607, n=38-48; Velocity: P= 0.0090, d=0.624, n=38-47; **Fig. 4B-F** and **videos S1-4**). Moreover, in control assembloids most of the cells migrated less than 300 µm, whereas a higher percentage of cells migrated more than 300 µm in ARX^PAE^ assembloids (**Fig. 4G**). To determine if the accelerated migration observed ARX^PAE^ assembloids was due to a premature migration, we analyzed the percentage of mobile/immobile DLX1/2-GFP^+^ cells at earlier time point (75 DIV). We observed a smaller percentage of mobile cells at 75 DIV compared to 90 DIV, but no differences were found in ARX^PAE^ assembloids compared to controls (**Fig. S18C**). To investigate if the faster migration of interneurons leads to an increase of neurons in the cortex, we quantified the GFP^+^ area in the CO and GEO side of the assembloids, and both ARX^PAE^ and control assembloids showed similar percentages of GFP^+^ area (**Fig. S18D**). DLX1/2-GFP^+^ cells, both in control and ARX^PAE^ assembloids, expressed GABA after migrating into the COs (**Fig. S19A**), but they did not express CB or CR (**Fig. S19B**). To determine whether the accelerated migration observed in ARX^PAE^ assembloids was due to a cell-autonomous effect, we fused ARX^PAE^ GEOs to control COs (condition 3) to create inter-individual assembloids, where interneurons would have PAE and the cortical side would express wild-type ARX. We found DLX1/2-GFP^+^ cells migrated a similar distance and had a similar velocity than ARX^PAE^ assembloids (Distance: P= 0.0125, d=0.522, n=48; Velocity: P= 0.0081, d=0.53, n=47-48, **Fig. 4D-G**). To test whether PAE in cortical cells was sufficient to affect interneuron migration, we fused control GEOs to ARX^PAE^ COs (condition 4). Interneurons in this condition migrated at a velocity and distance similar to those observed in control assembloids (Distance: P= 0.0370, d=-0.52, n=30-38; Velocity: P= 0.0423, d=-0.52, n=30-38; **Fig.4D-G**). These data suggested that PAE enhanced interneuron migration in a cell-autonomous manner.

**Figure 4.**
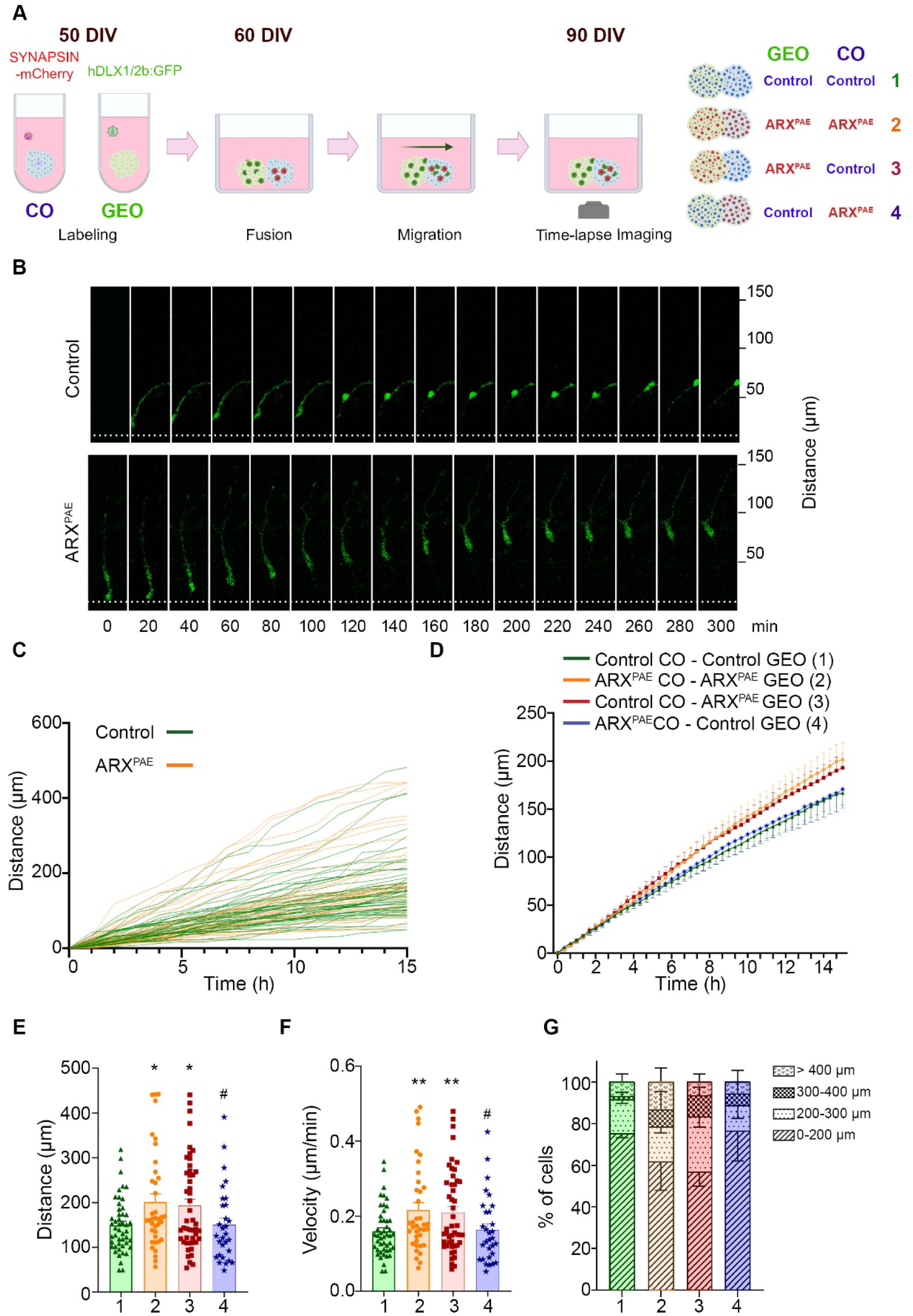
Poly-Alanine expansion in ARX accelerates interneuron migration in assembloids. **(A)** Experimental design: GEOs were labeled using a lenti-DLX1/2b:GFP virus at 50 DIV. COs were labeled using an AAV-SYNAPSIN-mCherry. After 10 days, one CO and one GEO were fused and neuronal migration were observed over 15 hours at 90 DIV. Four conditions were studied: (1) Control CO + Control GEO, (2) ARX^PAE^ CO + ARX^PAE^ GEO, (3) Control CO + ARX^PAE^ GEO, (4) ARX^PAE^ CO + Control GEO. **(B)** Representative images of DLX1/2-GFP^+^ interneurons over time in control and ARX^PAE^ assembloids. **(C)** The graph shows single tracks of DLX1/2-GFP+ interneurons over time in control and ARX^PAE^ assembloids. **(D)** The graph shows the mean distance at each timepoint of DLX1/2-GFP^+^ interneurons over time in all four experimental conditions. **(E)** The graph shows the mean of the total distance of DLX1/2-GFP^+^ interneurons in all four conditions. **(F)** The graph shows the velocity of DLX1/2-GFP^+^ interneurons in all four conditions. **(G)** The graph shows the distribution of the total distance of DLX1/2-GFP^+^ interneurons in all four experimental conditions. The results are the mean ± SEM from N= 30-48 neurons from 11-17 organoids from 2 clones x 3 lines per condition. * shows statistical analysis compared to controls. # shows statistical analysis compared to ARX^PAE^ assembloids. Two-tailed Student’s *t*-test, *, ^#^ = p < 0.05, **, ^##^ = p < 0.01, ns = not significant. DIV = days *in vitro*, COs = cortical organoids, GEOs = ganglionic eminence organoids.

As CXCR4 is important for neuronal migration (*63*) and it is a direct target of ARX, we checked if DLX1/2-GFP^+^ cells expressed CXCR4. DLX1/2-GFP^+^ cells expressed CXCR4 before and after migrating to the COs both in control and ARX^PAE^ assembloids at 90 DIV (**Fig. 5A**). Then, we determined if CXCR4 expression was altered due to PAE, and we found an increase in CXCR4 in ARX^PAE^ GEOs at 120 DIV by IHC (P= 0.0037, d=1.313, n=10-13; **Fig. 5B-C**) and scRNA-seq analysis (**Fig. S15**, plots per condition; and **Fig. S16**, plots per line). These results prompted us to explore whether CXCR4 inhibition is sufficient to reduce the enhanced migration observed in ARX^PAE^ assembloids. To test this hypothesis, after a 15-hour baseline recording, control and ARX^PAE^ assembloids were exposed to AMD3100, a CXCR4 inhibitor, and interneuron migration was evaluated for another 15 hours (**Fig. 5D**). Although AMD3100 did not affect interneuron migration in control assembloids, it reduced migration in ARX^PAE^ assembloids to levels comparable to controls (P= 0.0010, n=63-66; **Fig. 5E**). Overall, our results indicate that PAE affects interneuron differentiation and promotes migration which can be reduced by inhibiting CXCR4 expression.

**Figure 5.**
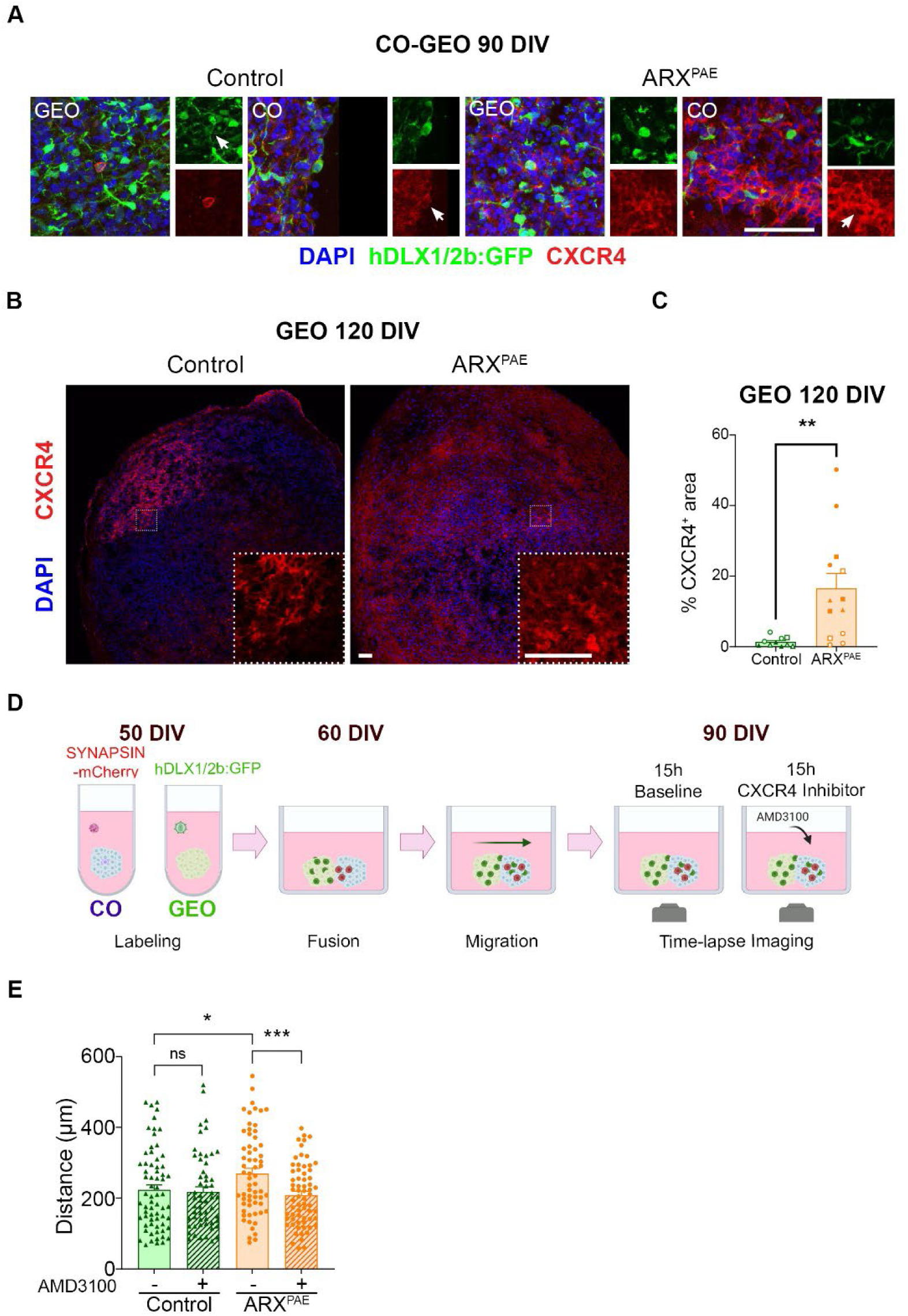
CXCR4 inhibition is sufficient to decrease interneuron migration in ARX^PAE^ assembloids. **(A)** The images show control and ARX^PAE^ assembloids at 90 DIV immunostained against GFP and CXCR4; and stained with DAPI. **(B)** The images show control and ARX^PAE^ GEOs at 120 DIV immunostained against CXCR4; and stained with DAPI. **(C)** Graph shows the percentage of CXCR4^+^ area in control and ARX^PAE^ GEO at 120 DIV. **(D)** Experimental design: COs and GEOs were labeled and fused as described in Fig.4. After a 15-hour baseline recording at 90 DIV, control and ARX^PAE^ assembloids were exposed to AMD3100, a CXCR4 inhibitor, and interneuron migration was evaluated for an additional 15 hours. **(E)** The graph shows the final distance of DLX1/2-GFP^+^ cells in control and ARX^PAE^ assembloids before and after AMD3100 exposure. The results are the mean ± SEM from N= 63-66 neurons from 23-24 organoids from 2 clones x 3 lines per condition. Two-tailed Student’s *t*-test, * = p < 0.05, ** = p < 0.01, *** = p < 0.001, ns = not significant. DIV = days *in vitro*, COs = cortical organoids, GEOs = ganglionic eminence organoids.

### PAE affects cortical development

The expression of ARX in progenitor cells in COs at 30 DIV and the increased expression of *ARX* in ARX^PAE^ COs prompted us to next investigate how PAE affects cortical development. We first analyzed the size of control and ARX^PAE^ COs over time. We found that ARX^PAE^ COs were larger than control COs at 14 and 21 DIV, but no difference was found at 60 DIV (14DIV: P= 0.0005, 21DIV: P= 0.0226, 60DIV: P= 0.3557; **Fig. S20**). The larger size of both ARX^PAE^ COs and GEOs might be compatible with macrocephaly in patients. Interestingly, one of the patients used in the study presented a head circumference of 61 cm at 18 years old, which is at the 99.99^th^ percentile (Z score of 4.07). To explore whether the increased size was due to an increase in proliferation, we labeled dividing cells against KI67 in sections from control and ARX^PAE^ COs at 30 DIV by IHC (*52*). We found a trend towards more KI67^+^ cells in ARX^PAE^ COs (P=0.064, d=0.782, n=12-13; **Fig. S21A-B**). Then, to obtain a more comprehensive profile of the cell composition within the control and ARX^PAE^ COs, we performed cell-type proportion analysis using the scRNA-seq dataset and observed an increase in the percentage of RGCs in ARX^PAE^ COs (**Fig. 6A**). To further investigate the effect of PAE on progenitor cells, we subclustered the RGCs and obtained 9 subclusters (**Fig. 6B**, and **Fig. S22** shows heatmaps displaying the expression of the specific set of genes by each subcluster). *ARX* was mostly expressed in subcluster 2, and some cells in subclusters 1 and 3; and its expression was increased in ARX^PAE^ COs (**Fig. S23**), consistent with the overall increase of ARX expression in ARX^PAE^ COs at 30 DIV described above (see **Fig. 2**). *NESTIN* and *HES5* were highly expressed in all 9 subclusters (**Fig. 6B** and **Fig. S23**), and their higher expression was observed in ARX^PAE^ COs; however, we found *HES1* and *PAX6* were decreased. PAX6, HES1 and HES5 (hes family bHLH transcription factor 1 and 5) are well-known markers of progenitor cells (*42, 64, 65*). Similar reduction of PAX6^+^ cells in ARX^PAE^ COs was found by IHC (P=0.0089, d=0.955, n=17; **Fig. 6C-D**). Moreover, RNAscope analyses confirmed a reduction of cellular *HES1* expression and increase in *HES5* in ARX^PAE^ COs (*HES1*: P=0.0415, d=-1.054, n=11-15; *HES5*; P=0.0465, d=0.707, n=9-15; **Fig. 6C-D**). However, we did not find any significant differences in other progenitor cell markers, such as SOX2 and VIMENTIN, in COs at 30 DIV (**Fig. S21C**). We then analyzed the expression of *TBR2* by RT-qPCR, a marker of IPs (*66, 67*); and we found a significant increase of *TBR2* expression in ARX^PAE^ COs compared to controls (P=0.0077, n=12-17; **Fig. 6E**). Furthermore, it has been shown *Arx* regulates the expansion of cortical progenitor cells in mice by repressing *Cdkn1c*, which negatively inhibits cell proliferation (*16*). Therefore, we then analyzed the expression of *ARX*, *CDKN1CA*, *CDKN1B* and *CDKN1C* in control and ARX^PAE^ COs at 30 DIV by RT-qPCR (**Fig. 6F**). While *ARX* expression was increased in ARX^PAE^ COs, as also observed by IHC and scRNA-seq, *CDKN1C* levels were decreased (*ARX*: P=0.0008, n=12-18; *CDKN1C*; P=0.0195, n=12-17; **Fig. 6F**). This data is consistent with previous findings in mice, and it could explain the increased number of RGCs and IPs observed in COs derived from patients. Together, these data suggested that *ARX* increase due to PAE produces an expansion of RGCs and IPs in COs at earlier time points by downregulating *CDKN1C*.

**Figure 6.**
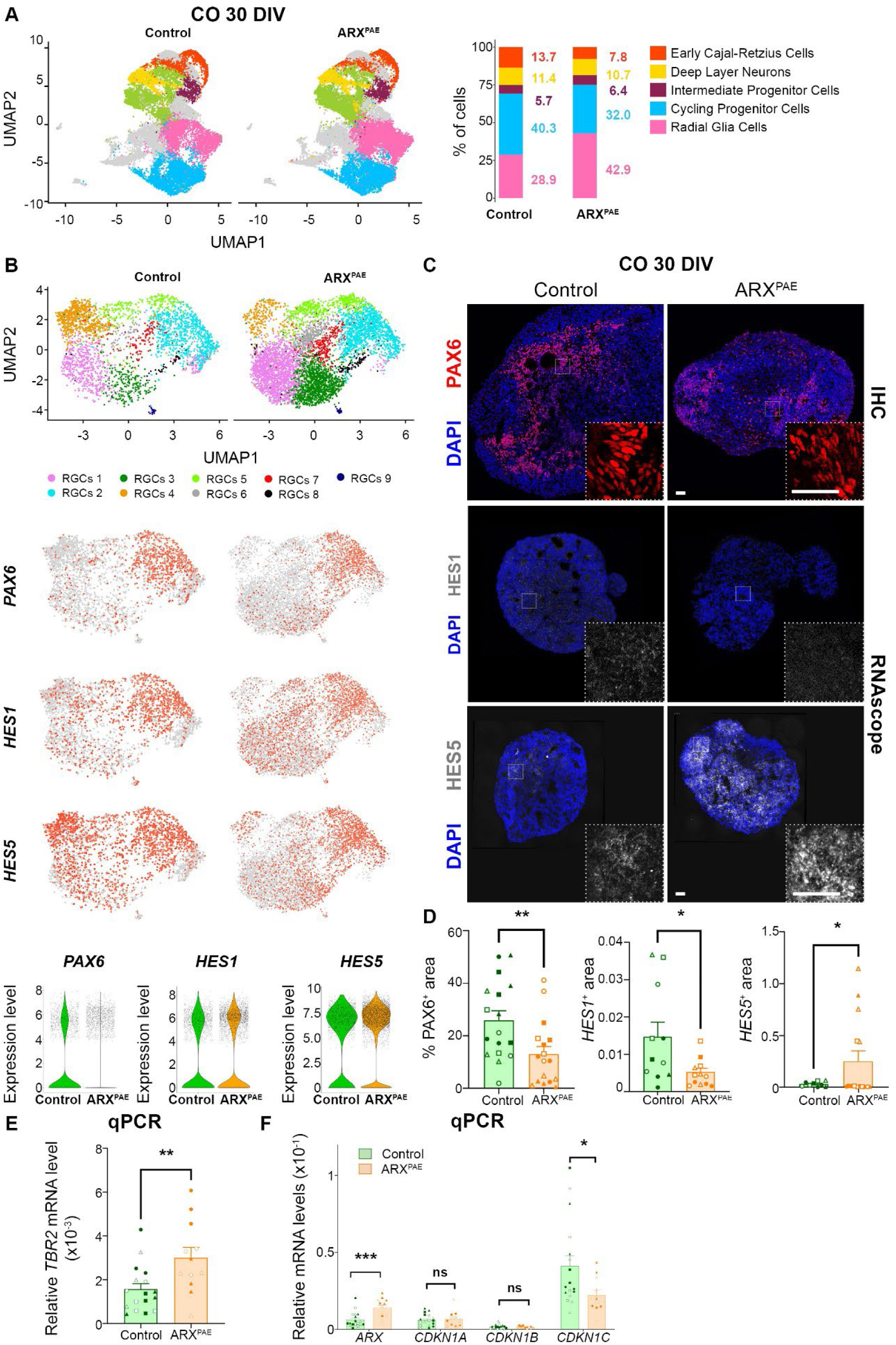
Poly-Alanine expansion affects radial glial cells and intermediate progenitors in cortical organoids. **(A)** UMAP plots from control and ARX^PAE^ COs at 30 DIV. Bar graph shows the percentage of cells in each cluster. **(B)** UMAP plots of subclustering of RGCs from control and ARX^PAE^ COs at 30 DIV. Feature plots showing *PAX6*, *HES1* and *HES5* expression in RGCs. Violin plots show the expression levels of *PAX6*, *HES1* and *HES5*. **(C)** The images show control and ARX^PAE^ COs at 30 DIV immunostained against PAX6, and images of *HES1* and *HES5* expression in control and ARX^PAE^ COs at 30 DIV by RNAscope, and stained with DAPI. **(D)** Graphs show the percentage of PAX6^+^ area in control and ARX^PAE^ COs at 30 DIV, and HES1^+^ and HES5^+^ area divided by DAPI area in control and ARX^PAE^ CO at 30 DIV. **(E)** The graph shows the relative *TBR2* mRNA level in control and ARX^PAE^ COs at 30 DIV measured by RT-qPCR. **(F)** The graphs show the relative *ARX, CDKN1A,* CDKN*1B* and *CDKN1C* mRNA levels in control and ARX^PAE^ COs at 30 DIV measured by RT-qPCR. DIV = days *in vitro*, COs = cortical organoids. Two-tailed Student’s *t*-test, * = p < 0.05, ** = p < 0.01, *** = p < 0.001, ns = not significant. The results are the mean ± SEM from N= 9-17 organoids from 2 clones x 3 lines per condition. Individual points represented data from individual organoid and each symbol represented the data from each clone/line (Line 1: Clone A ● Clone B ○; Line 2: Clone A ◼ Clone B ◻; Line 3: Clone A ▲ Clone B △).

Next, we investigated the long-term consequences of PAE in the cortex by analyzing COs at later time points. Although we did not see any major difference in the cell-type proportion in ARX^PAE^ COs compare to controls at 120 DIV by scRNA-seq, we observed a trend towards a decrease in deep-layer neurons (**Fig. 7A**). The initial expansion of RGCs and IPs and the decrease of deep-layer neurons at later time points prompted us to explore these cell types over time. While we did not detect any significant differences in PAX6^+^ cells or KI67^+^ cells in ARX^PAE^ COs compared to controls at 60 DIV (**Fig. S24A**), we found a significant decrease in PAX6^+^ cells in 90 DIV ARX^PAE^ COs (P=0.0081, d=-1, n=9-15; **Fig. 7B-C**) and a trend towards fewer PAX6^+^ cells at 120 DIV (P=0.2502, d=-0.472, n=12-13; **Fig. 7D-E**). We observed no difference in the percentage of VIM^+^ or HOPX^+^ cells (**Fig. S24C**), but we found a significant reduction in FAM107A^+^ oRGCs (P=0.0086, d=-1.052, n=12-15; **Fig. 7D-E**). Moreover, we did not observe any difference in CTIP2^+^ or SATB2^+^ neurons at 60 or 90 DIV (**Fig. S24A-B**), but we observed a significant decrease of NEUROD1^+^ cells and CTIP2^+^ deep-layer neurons in ARX^PAE^ COs at 120 DIV and a trend towards fewer SATB2^+^ upper-layer neurons (NEUROD1: P=0.0426, d=-0.772, n=16; CTIP2: P=0.0294, d=-0.828, n=15-16; SATB2: P=0.0829, d=-0.701, n=14; **Fig. 7D-E**). Similar results were found by scRNA-seq (**Fig. S25**, plots per condition, and **Fig. S26**, plots per line). To rule out whether the reduction in cortical neurons observed in ARX^PAE^ COs was due to increased cell death, we then analyzed cleaved caspase 3 (AC3) but we did not observe any significant difference in ARX^PAE^ COs compared to controls at any of the time points analyzed, except for a slight increase at 30 DIV (**Fig. S27**). Taken together, our data show that expansion in the second poly-alanine track leads to an increase of *ARX* in COs, which promotes the expansion of RGCs and IPs. These progenitor cells may undergo apoptosis resulting in an overall reduction in cortical neurons at later time points.

**Figure 7.**
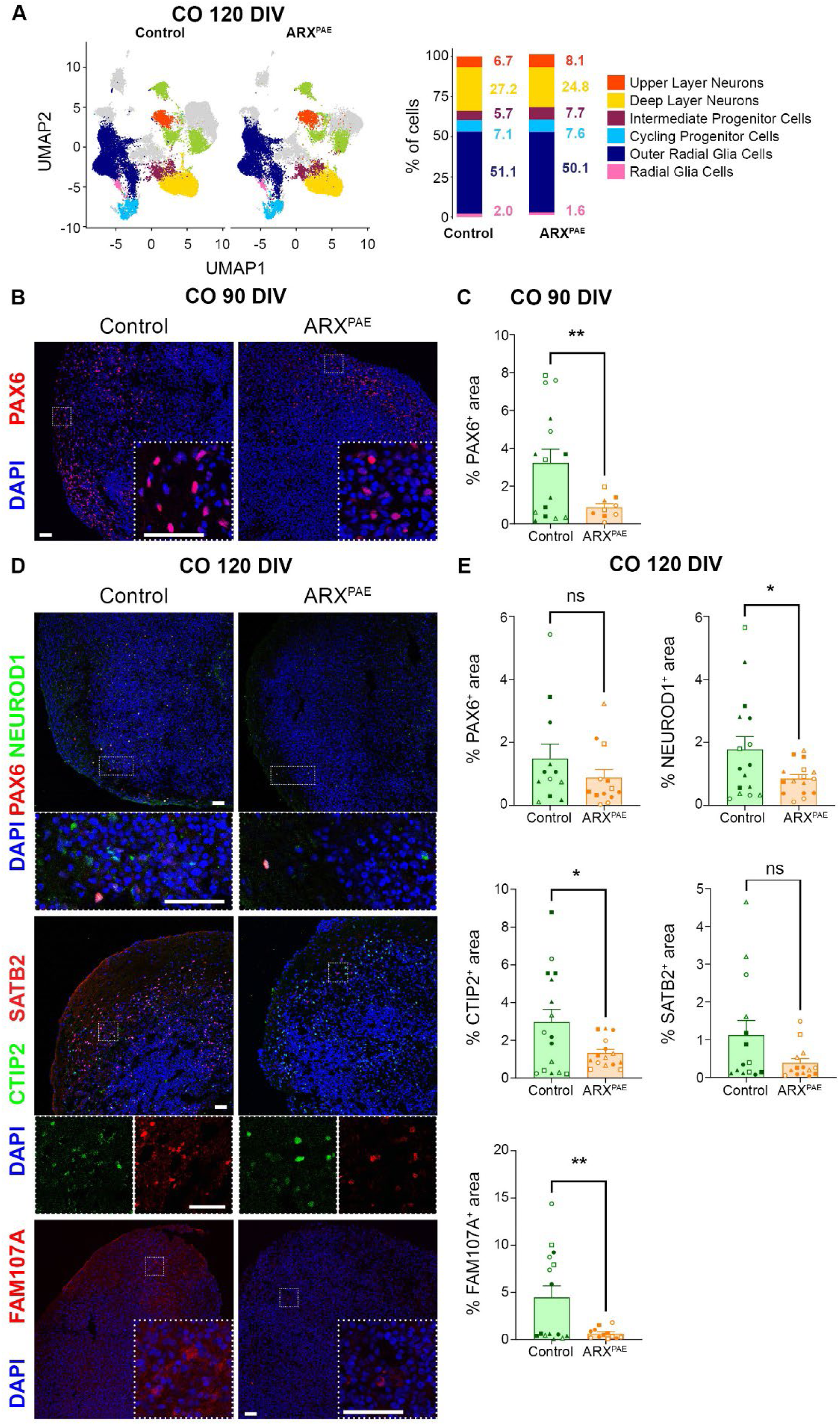
Poly-Alanine expansion compromise cortical neuronal differentiation in cortical organoids. **(A)** UMAP plots from control and ARX^PAE^ COs at 120 DIV. Bar graph shows the percentage of cells in each cluster. **(B)** The images show control and ARX^PAE^ COs at 90 DIV immunostained against PAX6, and stained with DAPI. **(C)** Graphs show the percentage of PAX6^+^ area in control and ARX^PAE^ COs at 90 DIV. **(D)** The images show control and ARX^PAE^ COs at 120 DIV immunostained against PAX6, NEUROD1, CTIP2, SATB2 and FAM107A; and stained with DAPI. **(E)** Graphs show the percentage of PAX6^+^, NEUROD1^+^, CTIP2^+^, SATB2^+^ and FAM107A^+^ area in control and ARX^PAE^ COs at 120 DIV. Scale bar = 50 µm. DIV = days *in vitro*, COs = cortical organoids. Two-tailed Student’s *t*-test, * = p < 0.05, ** = p < 0.01, ns = not significant. The results are the mean ± SEM from N= 12-17 organoids from 2 clones x 3 lines per condition. Individual points represented data from individual organoid and each symbol represented the data from each clone/line (Line 1: Clone A ● Clone B ○; Line 2: Clone A ◼ Clone B ◻; Line 3: Clone ▲ Clone B △).

### Impact of PAE on neuronal activity

We next aimed to investigate whether the cellular and molecular alterations observed in COs and GEOs derived from patients with PAE mutations in *ARX* gene impact neural network connectivity and functionality post-fusion. COs and GEOs were fused at 60 DIV and MEA recordings were performed at 120 DIV, an early developmental time point when we have a decrease of cortical neurons and defects in interneuron migration (**Fig. 8A**). We observed a larger number of electrodes active in the 30-min recording and an increase of the spike number in ARX^PAE^ assembloids compared to control (**Fig. 8B-C**). Some assembloids also showed synchronous burst activity in both conditions (1 out of 3 assembloids in controls and 2 out of 3 in ARX^PAE^, data not shown). We then investigated the role of the GABAergic system in the network at the receptor level, by exposing each assembloid to GABA and picrotoxin, a GABA_A_ receptor blocker. Surprisingly, GABA or picrotoxin had no effect on spiking and bursting activity in control and ARX^PAE^ assembloids at 120 DIV, indicating that the GABA_A_ polarity is not completely switched from excitatory to inhibitory to recapitulate earlier stages of brain development. Our data suggest that ARX^PAE^ organoids are hyperactive compared to controls.

**Figure 8.**
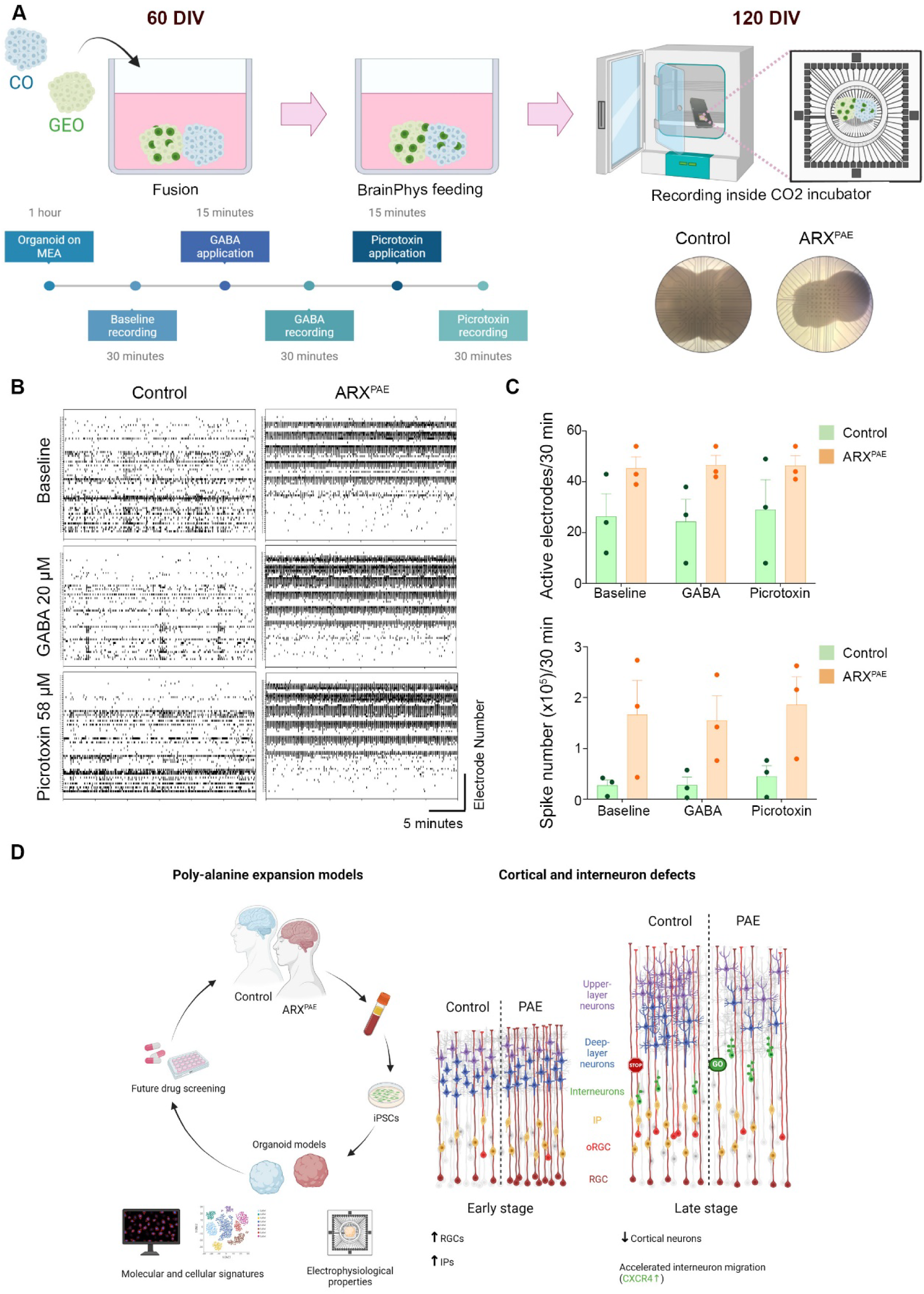
Poly-Alanine expansion promote hyperactivity in ARX^PAE^ assembloids. **(A)** Schematic illustration of MEA experiment. All recordings were performed inside a humid, CO_2_ incubator while BrainPhys media is bath applied to the assembloids. Light microscopy images show assembloids embedded on 3D-MEA covering the entire electrode pitch within each recording. Sequential recordings were carried out on each individual organoid, starting from one hour rest on the MEA, baseline recording, and continuing through GABA and picrotoxin application. **(B)** Representative 5 min-raster plots from control and ARX^PAE^ assembloids under baseline, GABA and picrotoxin conditions at 120 DIV. The vertical axis represents electrode numbers and the horizontal axis shows 5 minutes of a 30-minute recording. **(C)** Graphs show the number of active electrodes and the number of spikes in control and ARX^PAE^ COs at 90 DIV in 30-minute recording. The results are the mean ± SEM from N= 3 organoids from 2 lines per condition. **(D)** Development of an *in vitro* model using neural organoids to study PAE-related epilepsies and other human diseases. PAE in the second tract of *ARX* promotes interneuron migration due to an increased CXCR4 expression. Moreover, PAE produces an initial increase of RGCs and IPs and decrease of cortical neurons at later time points. Both defects promote hyperactivity. DIV = days *in vitro*, COs = cortical organoids, GEOs = ganglionic eminence organoids, IPs = intermediate progenitor cells and RGCs = radial glial cells.

Altogether, our results highlight the usefulness of neural organoids in studying epilepsy and other neurodevelopmental disorders and demonstrated PAE in the second tract of ARX have context-dependent consequences in cortical cells and interneurons (**Fig. 8D**). While it produces expansion of RGCs and IPs and loss of cortical neurons in the cortex, it affects interneuron differentiation and migration from the subpallium. These defects influence neuronal activity early on during development and may underlie the seizure activity observed in patients with PAE mutation in ARX.

## DISCUSSION

ARX is a transcription factor critical for brain development (*14, 16, 17*). Although mutations in *ARX* have been described in patients with a broad spectrum of neurological disorders, the specific consequences of each mutation are not completely understood. Moreover, many patients suffer from seizures refractory to treatment. To gain a more complete understanding of the pathobiology underlying these seizures, we studied how PAE in the second tract of *ARX*, one of the more common mutations found in patients with *ARX* mutations, impacts brain development using hiPSCs derived from patients with this mutation. hiPSCs were differentiated into CO and GEO, *in vitro* models of human cortical and GABAergic development respectively, and we found that ARX affects both brain areas through distinct mechanisms. PAE promotes an increase of RGCs and IPs in the cortex (dorsal progenitor cells, CO), but it stimulates interneuron migration from subpallium regions (GEO). We identified consistent cellular and molecular changes from multiple hiPSC lines derived from three healthy controls and three patients, illustrating reproducibility across experiments, batches, and cell lines in organoid models. This suggests that the observed differences are unlikely to be solely attributed to genetic background variation between lines. However, the lack of isogenic controls, stemming from the GC-rich and the repetitive nature of the region (PAE), prevent us from completely excluding this possibility. Taken together, our study provides crucial insights into mechanisms underlying PAE mutations and offers a potential human cellular model to develop new therapeutic interventions.

*ARX* is expressed in both the developing and adult brain (*14, 15*). During development, *Arx* is expressed in dorsal progenitor cells, while in adults, its expression is restricted to interneurons originating in the GE (*14, 16, 17*). In this study, we found ARX expression in both proliferating progenitor cells and differentiated cortical neurons in COs. Consistent with murine studies, ARX was also expressed in ventral progenitor cells and interneurons in GEOs, with higher expression observed compared to dorsal tissue. Moreover, our analysis of publicly available brain transcriptome data revealed the expression of *ARX* in both human embryonic and postnatal cortical and subpallial tissues, demonstrating its widespread expression during brain development in both dorsal and ventral compartments. Our data also demonstrate that PAE affects *ARX* expression in a context-dependent manner, for both cell type and developmental timing. For instance, we observed an increase in the number of ARX^+^ cells in COs, while its expression was decreased in GEOs at an early timepoint. This contrasts with findings from mouse studies, where a reduction in ARX protein was observed in embryos with PAE mutations in the first or second tract of ARX (*19, 68*). Similarly, adult mice with PAE in the first tract exhibited a reduction in the number of ARX^+^ cells in the cortex, hippocampus, and striatum (*28*). These differences suggest that PAE might have distinct effects on *ARX* expression in mice compared to humans. Additionally, studies have shown the presence of nuclear aggregates *in vitro* when *ARX* containing PAE is overexpressed (*58, 59*), but not *in vivo* (*19, 28, 68*), suggesting that under physiological conditions, ARX may not form nuclear inclusions. Moreover, cytoplasmic expression of Arx has also been observed in mice with PAE (*28, 60*). Interestingly, we did not detect any nuclear inclusions or cytoplasmic expression of ARX in either COs or GEOs, indicating that a different mechanism might be involved in humans. Further analysis of *ARX* expression in various brain regions over time is necessary to elucidate how PAE impact *ARX* expression in human with the ultimate goal of developing novel strategies to treat these and related neurodevelopmental disorders.

A reduction of GABAergic interneurons has been observed in the cortex of both KO mice and PAE models (*19–22, 25, 28*), as well as in zebrafish larvae (*69*). Patients with XLAG similarly exhibit reduced numbers of interneurons (*70*). Given that interneurons originate in the GE and migrate tangentially to the cortex, the observed reduction in cortical interneurons suggests that ARX plays a crucial role in neuronal migration (*54, 61*). Indeed, studies have shown the accumulation of cells in the GE following inactivation or overexpression of *Arx* (*15, 17, 25*). Moreover, aberrant migration of *Arx* KO cells has been demonstrated in forebrain slice culture experiments (*18*). Notably, Lee and colleagues found that while PAE in the first tract inhibits neuronal migration, PAE in the second tract does not affect neuronal migration (*25*). Our results show that human PAE in the second tract initially compromises GABAergic differentiation and subsequently enhances neuronal migration to the cortex. Through time-lapse experiments, we observed that DLX1/2-GFP^+^ interneurons with PAE exhibited accelerated migration, advancing further and faster into cortical tissues in a cell-autonomous manner, as shown by inter-individual assembloid experiments. However, despite this accelerated migration, we did not observe an increased number of interneurons in the cortex. This discrepancy raises the possibility that interneurons with PAE continue to migrate due to failure in encountering their final target or the absence of crucial stop signals. Alternatively, these migrating interneurons may fail to integrate into the existing circuitry, potentially leading to cell death. Further experiments using retrograde rabies viruses tracing are needed to provide insight into how PAE expansion in ARX affects interneuron connectivity after migrating to the cortex. Notably, we did observe an increase in the expression of CXCR4 in GEOs. The CXCR4/CXCL12 signaling axis plays a critical role in interneuron migration (*56, 57, 63, 71*). Previous studies have shown that when a CXCR4 blocker is added to human assembloids similar to those presented in our study, the migration velocity of interneurons is decreased (*32*). Importantly, CXCR4 is a direct target of ARX, with ARX promoting its expression (*22, 72*). Therefore, it is conceivable that PAE enhances the binding capacity of ARX to CXCR4, thereby promoting its expression and accelerating neuronal migration. Indeed, AMD3100, a CXCR4 inhibitor, was sufficient to reduce migration in ARX^PAE^ assembloids to control levels, while it did not affect interneuron migration in controls. These findings suggest that, although other mechanisms may contribute to interneuron migration, PAE in ARX predominantly promotes migration through CXCR4 signaling.

RGCs are the neural stem cells in the cortex that arise from the neuroepithelium and eventually differentiate into neurons (*73, 74*). In our study we found that PAE in *ARX* leads to an increase in RGCs and IPs along with an increase in *ARX* expression in COs at 30 DIV. We also observed a mixed expression of canonical neural progenitor markers in RGCs at 30 DIV COs with PAE. For instance, although the expression of genes such as *NESTIN* and *HES5* (neural progenitor markers) was upregulated, there was a downregulation of other canonical genes such as *PAX6* and *HES1*. Our findings demonstrated the heterogeneity in RGCs and partially align with previous studies conducted in mice, which demonstrated that *Arx* KO mice exhibit thinner cortices and reduced numbers of BrdU^+^ cells (*15*). Moreover, abrogation of *Arx* from pallial progenitor cells decreases the number of BrdU^+^, Pax6^+^ and Tbr2^+^ cells, but increases Tuj1^+^ cells (*16*). Similarly, downregulation of Arx in cortical progenitor cells produces a premature cell cycle exit, while overexpression of Arx extended the cell-cycle length of progenitor cells in mouse embryos (*17*). In addition, *Arx* KO mice exhibited an overexpression of *Cdkn1c* (*16*), and we found a reduction of *CDKN1C* levels in ARX^PAE^ COs at 30 DIV and an increase in *ARX* expression. This data suggests that *CDKN1C* might be a direct target of ARX and might be responsible of the initial RGCs and IPs expansion observed in ARX^PAE^ COs. Additional experiments are needed to fully understand how PAE affects the interaction of ARX with other direct targets and with its binding partners to regulate gene expression during cortical development.

We also found a reduction of cortical neurons at later timepoints, possibly due to the initial expansion of RGCs and IPs in ARX^PAE^ COs and depletion of the progenitor pool. The initial expansion of RGCs and IPs could explain the increased size observed in ARX^PAE^ COs at 14 and 21 DIV, as well as the macrocephaly observed in some patients (*75*). However, it is important to note that not all of these progenitor cells will differentiate into neurons; some may undergo apoptosis, as suggested by the trend toward increased AC3 staining in ARX^PAE^ COs at 30 DIV. In addition, PAE in ARX may lead to a reduction of RGCs and oRGCs at later time points, as evidenced by the decreased expression of PAX6 and FAM107A in ARX^PAE^ COs at 90 and 120 DIV, respectively. Taken together, these findings would be consistent with the decrease in cortical neurons at later time points. While neuronal cortical defects specifically related to PAE have not been extensively studied in mouse models or patients, defects in the organization of pyramidal neurons have been documented in conditions such as X-linked lissencephaly with abnormal genitalia (XLAG) (*70*) and in *Arx* KO mice (*15*). Moreover, reductions in upper-layer cortical neurons have been shown in postnatal *Arx* cKO mice (*16*). These findings collectively suggest that both KO mutations and PAE in ARX result in a reduction in cortical neurons.

Patients with PAE in *ARX* frequently develop epilepsy, among other neurological manifestations (*76, 77*). It has been proposed that a defect in interneuron numbers or function is the primary cause of seizure development in mouse models of PAE (*20, 21*), while cortical defects contribute to other behavioral phenotypes (*24*). Human organoid models display complex neuronal oscillations resembling human brain activity, offering an opportunity to model cortical function in various disease states (*38, 78*). Thus, to investigate the impact of PAE mutations on neuronal function at early stages of development, particularly within a network-wide context, we used MEA recordings. Our results revealed heightened network activity in ARX^PAE^ assembloids, as evidenced by increased spike number compared to control assembloids at 120 DIV. ARX^PAE^ assembloids showed broader network activity as more electrodes were active through each session of recording, in every condition. Our observation suggests that synchronous bursting (where 25 electrodes are synchronously bursting) is more frequent in ARX ^PAE^ organoids, which is a common feature observed in animal models and patients with epilepsy (*79–81*). Network activity in controls and ARX^PAE^ assembloids had subtle responsiveness to GABA at 120 DIV. This data suggests that GABA_A_ polarity was not completely switched from excitatory to inhibitory at this time point. Although our results showed hyperactivity in ARX^PAE^ assembloids, longer MEA recording, as well as analysis of older assembloids would be useful to fully evaluate possible epileptic-like behavior in a more mature network. Furthermore, as ARX is also expressed in other brain regions, including the hippocampus, striatum and thalamus (*14, 15, 82*), it is possible that organoid models lacking those structures cannot fully recapitulate the epilepsy of patients with PAE. In fact, Beguin and colleagues proposed the hippocampus as the starting point of epileptic activity in *Arx* mouse models (*83*). Developing more complex organoid models would be critical to study diseases affecting several brain regions and would permit researchers to disentangle the contribution of each region to the phenotypes.

In summary, our results demonstrate that ARX is a critical player during brain development, and also highlight the usefulness of neural organoids in studying epilepsy and other human neurodevelopmental diseases (**Fig. 8D**). We present evidence that PAE in the second tract of *ARX* promotes interneuron migration which is associated with altered CXCR4 expression. Furthermore, PAE produces an initial increase of RGCs and IPs followed by a decrease in cortical neurons at later timepoints. Both defects in cortical and GABAergic neurons contribute to hyperactive reminiscent of *Arx* mouse models and patients. This study provides new information for understanding the pathological mechanisms under PAE mutation and demonstrates a unique human cellular model to develop potential treatments.

## MATERIALS AND METHODS

### Study design

The goal of this study was to determine the role of ARX PAE mutations in human brain development and epilepsy. PAE mutations is one of the most prevalent mutations in *ARX* and are associated with ID, ASD, and epilepsy. Moreover, many epileptic patients show resistance to current treatments, emphasizing the need to develop new model for drug screening. We used human neural organoid models derived from three male patients with a PAE in the second tract of *ARX*, along with three healthy male controls. We focused on male patients because *ARX* is located on the X Chromosome. Two types of human regionalized neural organoids from hiPSCs: COs and GEOs were generated to mimic cortical neuron and interneuron development. Organoids were analyzed by scRNA-seq, IHC, time-lapse imaging and MEA recording at different time points. All quantifications were performed in a blinded manner. A total of six lines per condition (2 clones per subject) were used for all experiments or otherwise indicated in the figure legends. Sample size is indicated in the figure legends and it was determined by the investigators based on previous work. Outliers were removed from each dataset using GraphPad Prism software.

### Pluripotent Stem Cell Generation and Maintenance

A total of six human induced pluripotent stem cell (hiPSC) lines (**Table S1**) were generated from blood collected from three control subjects and three ARX^PAE^ patients using an episomal reprogramming kit (Invitrogen) or CytoTune-iPS Sendai Reprogramming kit (Thermofisher) and two clones per line were used. Approval for the study was obtained from the University of Texas at San Antonio IRB panel and informed consent was obtained from all subjects. Pluripotency markers were confirmed by immunocytochemistry and flow cytometry; and normal karyotype and absent of mycoplasma was maintained throughout the study in all hiPSC lines (**Fig. S8**). We also confirmed no integration of reprogramming genes. hiPSCs were maintained in mTeSRTM1 medium (Cat. No. 05851, Stemcell Technologies) on six-well plates (Cat. No. 3506, Corning) coated with growth factor-reduced Matrigel (Cat. No. 356230, BD Biosciences) and adding ROCK inhibitor (final concentration 10 µM, Cat. No. S-1049, Selleck Chemicals). The cells were maintained with daily medium change without ROCK inhibitor until they reached 70% confluency. Then, they were detached using versene solution (Cat. No. 15040-066, Thermo Fisher Scientific) and plated 1:10 - 1:20.

### Genotyping

Poly-alanine expansion on the second tract of *ARX* were confirmed by sequencing. Genomic DNA was extracted from hiPSCs using DNeasy® Blood & Tissue Kit (QIAprep #69504) following manufacturer’s instructions. OneTaq® Hot Start DNA Polymerase (New England BioLab Cat. No. M0481) was used to amplify the second alanine tract of *ARX* using forward (GGGGGCCGCCTCCTTCAG) and reverse (CCTGGTGAAGACGTCCGGGTAGTG) primers. PCR conditions were as follow, 94 °C for 4 minutes; 94 °C 30 sec, 60 °C 30 sec, 68 °C 1 minute for 35 cycles; 68 °C 5 minutes. PCR products (798 bp) were visualized on an agarose gel and sent for Sanger sequencing.

### Flow cytometry analysis

To analyze pluripotent stem cell markers, hiPSCs were dissociated using accutase (Sigma). 10^6^ cells were resuspended in D-PBS/ 1% fetal bovine serum (FBS). After two washes with D-PBS/1% FBS, cells were incubated for 20 minutes on ice with primary antibodies (anti-SSEA-4 PE antibody, 1:20, Stemgent #09-0003; anti-TRA1-60 PE, 1:5, BD# 560193). Then, they were washed three times and resuspended in 300 µl D-PBS/1% FBS and analyzed by flow cytometer (BD FACSAria) with 10,000 events per determination. Post-acquisition analysis was done using Flowjo software (Tree Star Inc) and the percentage of cells expressing each marker was determined.

### Organoid Culture

Human cortical organoids (CO) and human ganglionic eminence organoids (GEO) were generated using the method described by Pasca and colleagues with slight modifications (*32, 35*). Briefly, hiPSCs were dissociated using accutase (Sigma) and 9,000 cells were resuspended in 150 µl neural induction medium (DMEM/F12, 20% knock-out serum replacement, Glutamax, MEM-NEAA, 0.1 mM 2-mercaptoethanol, penicillin/streptomycin) containing Y27632 (20 µM). Media was changed daily. On days 1-5, the SMAD inhibitors, dorsomorphin (5 µM, Sigma) and SB-431542 (10 µM, Tocris) were added. On days 6-24, media was replaced with neural medium (Neurobasal A, B-27 without Vitamin A, Glutamax, penicillin/streptomycin) containing the growth factors, bFGF (20 ng/ml, Peprotech) and EGF (20 ng/ml, Peprotech). On day 25, organoids were transferred to neural medium containing BDNF (20 ng/ml, Peprotech) and NT-3 (20 ng/ml, Peprotech) until day 42. GEOs also received the Wnt pathway inhibitor, IWP-2 (5 µM, Selleckchem) on days 4-22 and the Smo pathway activator, SAG (100 nM, Selleckchem) on days 12-22 (**Fig. 1A**). After day 25, organoids were maintained on an orbital shaker to promote oxygenation. Each organoid was maintained individually in wells of 96-well plate until day 24 and transferred to 24-well plate after that.

### Interneuron Migration

On day 50, GEOs were infected with lentivirus hDlx1/2b:GFP (gift from Dr. John Rubenstein) and CO with an AAV-DJ1-hSYN1:mCherry (Stanford Gene Vector and Virus Core at Stanford University School of Medicine). On day 60, one GEO and one CO were transferred to a well of a 24-well plate and placed in close contact, containing neural media and incubated without disruption for 3-4 days. On day 90, assembloids were transferred to glass-bottom plates (Corning) and the migration of hDLX1/2b:GFP interneurons into cortical tissue was recorded under environmentally controlled conditions (37 °C, 5% CO2). Time-lapse images were taken every 20 min for 15 hours using a confocal Leica Microscope with a motorized stage (TCS SPE8) (**Fig. 4A**). In each field, we took images every 3 µm up to 150-200 µm thickness. For analysis, neurons were selected randomly from hDLX1/2b:GFP interneurons presented in the cortical side and only neurons that did not move out of the field during the recording time were analyzed. Organoids with major drift were discarded. In some experiments, after 15 hours time-lapse imaging, assembloids were exposed to AMD3100 (CXCR4 inhibitor, 100 nM; TOCRIS #3299) and interneuron migration was evaluated for another 15 hours. Some assembloids were fixed, sliced, and imaged. Image J was used to determine the distance and the average velocity of the hDLX1/2b:GFP cells when mobile. Videos were processed using Leica software.

### Immunohistochemistry

Organoids were fixed in 4% paraformaldehyde overnight at 4 °C and incubated in 30% sucrose for 48 hours at 4 °C. Then, organoids were embedded in OCT compound and frozen. A cryostat was used to cut 14-µm sections, which were mounted on slides. For immunohistochemistry, slides were incubated in blocking solution (0.3% Triton X-100, 3% normal donkey serum in TBS) for 1 hour at room temperature, and primary antibody (**Table S2**) overnight at 4 °C in a humidified chamber. After washing three times, slides were incubated in secondary antibody (Jackson Immunoresearch, 1:500) for 2 hours at room temperature. Slides were washed, 4’,6-diamidino-2-phenylindole (DAPI; Sigma, Cat. No. D9542) was added to label nuclei and then coverslipped using polyvinyl Alcohol solution (PVA; Sigma, Cat. No. BP168-122). Four sections per organoids were imaged using a Leica Microscope (Spe8Info) and fluorescence intensity area was analyzed using Image J software. Same acquisition conditions were used for each marker in controls and ARX^PAE^ organoids. A threshold for each marker was applied to discard non-specific background fluorescence. As some of the markers were not distributed homogenously, all the quantifications were performed in the whole section. For nuclear markers, data were expressed as the percentage of area labelled for each marker divided by DAPI area. For cytoplasmic markers, data were expressed as the percentage of area labelled for each marker divided by total area of the section. Each data point represents the mean of 4 sections. For most of our analysis 3 organoids per clone per line per condition (total of 18 organoids per condition) were used. All analyses were conducted blindly.

### RNAscope

RNAscope was performed on cryostat sections following manufacturer’s instructions with antigen retrieval (ACDBio). Human probes were custom-made by ACDBio for *HES1* and *HES5* mRNA. Four sections per organoids were imaged using a Leica Microscope (Spe8Info) at 40x oil immersion objective. Quantification was performed using CellProfiler. Data are shown as the ratio of *HES1* or *HES5* mRNA to DAPI expression.

### Dissociation of organoids and single cell RNA sequencing (scRNA-seq)

For single-cell dissociation and RNA seq, COs and GEOs were dissociated using Worthington papain dissociation kit (Cat. No. LK003150) per the instruction manual. Each developmental timepoint (30 DIV and 120 DIV) and each subtype of organoid (cortical and ganglionic eminence) was processed separately. A total of 3 lines, and one clone per line, were used for each condition. Briefly, 3-4 organoids for each timepoint and subtype were collected in a small petri dish with 5 ml of prewarmed Worthington papain solution and 250 µl of Worthington DNase solution. The organoids were chopped into small pieces using sterile blades and the dishes were placed on a digital rocker in a cell culture incubator for 60 minutes at 37 °C. Digested tissue was collected in a 15-ml tube and 5 ml of room temperature Earle’s Balanced Salt Solution (EBSS) was added. The mixture was then triturated by gently pipetting up and down 10 times with 10-ml plastic pipette followed by 1-ml and 200-µl pipettes. Any undissociated tissue was allowed to settle at the bottom of the tube and the cloudy suspension was carefully aspirated in a clean 15-ml tube. Next, 3.15 ml of inhibitor solution (2.7 ml of EBSS, 300 µl of reconstituted Worthington albumin-ovomucoid solution and 150 µl of DNase solution) was carefully layered over the cloudy suspension and tubes were centrifuged at 300 g for 5 minutes at room temperature. The supernatant was carefully discarded, and cell pellet was gently suspended in 500 µl of cold Neurobasal-A media. Cell viability was determined using trypan blue.

10x Genomics single cell libraries were generated by the University of Texas at San Antonio (UTSA) Genomics Core. Cell suspensions were loaded into 10x Genomics microfluidics chips from Chip B Single Cell kit (Cat. No. 1000153) and onto the 10x Chromium controller to capture single cells in Gel Beads-in-emulsion (GEMs). 7,000 cells/sample were targeted for capture in GEMs preceding library prep following manufacturer’s recommendations for 10x Genomics 3’ Gene Expression Reagent kit (Cat. No. 1000269) with v3 chemistry. Sequencing was performed on either NextSeq 500 at University of Texas Health Science Center (UTHSC) at San Antonio or NovaSeq 6000 at North Texas Genome Center (NTGC) to generate 50,000 reads/cell in each sample.

### scRNA-seq data analysis

Cell Ranger from 10x Genomics and Seurat were used for all scRNA-seq data analysis as described below. Base calls for each sample were converted into FASTQ reads using cellranger mkfastq. FASTQ reads were aligned to the GRCh38 -3.0.0 human reference genome assembly and cell-by-gene count matrices were generated using cellranger count function with default parameters except ‘--expect-cells’ set to 8,000. Libraries for Control 1 and ARX^PAE^ 1 at COs 120 DIV were re-sequenced to increase the read depths and files were aggregated using cell ranger. All further analysis was performed using Seurat v.3.2 in in R (v.3.6.0) or Seurat v.4.0 in R (v.4.0.3) (*84, 85*). Briefly, for each timepoint (30 DIV and 120 DIV) and organoid subtype (cortical and ganglionic eminence), SeuratObject was initialized for controls (Control 1, Control 2, Control 3) and ARX^PAE^ (ARX^PAE^ 1, ARX^PAE^ 2, ARX^PAE^ 3) separately using filtered feature matrices. Genes that were expressed in ≥10 cells (min.cells=10) and cells with at least 500 detected genes (min.features = 500) were used as filtering parameters for creating each SeuratObject. Next, all six SeuratObjects were merged together using merge function from Seurat to create a single SeuratObject for all further analysis. Mitochondrial and ribosomal genes were calculated and cells with nFeature_RNA>500 and nFeature_RNA<6000 were retained after further filtering and quality control. Unique molecular identifier (UMI) counts were normalized for each cell by the total expression multiplied by 10^6^ and log-transformed. Variable genes were identified using mean.var.plot method in Seurat and ScaleData function was used to regress out all the unwanted sources of variations including mitochondrial and ribosomal genes. Principal Component Analysis (PCA) was run on scaled data and the top principal components (PCs) were chosen based on Seurat’s ElbowPlots (19 PCs were chosen for each merged SeuratObject). RunHarmony function from harmony package (v.0.1.0) was used for batch correction and cells were clustered in PCA space using Seurat’s FindNeighbors function with reduction=‘harmony’ and dims=1:30 followed by FindClusters with resolution=0.5. Uniform manifold approximation and projection (UMAP) in Seurat was finally used as a non-linear dimensionality reduction technique to visualize and further explore the cells in each dataset.

Upregulated genes in each cluster were identified using the VeniceAllMarker tool from Signac package (v.1.3.0) from BioTuring (https://github.com/bioturing/signac). Each cluster was annotated based on the expression of top 20 genes by cross comparation to human fetal tissue and human organoid datasets (*38, 86, 87*). If the top 20 genes were insufficient to define the cluster, it was labeled as “Undefined”. DoHeatmap function from Seurat was used for creating heatmaps representing the expression of genes used to identify cell types. Celltype proportion in each dataset was calculated using glmer function from R package lme4 (v.1.1.27.1) and the script was adapted from Paulsen *et al.* (*88*). For RGCs in 30 DIV cortical organoid dataset, cells were further subclustered. The datasets presented in this study can be found in NIH repositories (accession number GSE215362). The codes used in this study can be found on https://github.com/parulvarma123/ARX_Organoids_scRNAseq.

### RNA isolation, RT-PCR and quantitative PCR

Organoid RNA was extracted using the Qiagen RNeasy Micro Kit according to manufacturer instructions. The concentration and purity of the RNA samples were measured by using Nano-drop (Thermo Fisher Scientific). The extracted RNA (500 ng) was reverse transcribed according to the protocol supplied with SuperScriptIII First-Strand Synthesis System for RT-PCR (Catalog# 18080-051, Invitrogen Life Technologies). Quantitative real-time PCR (qRT-PCR) was carried out using ViiA (Applied Biosystems, Foster City, CA, USA) using PowerUp™ SYBR™ Green Master Mix for qPCR (Catalog# A25741, Applied Biosystems). Reactions were run in triplicate and expression of each gene were normalized to the geometric mean of GAPDH as a housekeeping gene and analyzed using the ΔΔCT method. The primer sequences of each gene are listed in **Table S3**.

### Multi-Electrode Array (MEA) Assay

All electrophysiological recordings were carried out inside a humid (i.e., 95% R.H.) incubator at 37℃ and 5% CO_2_. To minimize stress on the brain organoids during electrophysiological recordings, they were maintained using BrainPhys^TM^ hPSC Neuron kit (05795; STEMCELL Technology) through 2-7 days immediately preceding the recording sessions. For each session, individual organoids were placed on either the 3D (60-3DMEA200/12/80iR-Ti, Multichannel system, Harvard Bioscience) or 2D commercially available Microelectrode Arrays (MEAs) (60MEA200/30iR-Ti, Multichannel systems, Harvard Bioscience). No differences were found between recording using 2DMEA or 3D MEA microelectrodes, as the manufacturer produce all MEAs with similar impedance, therefore the data were combined. To ensure optimal attachment to the MEA surface, 200 µL of BrainPhys^TM^ hPSC Neuron media was also bath applied. By using only 200 µl of media, we prevented the organoids from floating while the brain organoid was fully immersed in nutritious media. Recordings were made on three organoids at 120 DIV (n=4) in each group. All assembloids used in this experiment were either purely ARX ^PAE^ or control, created by the fusion of CO and GEO that carries identical genotypes. Sequential recordings were then performed on each organoid, as it follows: baseline conditions, 20 µM GABA (56-12-2, Sigma-Aldrich), and finally 58 µM picrotoxin (124-87-8, Sigma-Aldrich) (**Fig. 8A**). These chemicals were applied to the culture medium (bath applied), and careful media exchange occurred every 45 minutes to prevent organoid movement on the electrodes. Recording of electrical activity from each of individual organoid on MEA started 10 minutes after the chemical application, allowing their full dilution in the medium and discarding artifacts and transient responses. To prevent media evaporation and optimize recording conditions, each MEA was sealed with fluorinated Teflon thin film (MEA-MEM-set5, Multichannel systems, Harvard Bioscience) that selectively allows gas permeability but blocks water evaporation and microorganism contamination. Each recording session lasted 30 minutes. Each 2D-MEA featured 60 Titanium nitrate (TiN) microelectrodes, with 30 µm of diameter, arranged in a uniform 8 x 8 layout with a 200 µm inter-electrode pitch. Similarly, each 3D-MEA featured 60 TiN microelectrodes with 50 µm diameter and 100 µm pitch, but were organized in 3D with a height of 80 µm. Raw electrical potentials, detected extracellularly from each microelectrode, were amplified using an electronic amplifier (ME2100-Mini, Multichannel systems, Harvard Bioscience), sampled at 25 kHz/channel, and digitized at 16 bits resolution. The recorded extracellular potential traces were stored on disk via software (Experimenter, Multichannel systems, Harvard Bioscience) for subsequent offline analyses. To detect the time of occurrence of putative action potentials (i.e. spike times), a peak detection algorithm with adaptive threshold was employed, so that spontaneous network-wide synchronization of spike times, i.e., network bursts (*89, 90*) could be detected and quantified. The events detected over time at each distinct MEA microelectrode were visualized as a raster plot and further analyzed by conventional spike train analysis (*89, 90*). The degree of functional interaction between pairs of MEA microelectrodes was derived using conventional cross-correlation analysis of spike times(*91*), limited to inter-spike delays less than 500 ms and quantified by 3 ms bins. To account for firing rate modulation, a normalization procedure divided each cross correlogram by the square root of the product of the number of spikes in either of the spike trains. The peak value from the cross correlogram represented the connectivity strength of the electrode pairs. Distributions of these peak values across various experimental conditions were generated to facilitate a comparison of coupling strength. To assess the significance of these peaks, their corresponding inter-spike intervals (ISI) were randomly shuffled to achieve surrogate spike times with the identical distribution of ISI. The peaks were considered significant, if they showed values larger than the mean plus 3 standard deviation of the cross correlogram of the surrogate.

### Human Fetal Brain Tissue Analysis

Fetal brain bulk RNA sequencing dataset was downloaded from BrainSpan: Atlas of the Developing Human Brain (https://brainspan.org/). RPKM (reads per kilobase transcripts per million mapped reads) values for *ARX* gene were mined and normalized for each sample using the following formula RPKM(gene)/(∑RPKM(all genes in the sample)) X 10^6^. Graphpad Prism was used to plot ARX normalized counts from 8 weeks post-conception (wpc) to 40 years of age.

### Statistical analysis

Outliers were removed from each dataset using GraphPad Prism software (Q=1%). A two-tailed Student’s *t*-test was used to compare the mean ± standard error of the mean (SEM) values, with Welch’s correction when the F-test indicated significant differences between the variances of both groups. All analyses were carried out with GraphPad Prism software and the differences were considered as statistically significant when P<0.05. Effect size (d) was calculated using: https://www.psychometrica.de/effect_size.html. A large effect size was considered when d ≥ 0.8.

## Supporting information

Supplemental excel sheets

## List of Supplementary Materials

Figures S1 to S27

Tables S1 to S3

Data file S1 (Excel file)

Videos S1 to S4

## Acknowledgments

We would like to thank Raul Wilshire, Mia Kay Margarete Soerensen and Yanessa Vitela for their technical support, and Sean C. Goetsch and Jay W. Schneider for their initial help with PSC cultures. We also thank Eric D. Marsh (CHOP), Cheryl Shoubridge (The University of Adelaide) and Mary-Colette Lybrand for patient recruitment. We would like to thank Aline McKenzie for manuscript editing; and Eric D. Marsh (CHOP) and Jeffrey A. Golden (Cedars-Sinai), and their labs, for critical discussion of the manuscript. We would like to thank Sonal Goswami (UTSA) for her help on MEA and patch-clamp recordings. We thank the UTSA Genomics Core and Sean Vargas for single cell library preparations. We also acknowledge the members of Research Computing Support Group at UTSA. Computational work in this manuscript was supported by High-Performance Clusters at UTSA and Texas Advanced Computing Center (TACC) at UT Austin. We also thank Drs. Morohashi and Kitamura for the ARX antibody. Some figures were created with BioRender.com.

## Author contributions

Conceptualization, V.N.-E., P.V., Z.R.L., D.M.T. and J.H.; Methodology, V.N.-E., P.V., Z.R.L. and J.H.; Software, P.V.; Formal Analysis, V.N.-E., P.V., S.M., A.H., and Z.R.L.; Investigation, V.N.-E., P.V., S.M., Z.R.L., J.C., S.G.-A., C.N. and M.J.S.; Resources, C.N. and D.M.T., Writing – Original Draft V.N.-E.; Writing – Review & Editing, V.N.-E., P.V., S.M., Z.R.L. and J.H; Visualization, V.N.-E., P.V., Z.R.L. and J.H; Supervision, V.N.-E, M.G. and J.H.; Funding Acquisition, V.N.-E. and J.H.

## Data and materials availability

The scRNA-seq datasets presented in this study can be found in NIH repositories (accession number GSE215362). The codes used in this study can be found on https://github.com/parulvarma123/ARX_Organoids_scRNAseq.

---

**Figure S1.**
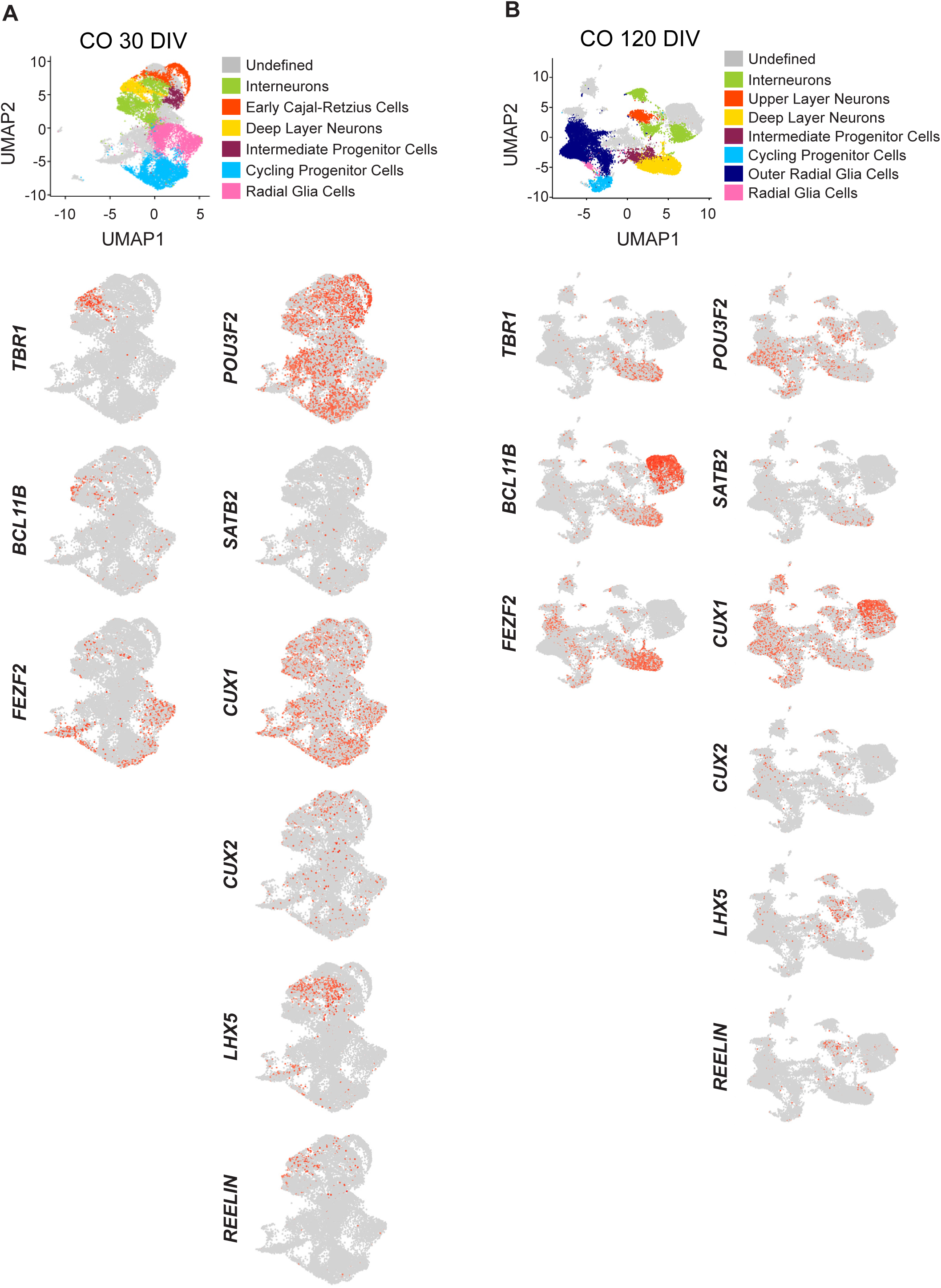
Cortical organoid characterization. **(A)** UMAP plot from control COs at 30 DIV and feature plots showing *FEZF2*, *BCL11B*, *TBR1*, *POU3F2*, *SATB2*, *CUX1*, *CUX2*, *LHX5* and *REELIN* expression. **(B)** UMAP plot from control COs at 120 DIV and feature plots showing *FEZF2*, *BCL11B*, *TBR1*, *POU3F2*, *SATB2*, *CUX1*, *CUX2*, *LHX5* and *REELIN* expression. N= 3-4 organoids from 1 clone x 3 lines.

**Figure S2.**
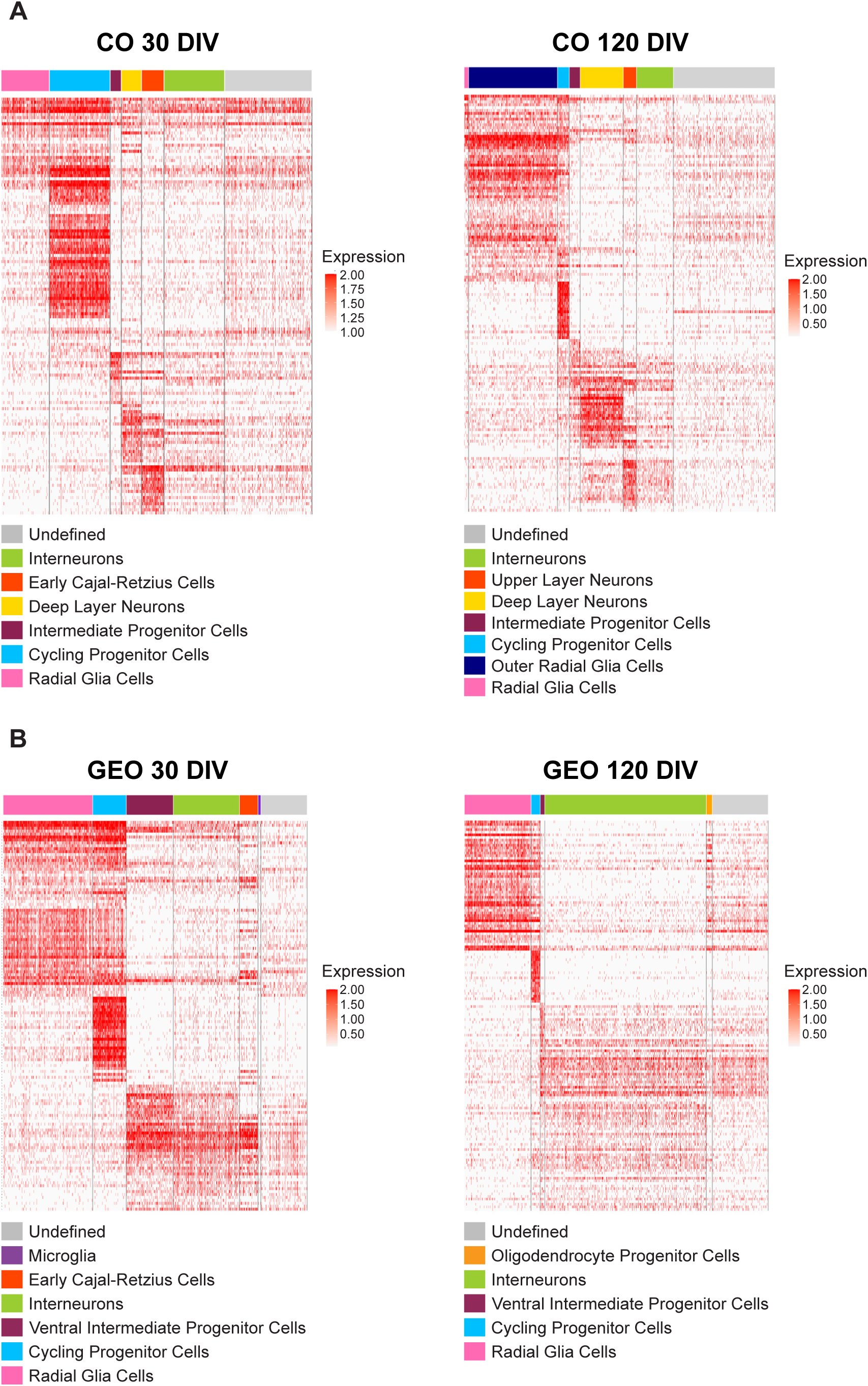
Gene expression in human neural organoids. Heatmaps show gene expression for each cluster in COs and GEOs at 30 and 120 DIV. N= 3-4 organoids from 1 clone x 3 lines.

**Figure S3.**
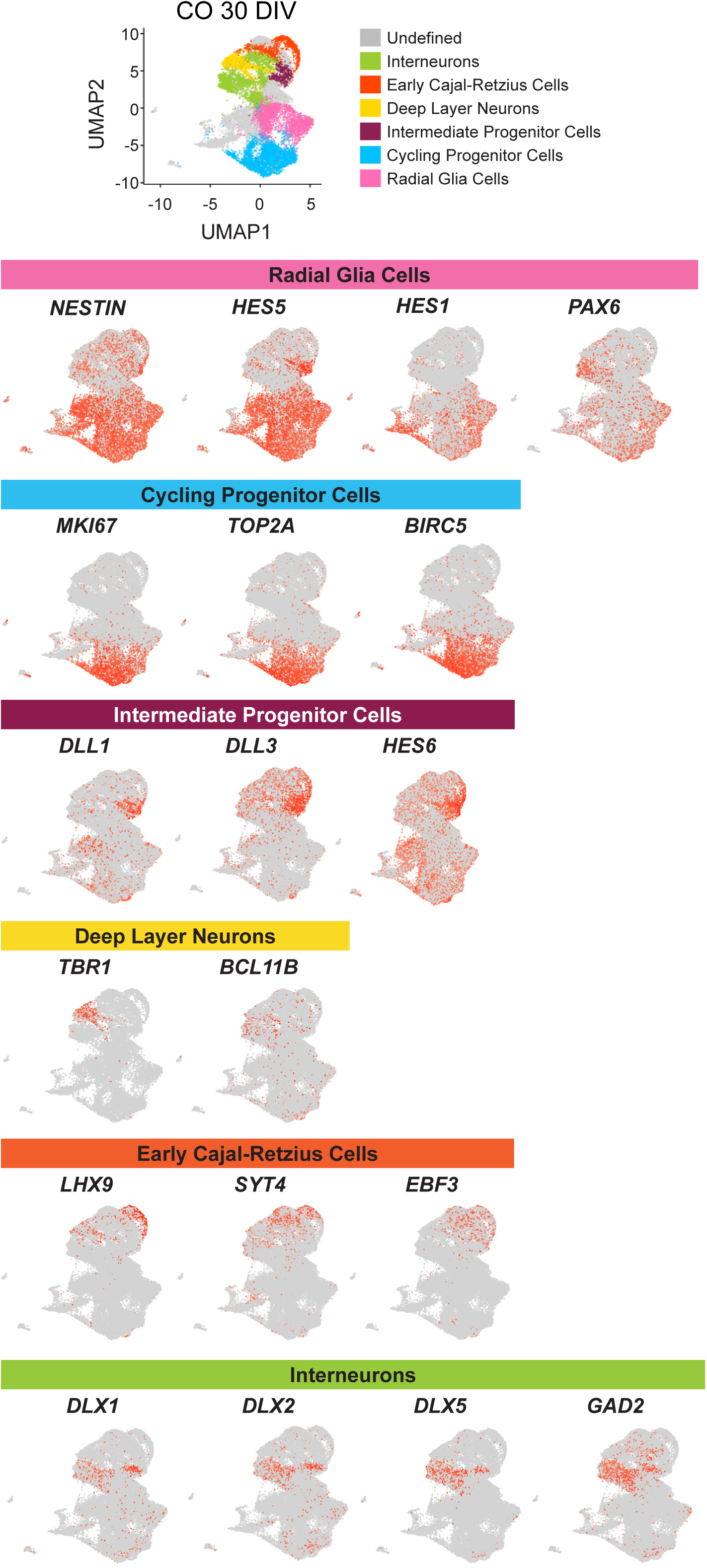
Cortical organoid characterization at 30 DIV. UMAP plot from control COs at 30 DIV and feature plots showing expression of top representative genes per cluster. N= 3-4 organoids from 1 clone x 3 lines.

**Figure S4.**
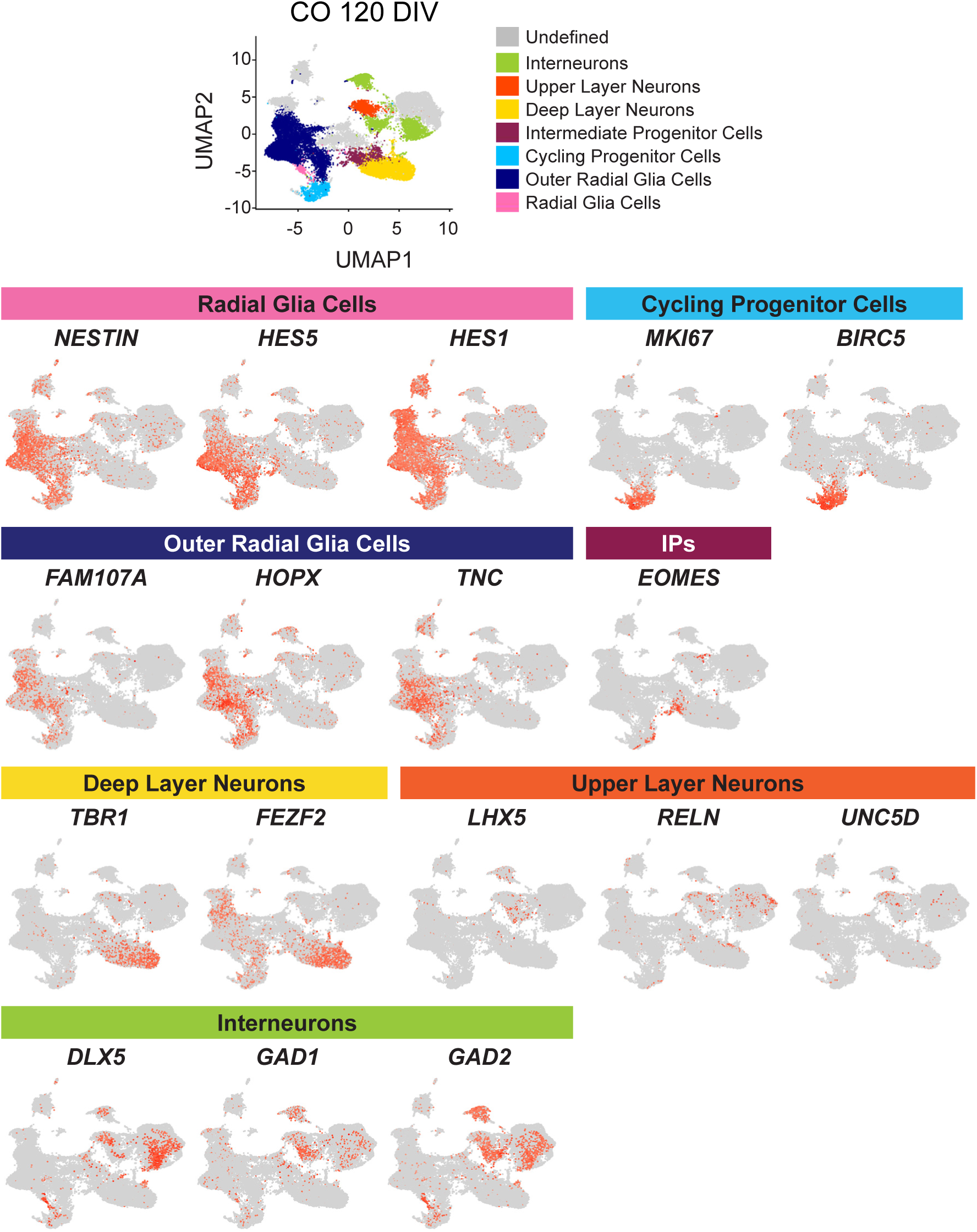
Cortical organoid characterization at 120 DIV. UMAP plot from control COs at 120 DIV and feature plots showing expression of top representative genes per cluster. N= 3-4 organoids from 1 clone x 3 lines.

**Figure S5.**
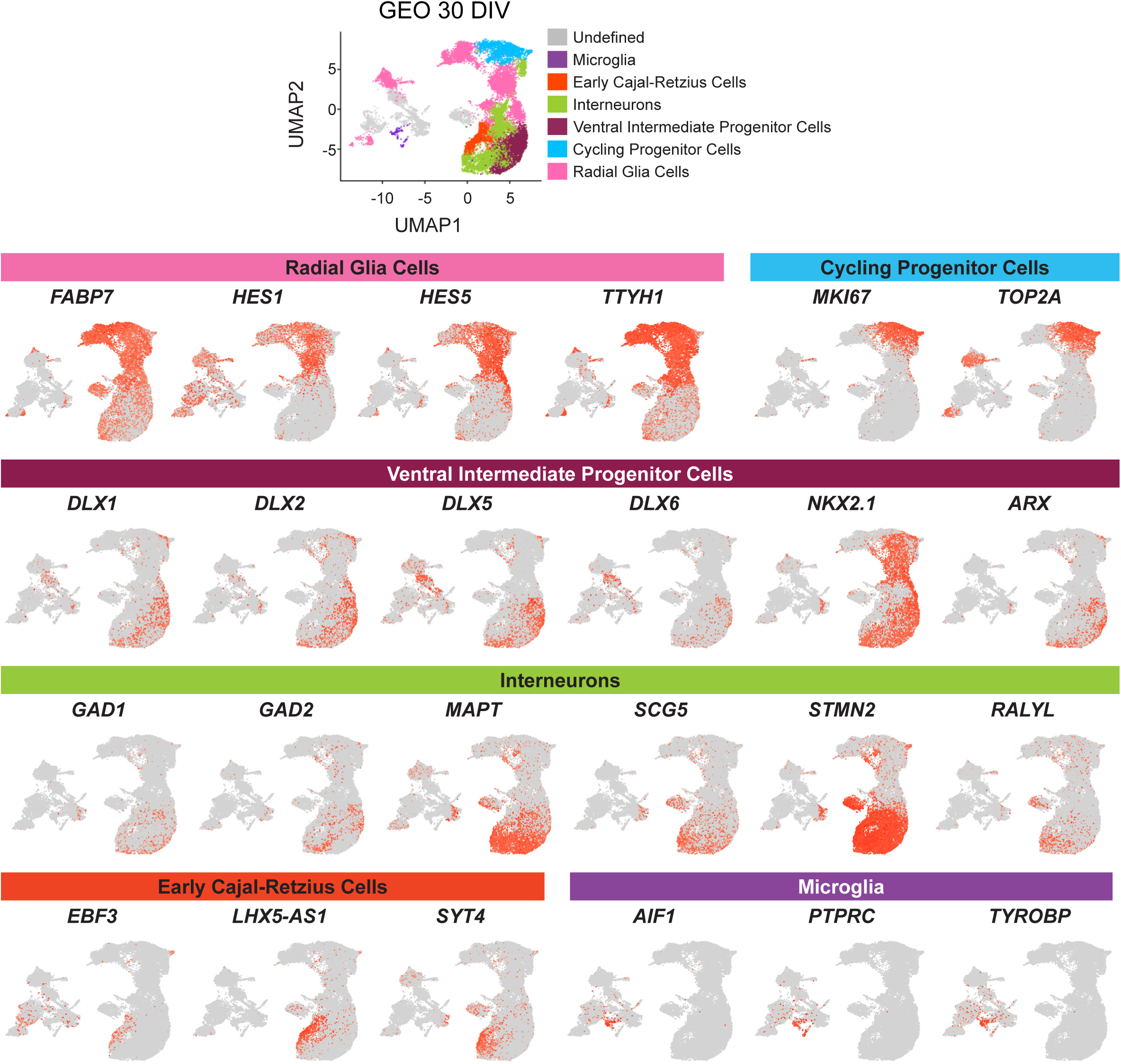
Ganglionic eminence organoid characterization at 30 DIV. UMAP plot from control GEOs at 30 DIV and feature plots showing expression of top representative genes per cluster. N= 3-4 organoids from 1 clone x 3 lines.

**Figure S6.**
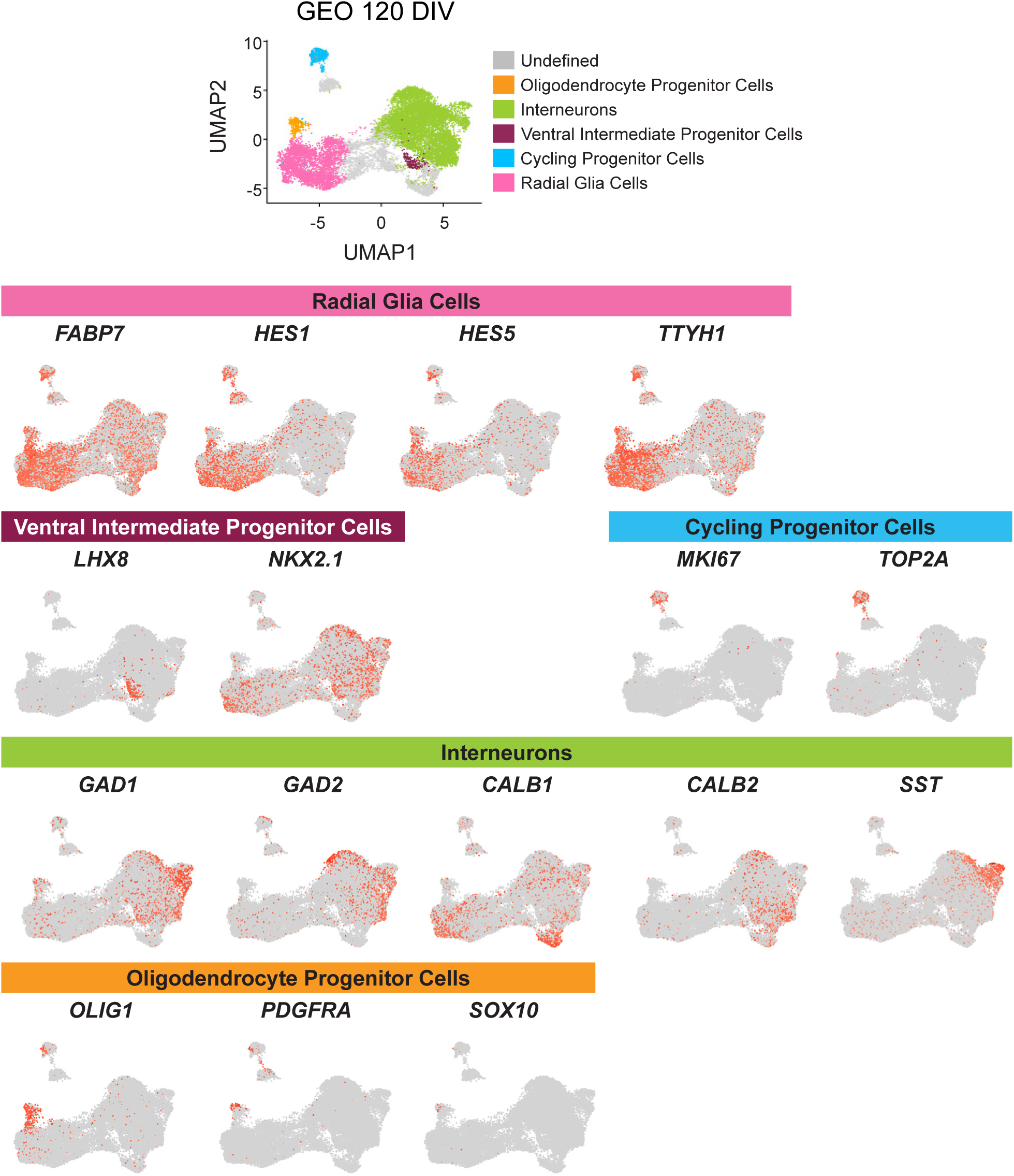
Ganglionic eminence organoid characterization at 120 DIV. UMAP plot from control GEOs at 120 DIV and feature plots showing expression of top representative genes per cluster. N= 3-4 organoids from 1 clone x 3 lines.

**Figure S7.**
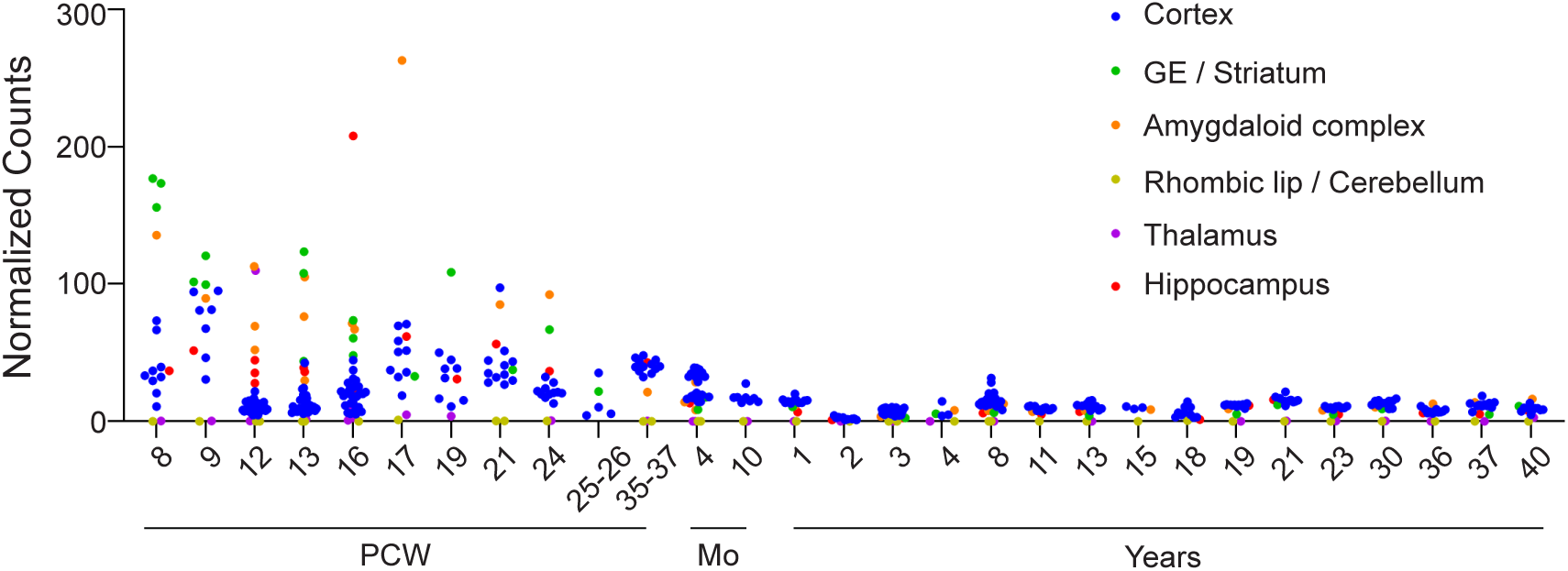
ARX expression in human tissues. Data from fetal and postnatal brain tissue demonstrating *ARX* expression in different brain regions from 8 weeks post conception (PCW) to adulthood. Green dots indicate ganglionic eminences (GE) and striatum; blue dots indicate cortical tissue. Mo = months.

**Figure S8.**
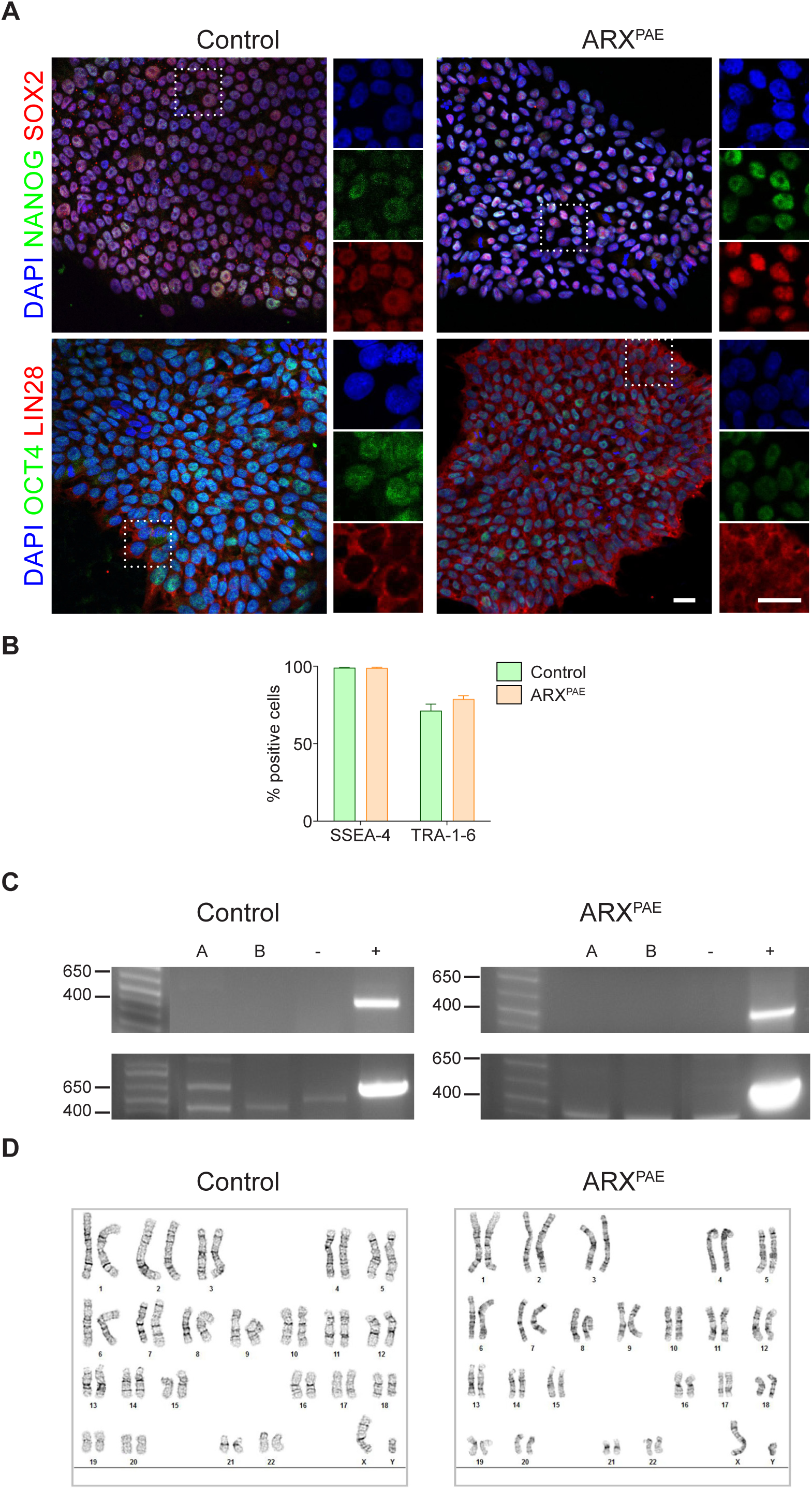
hiPSC characterization. **(A)** The images show hiPSCs colonies derived from control and ARX^PAE^ patients immunostained against NANOG, SOX2, OCT4 and LIN28; and stained with DAPI. **(B)** The graph shows the percentage of hiPSCs derived from control and ARX^PAE^ patients positive for SSEA-4 and TRA1-6 analyzed by flow cytometry. **(C)** The images show agarose gels of PCR product of reprogramming vectors. **(D)** The images show a representative karyotype from control and ARX^PAE^ hiPSCs. Scale bar = 25 µm. The results are the mean ± SEM from N= 2 independent cultures from 2 clones x 3 lines per condition.

**Figure S9.**
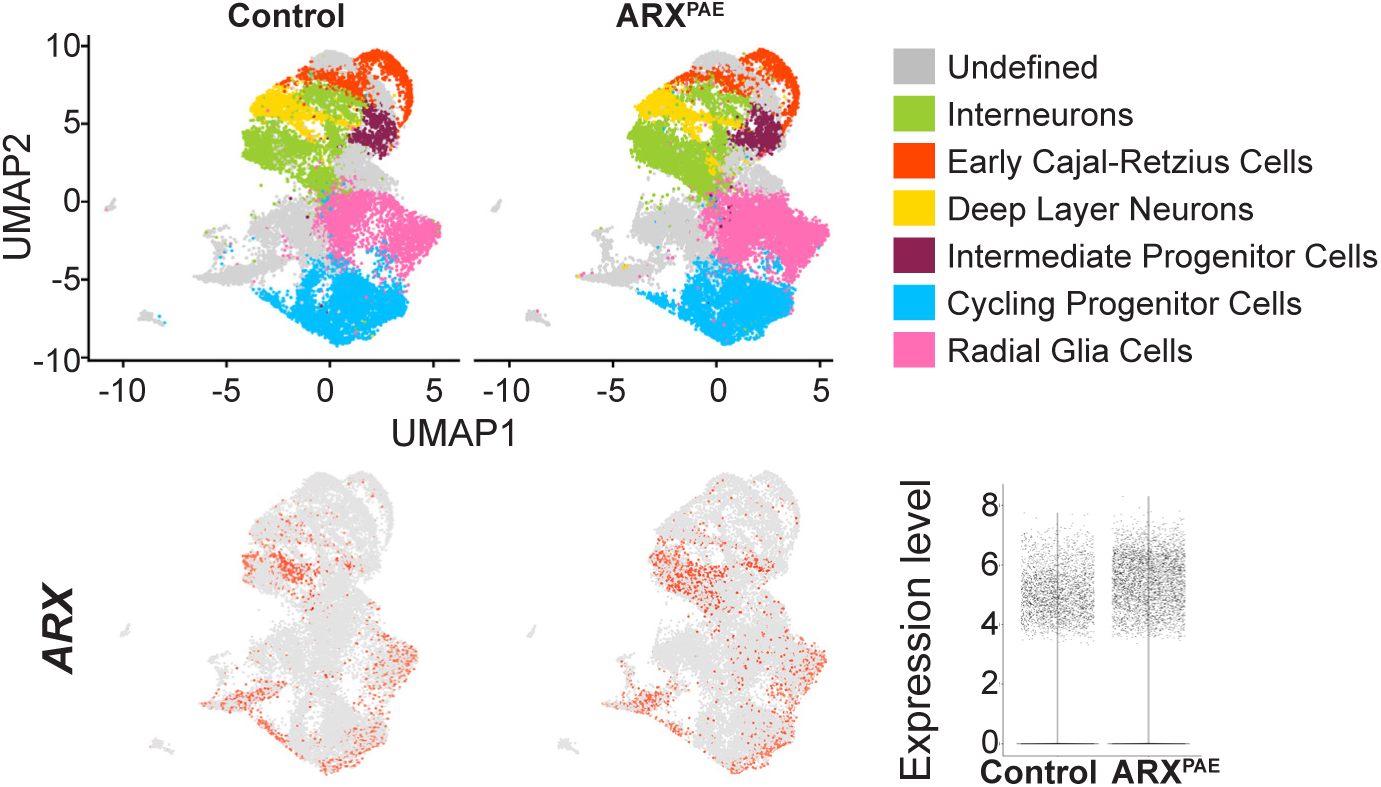
Gene expression in 30 DIV COs by scRNA-seq. UMAP plots from control and ARX^PAE^ 30 DIV COs showing the proportion of cells in each cluster and feature plots showing *ARX* expression per condition. Violin plots show the expression levels of *ARX.* N= 3-4 organoids from 1 clone x 3 lines per condition.

**Figure S10.**
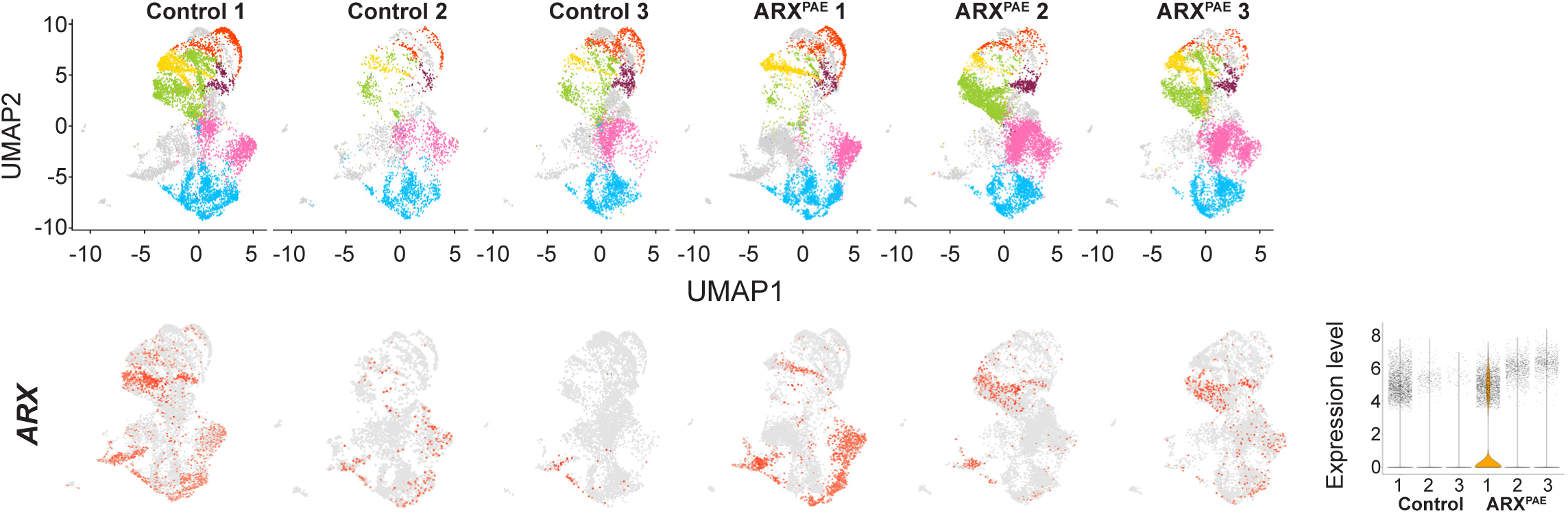
Gene expression in 30 DIV COs by scRNA-seq per line. UMAP plots from control and ARX^PAE^ 30 DIV COs showing the proportion of cells in each cluster and feature plots showing *ARX* expression per line. Violin plots show the expression levels of *ARX* per line. N= 3-4 organoids from 1 clone x 3 lines per condition.

**Figure S11.**
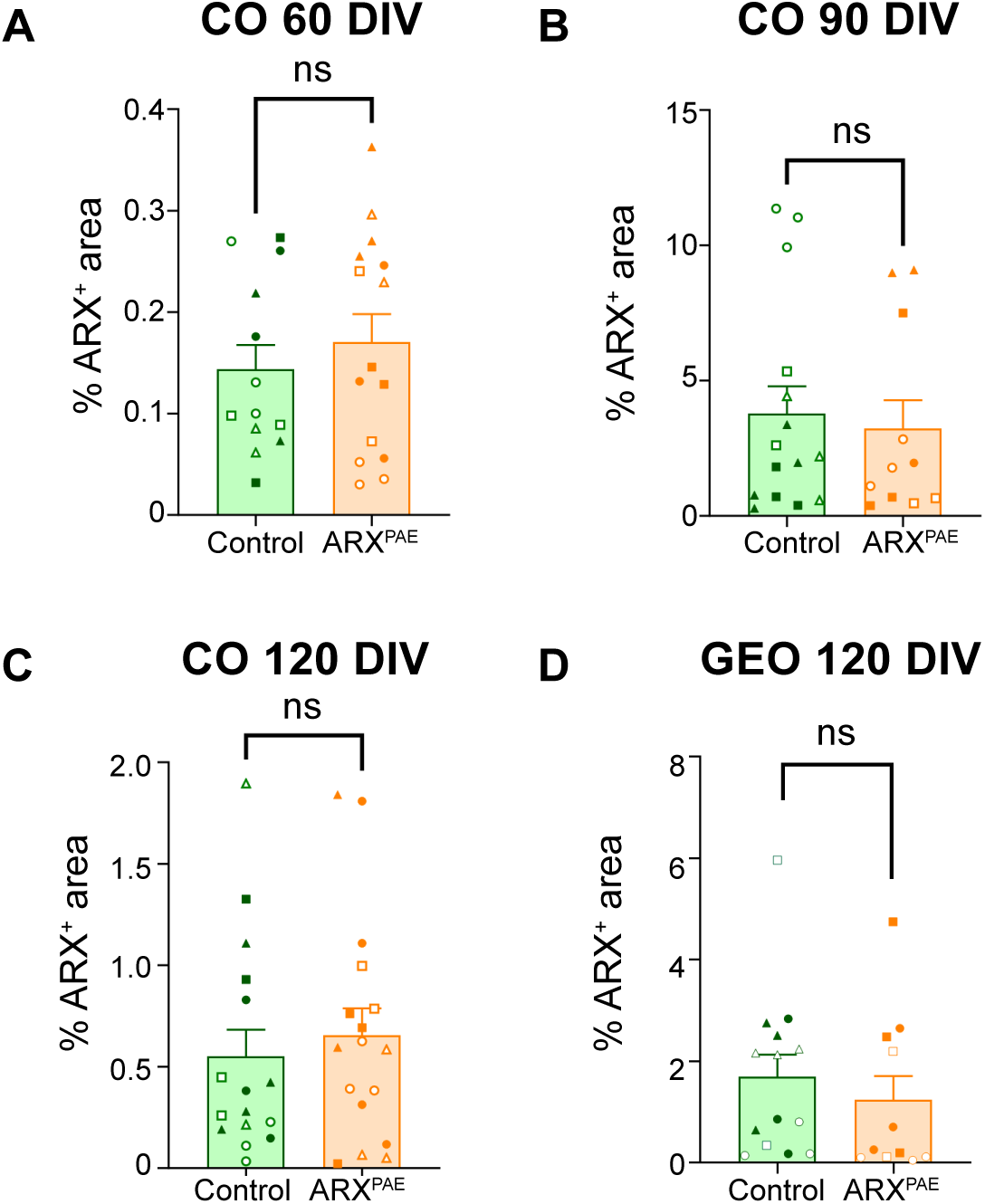
Expression of *ARX* in COs and GEOs. Graphs show the percentage of ARX^+^ area in control and ARX^PAE^ COs at 60 DIV **(A)**, 90 DIV **(B)** and 120 DIV **(C)**; and GEOs at 120 DIV **(D)**. Two-tailed Student’s *t*-test, ns = not significant. The results are the mean ± SEM from N= 18 organoids from 2 clones x 3 lines per condition. Individual points represented data from individual organoid and each symbol represented the data from each clone/line (Line 1: Clone A ● Clone B ○; Line 2: Clone A ◼ Clone B ◻; Line 3: Clone A ▲ Clone B △).

**Figure S12.**
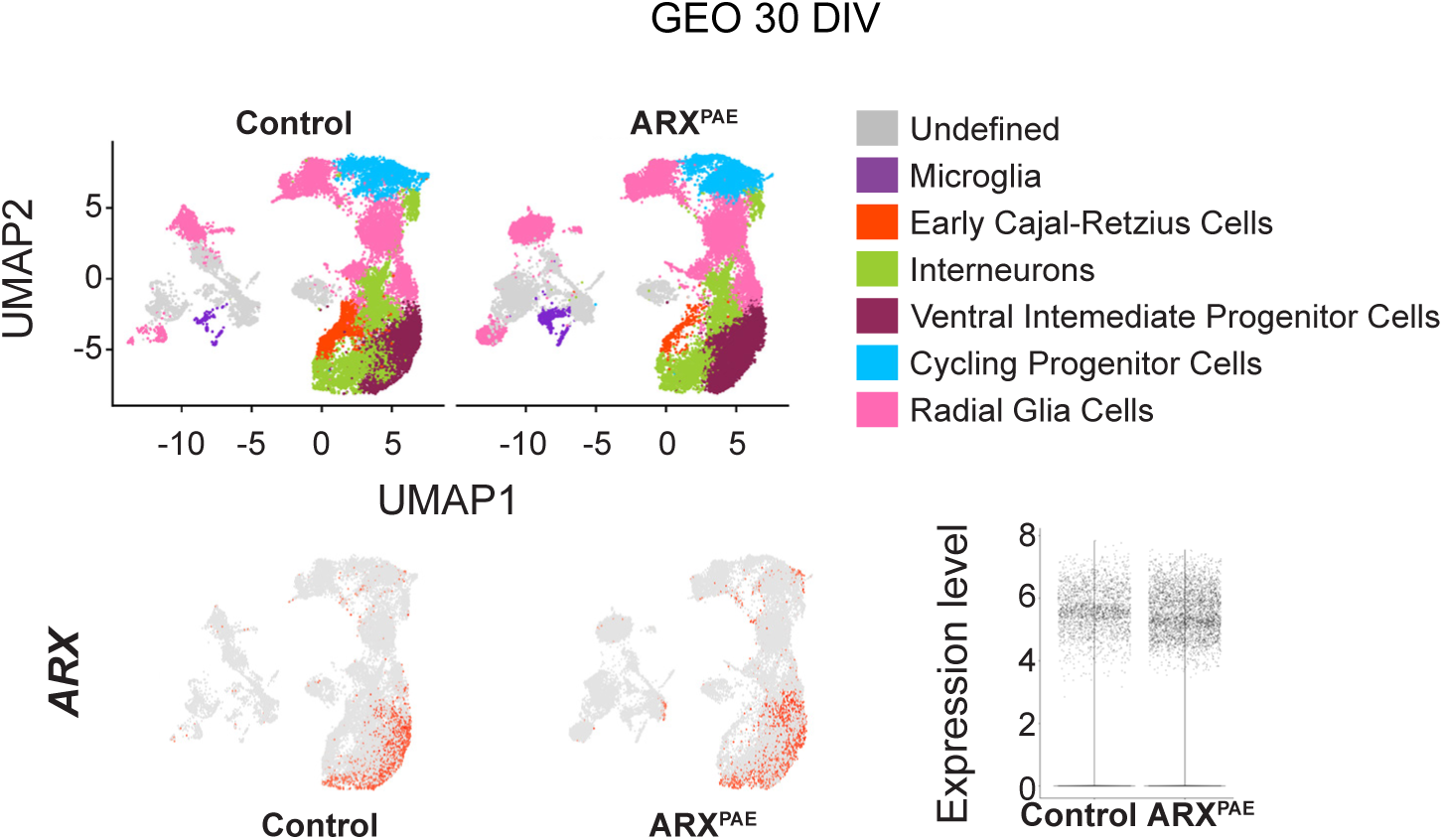
Gene expression in 30 DIV GEOs by scRNA-seq. UMAP plots from control and ARX^PAE^ 30 DIV GEO showing the proportion of cells in each cluster and feature plots showing *ARX* expression per condition. Violin plot shows the expression level of *ARX.* N= 3-4 organoids from 1 clone x 3 lines per condition.

**Figure S13.**
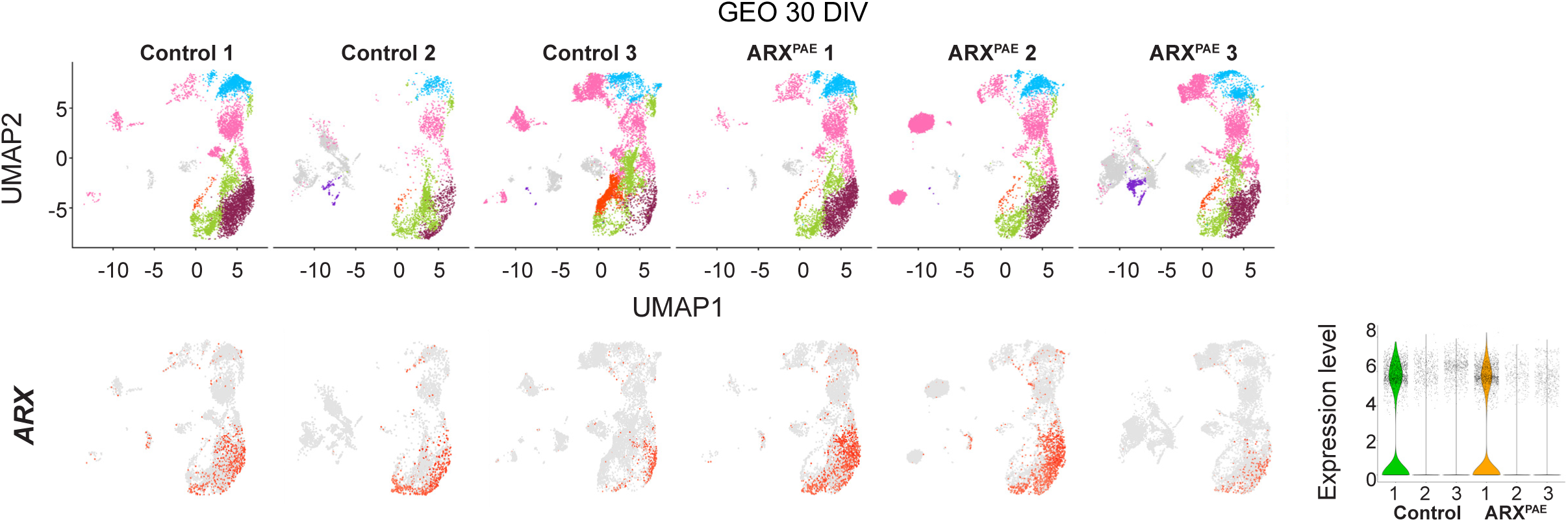
Gene expression in 30 DIV GEOs by scRNA-seq per line. UMAP plots from control and ARX^PAE^ 30 DIV GEO showing the proportion of cells in each cluster and feature plots showing *ARX* expression per line. Violin plot shows the expression level of *ARX.* N= 3-4 organoids from 1 clone x 3 lines per condition.

**Figure S14.**
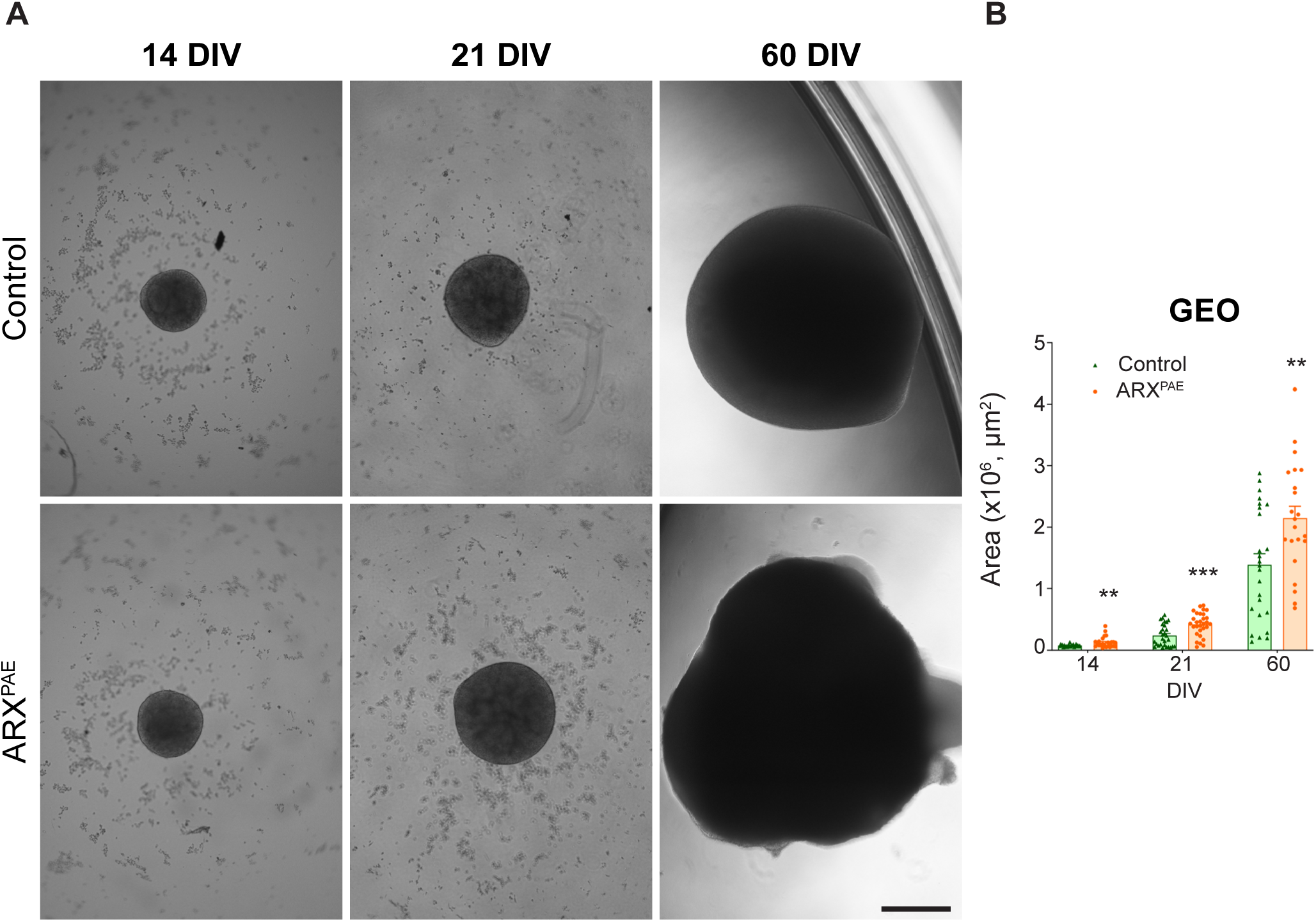
Poly-Alanine expansion in ARX promotes GEO growth. **(A)** The representative bright field images of control and ARX^PAE^ GEOs at 14, 21 and 60 DIV. **(B)** Graph shows the area of control and ARX^PAE^ GEOs at 14, 21 and 60 DIV. Scale bar = 500 µm. DIV = days *in vitro*, GEOs = ganglionic eminence organoids. Two-tailed Student’s *t*-test, ** = p < 0.01, *** = p < 0.001, ns = not significant. The results are the mean ± SEM from N= 22-30 organoids from 2 clones x 3 lines per condition.

**Figure S15.**
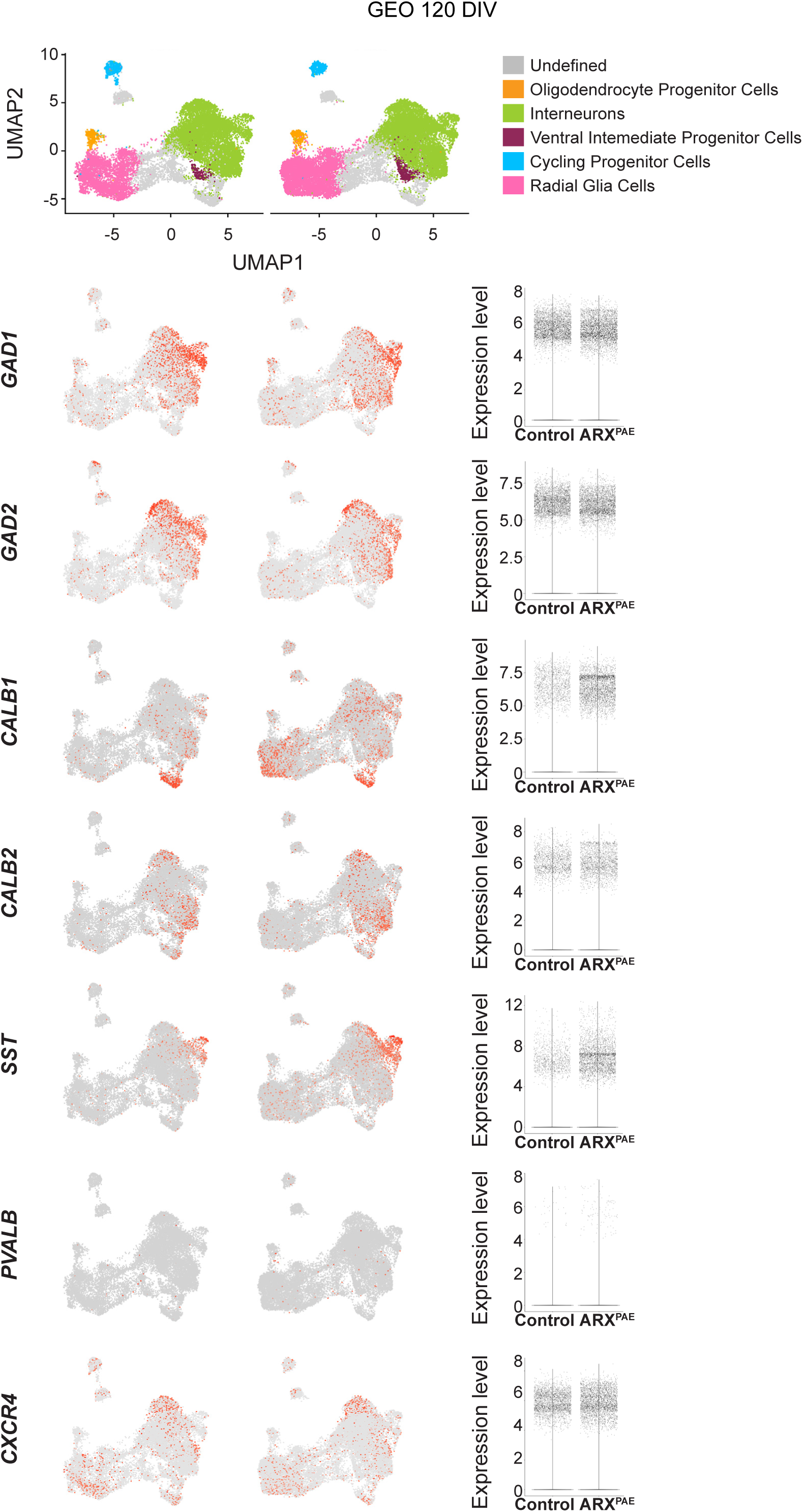
Gene expression in 120 DIV GEOs by scRNA-seq. UMAP plots from control and ARX^PAE^ 120 DIV GEOs showing the proportion of cells in each cluster and feature plots showing *GAD1*, *GAD2, CALB1, CALB2, SST, PVALB* and *CXCR4* expression per condition. Violin plots show the expression levels of *GAD1*, *GAD2, CALB1, CALB2, SST*, *PVALB* and *CXCR4.* N= 3-4 organoids from 1 clone x 3 lines per condition.

**Figure S16.**
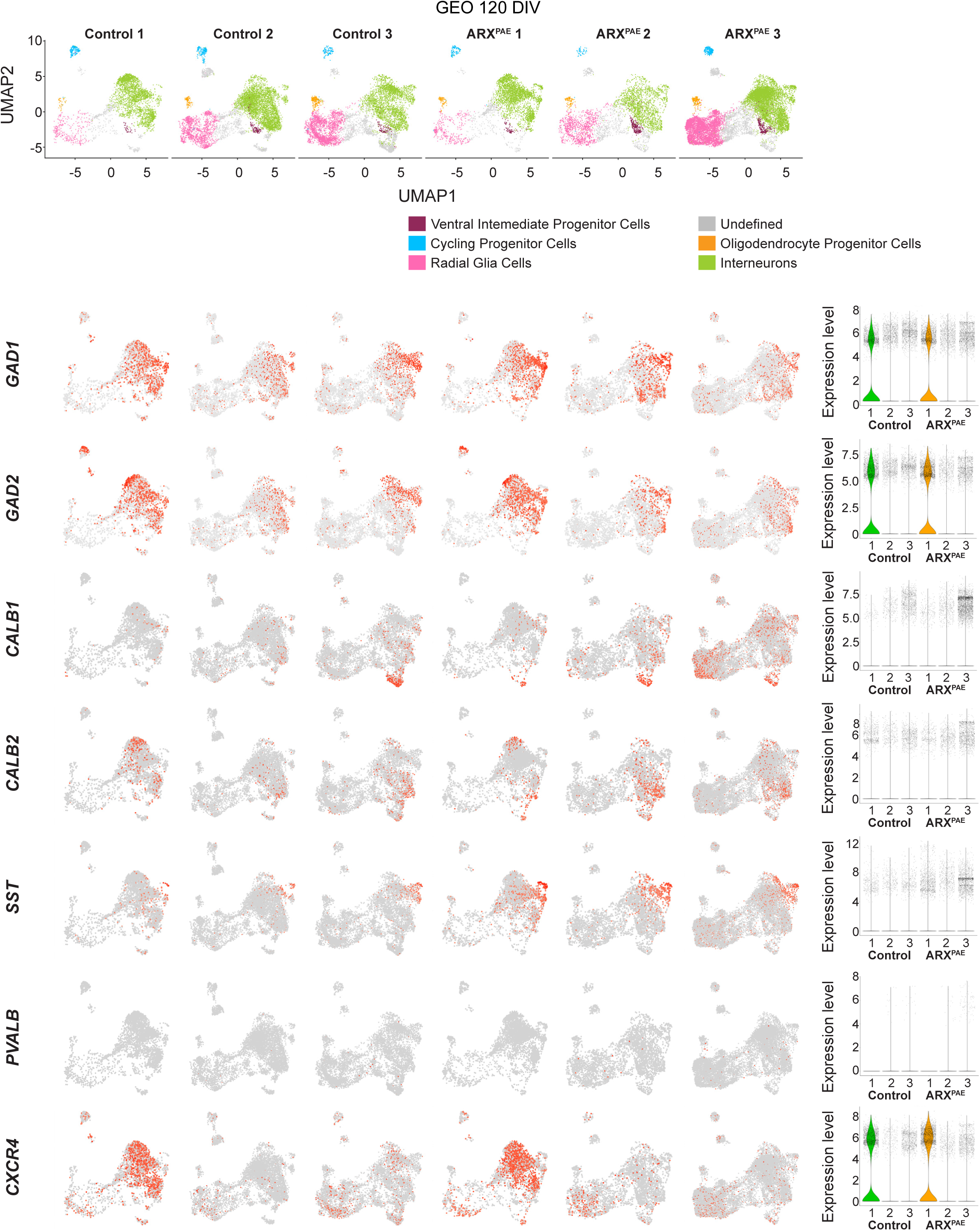
Gene expression in 120 DIV GEOs by scRNA-seq per line. UMAP plots from control and ARX^PAE^ 120 DIV GEOs showing the proportion of cells in each cluster and feature plots showing *GAD1*, *GAD2, CALB1, CALB2, SST, PVALB* and *CXCR4* expression per line. Violin plots show the expression levels of *GAD1*, *GAD2, CALB1, CALB2, SST, PVALB* and *CXCR4* per line. N= 3-4 organoids from 1 clone x 3 lines per condition.

**Figure S17.**
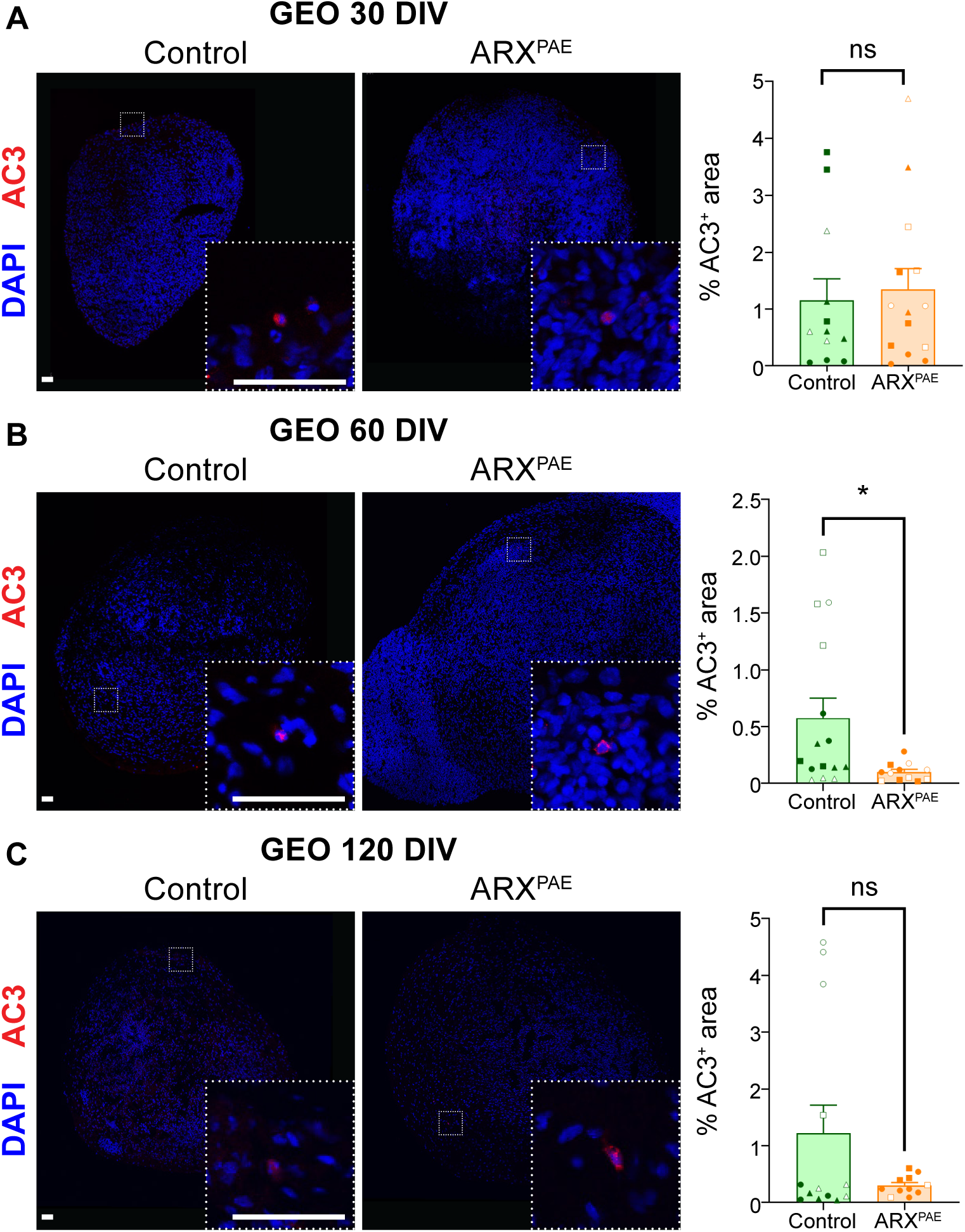
ARX^PAE^ does not promote cell death in GEOs. **(A)** The images show control and ARX^PAE^ GEOs at 30 DIV immunostained against AC3 and stained with DAPI. Graph shows the percentage of AC3^+^ area in control and ARX^PAE^ GEOs at 30 DIV. **(B)** The images show control and ARX^PAE^ GEOs at 60 DIV immunostained against AC3 and stained with DAPI. Graph shows the percentage of AC3^+^ area in control and ARX^PAE^ GEOs at 60 DIV. **(C)** The images show control and ARX^PAE^ GEOs at 120 DIV immunostained against AC3 and stained with DAPI. Graph shows the percentage of AC3^+^ area in control and ARX^PAE^ GEOs at 120 DIV. Scale bar = 50 µm. DIV = days *in vitro*, GEOs = ganglionic eminence organoids. Two-tailed Student’s *t*-test, * = p < 0.05, ns = not significant. The results are the mean ± SEM from N= 18 organoids from 2 clones x 3 lines per condition. Individual points represented data from individual organoid and each symbol represented the data from each clone/line (Line 1: Clone A ● Clone B ○; Line 2: Clone A ◼ Clone B ◻; Line 3: Clone A ▲ Clone B △).

**Figure S18.**
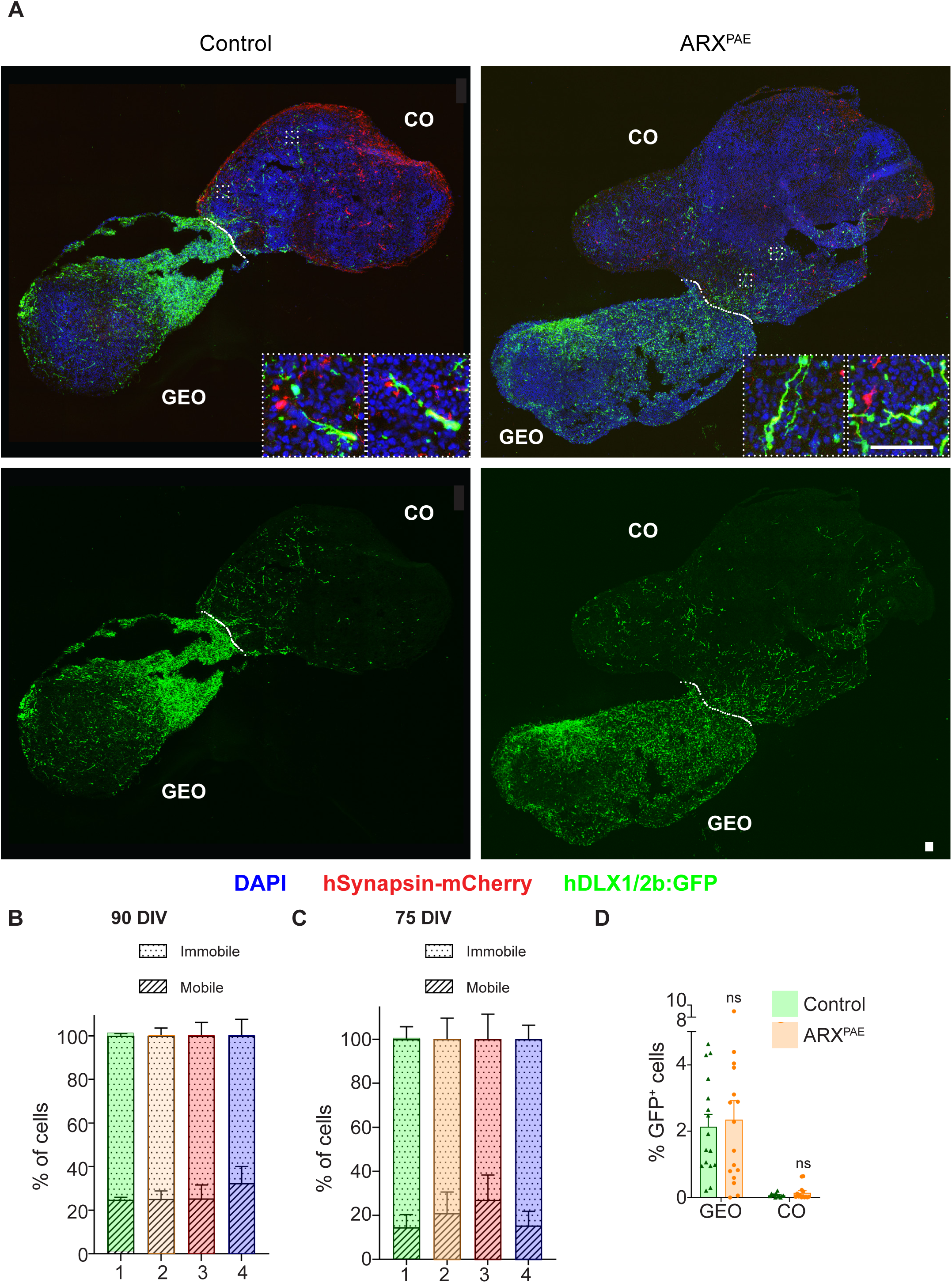
Poly-Alanine expansion does not affect the number of migrating interneurons in assembloids. **(A)** The images show control and ARX^PAE^ assembloids at 90 DIV, 30 days after fusion, immunostained against mCherry and GFP; and stained with DAPI. COs were infected with an AAV-hSYNAPSIN-mCherry virus and GEOs with a lenti-DLX1/2b:GFP virus at 50 DIV, 10 days before fusion. **(B)** The graph shows the percentage of mobile and immobile DLX1/2-GFP^+^ interneurons in all four experimental conditions at 90 DIV. **(C)** The graph shows the percentage of mobile and immobile DLX1/2-GFP^+^ interneurons in all four experimental conditions at 75 DIV. **(D)** The graph shows the percentage of DLX1/2-GFP^+^ area in COs and GEOs in control and ARX^PAE^ assembloids. COs = cortical organoids, GEOs = ganglionic eminence organoids. Scale bar = 100 µm. The results are the mean ± SEM from N= 30-48 neurons from 11-17 organoids from 2 clones x 3 lines per condition.

**Figures S19.**
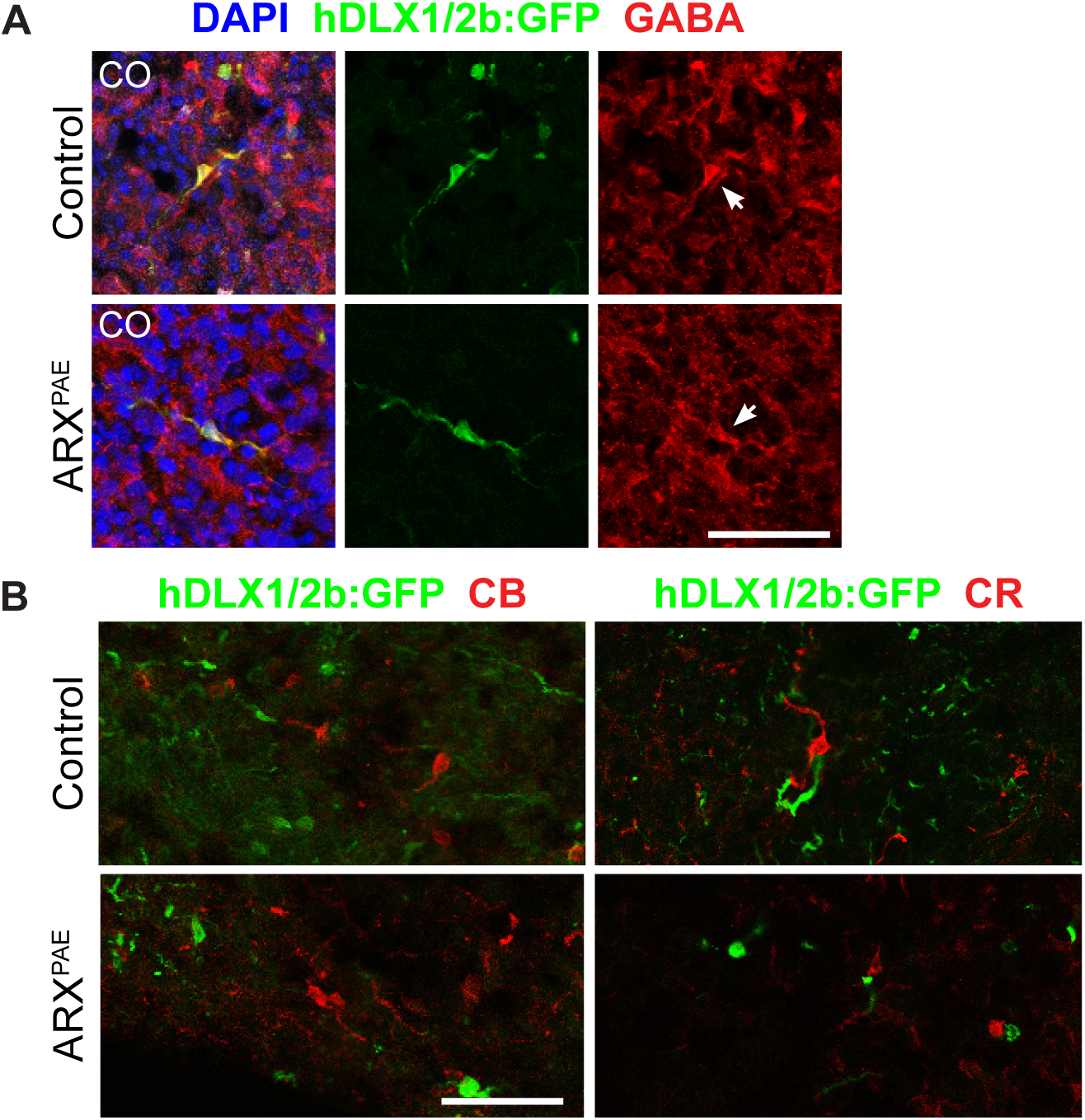
DLX1/2-GFP^+^ cells expressed GABA after migrating into the Cos. **(A)** The images show control and ARX^PAE^ assembloids at 90 DIV immunostained against GFP and GABA; and stained with DAPI. **(B)** The images show control and ARX^PAE^ assembloids at 90 DIV immunostained against GFP, CB and CR; and stained with DAPI. COs = cortical organoids. Scale bar = 50 µm. N= 30-48 neurons from 11-17 organoids from 2 clones x 3 lines per condition.

**Figure S20.**
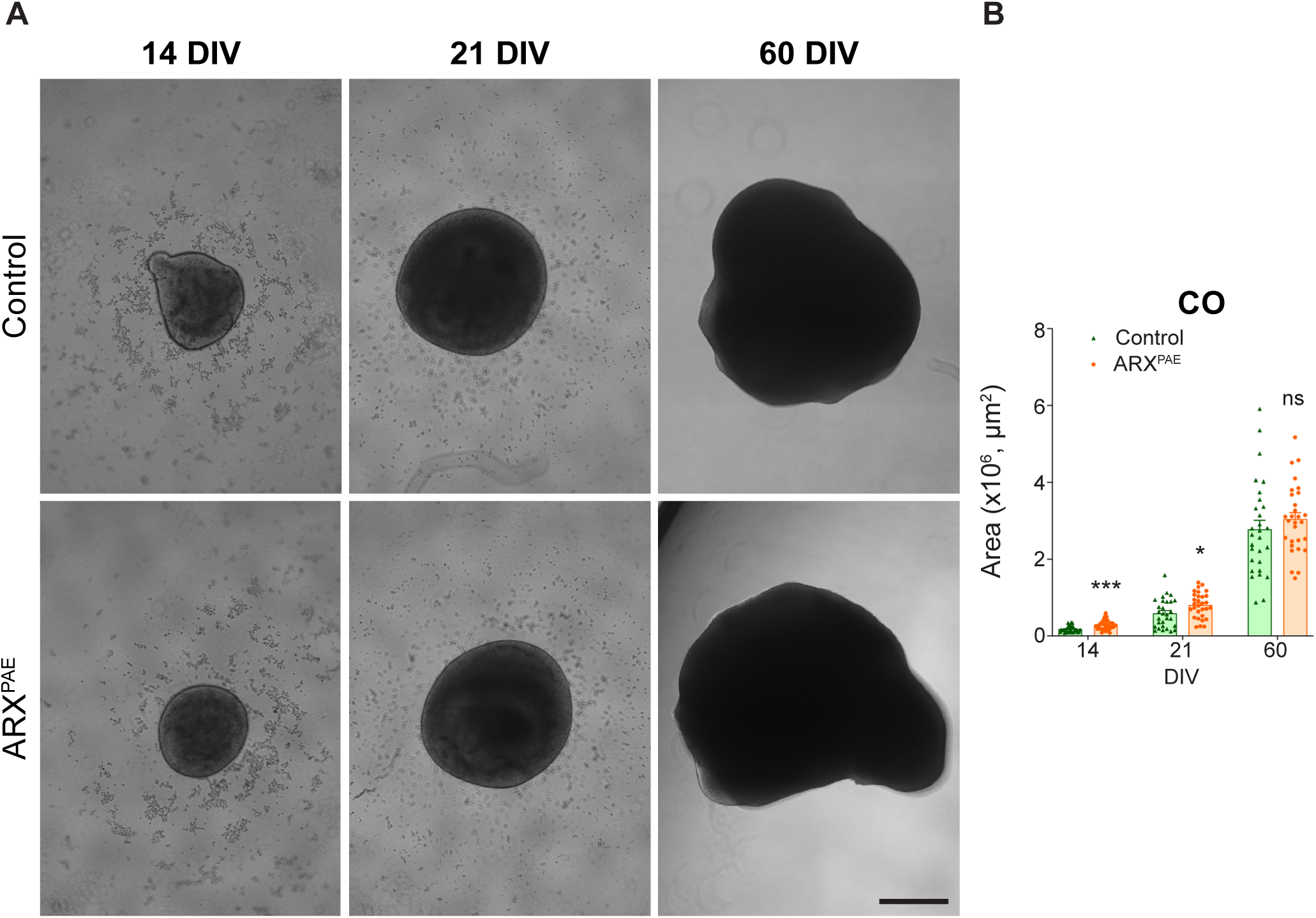
Poly-Alanine expansion in ARX promotes CO growth. **(A)** The representative bright field images of control and ARX^PAE^ COs at 14, 21 and 60 DIV. **(B)** Graph shows the area of control and ARX^PAE^ COs at 14, 21 and 60 DIV. Scale bar = 500 µm. DIV = days *in vitro*, COs = cortical organoids. Two-tailed Student’s *t*-test, * = p < 0.05, *** = p < 0.001, ns = not significant. The results are the mean ± SEM from N= 22-30 organoids from 2 clones x 3 lines per condition.

**Figure S21.**
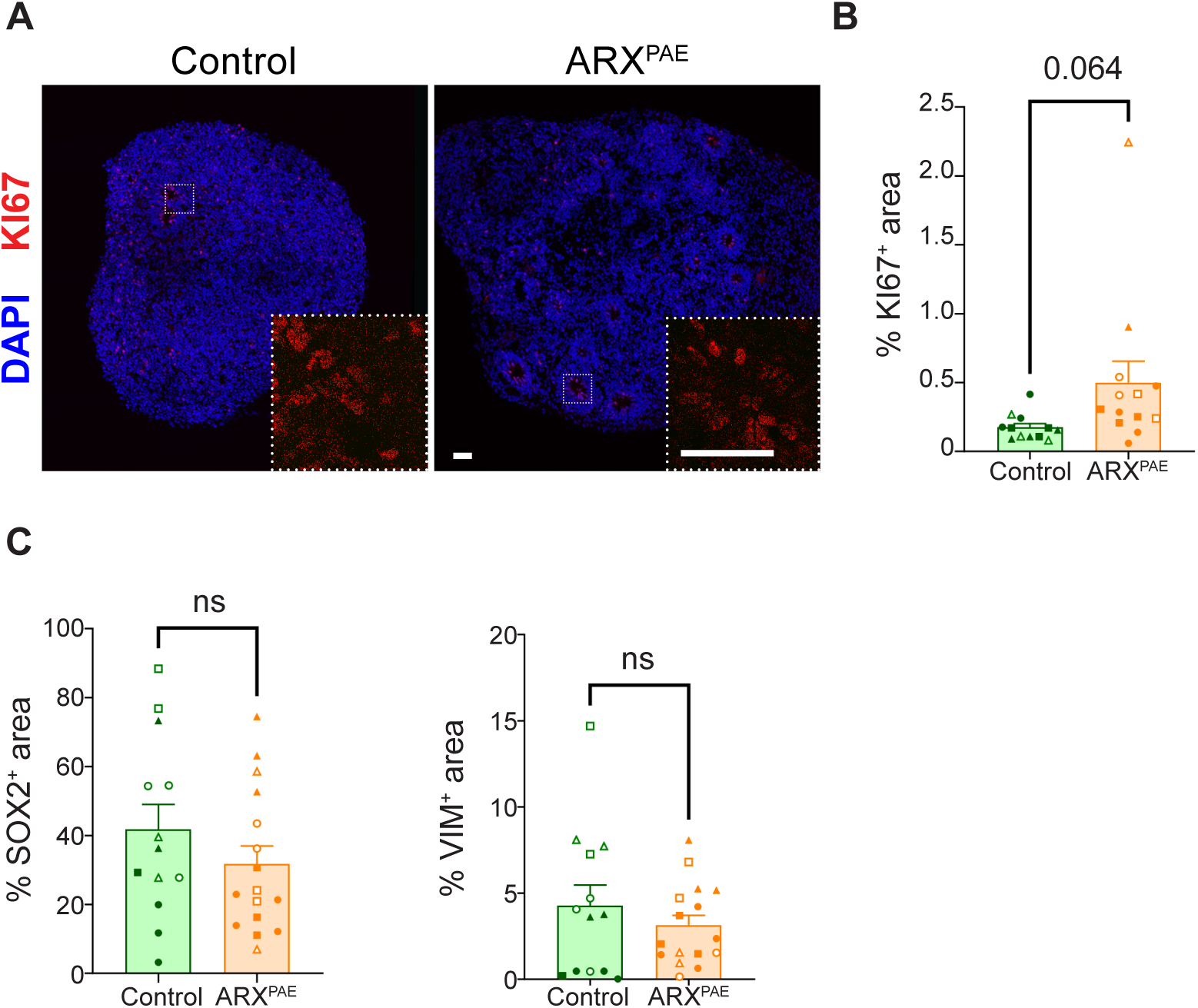
Characterization of control and ARX^PAE^ Cos. **(A)** The images show control and ARX^PAE^ COs at 30 DIV immunostained against KI67, and stained with DAPI. **(B)** Graph shows the percentage of KI67^+^ area in control and ARX^PAE^ COs at 30 DIV. **(C)** Graphs show the percentage of SOX2^+^ and VIM^+^ area in control and ARX^PAE^ COs at 30 DIV. Two-tailed Student’s *t*-test, ns = not significant. The results are the mean ± SEM from N= 18 organoids from 2 clones x 3 lines per condition. Individual points represented data from individual organoid and each symbol represented the data from each clone/line (Line 1: Clone A ● Clone B ○; Line 2: Clone A ◼ Clone B ◻; Line 3: Clone A ▲ Clone B △).

**Figure S22.**
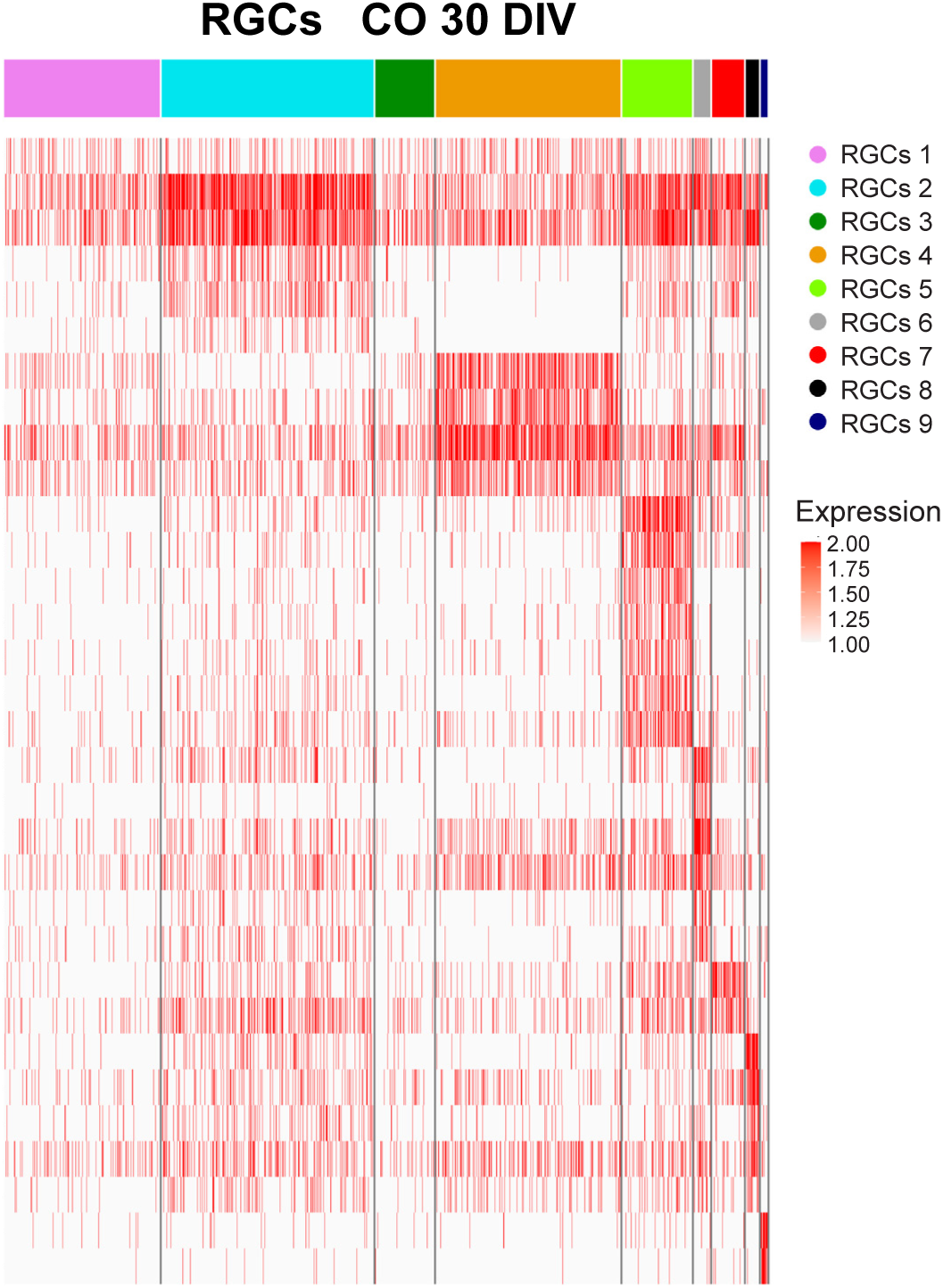
Gene expression in RGCs in 30 DIV Cos. Heatmap shows gene expression for each subcluster of RGCs in COs at 30 DIV. N= 3-4 organoids from 1 clone x 3 lines per condition.

**Figure S23.**
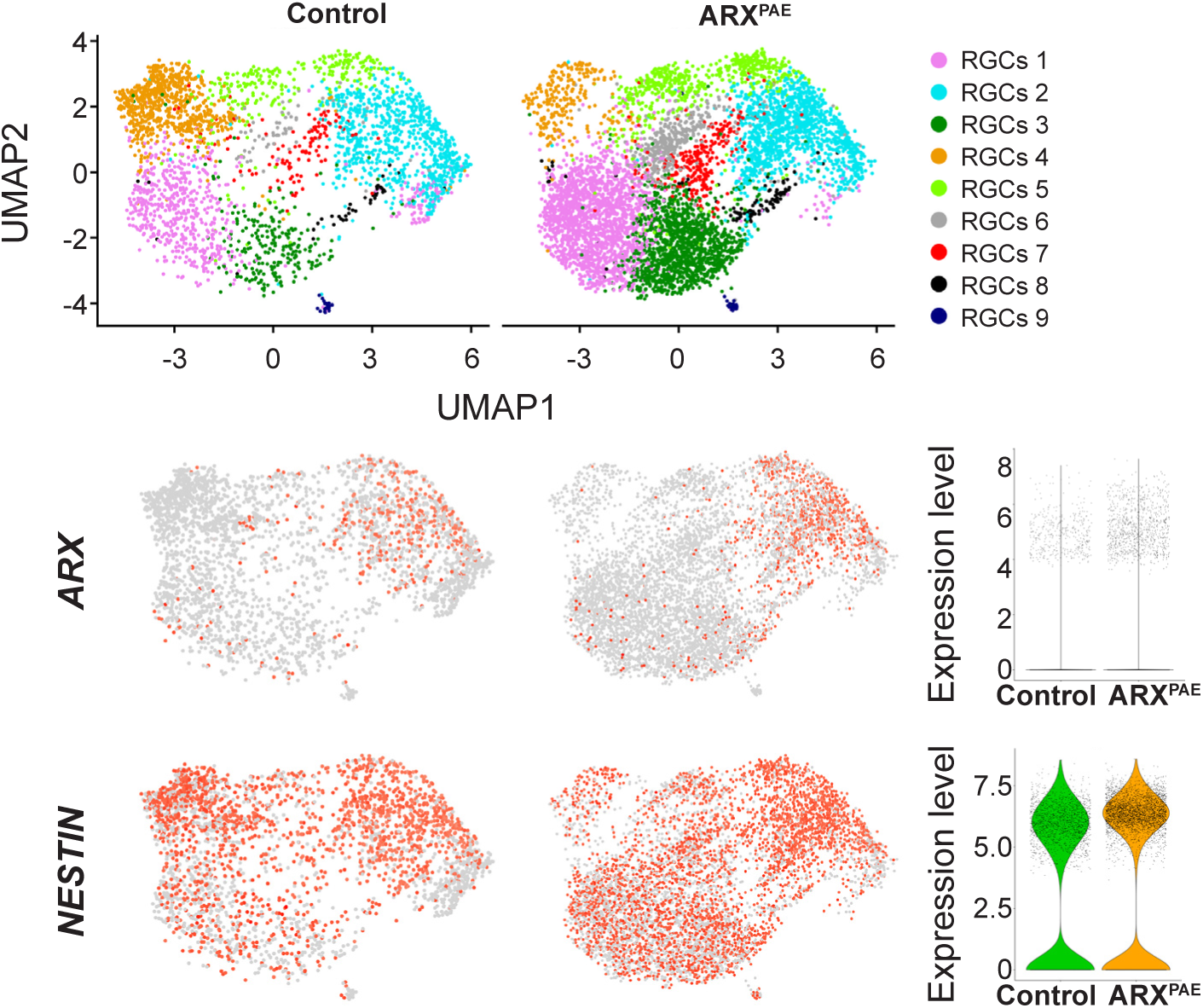
*ARX* and *NESTIN* expression in RGCs in 30 DIV Cos. UMAP plots of subclustering of RGCs from control and ARX^PAE^ COs at 30 DIV. Feature plots showing *ARX* and *NESTIN* expression in RGCs. Violin plots show the expression levels of *ARX* and *NESTIN*.

**Figure S24.**
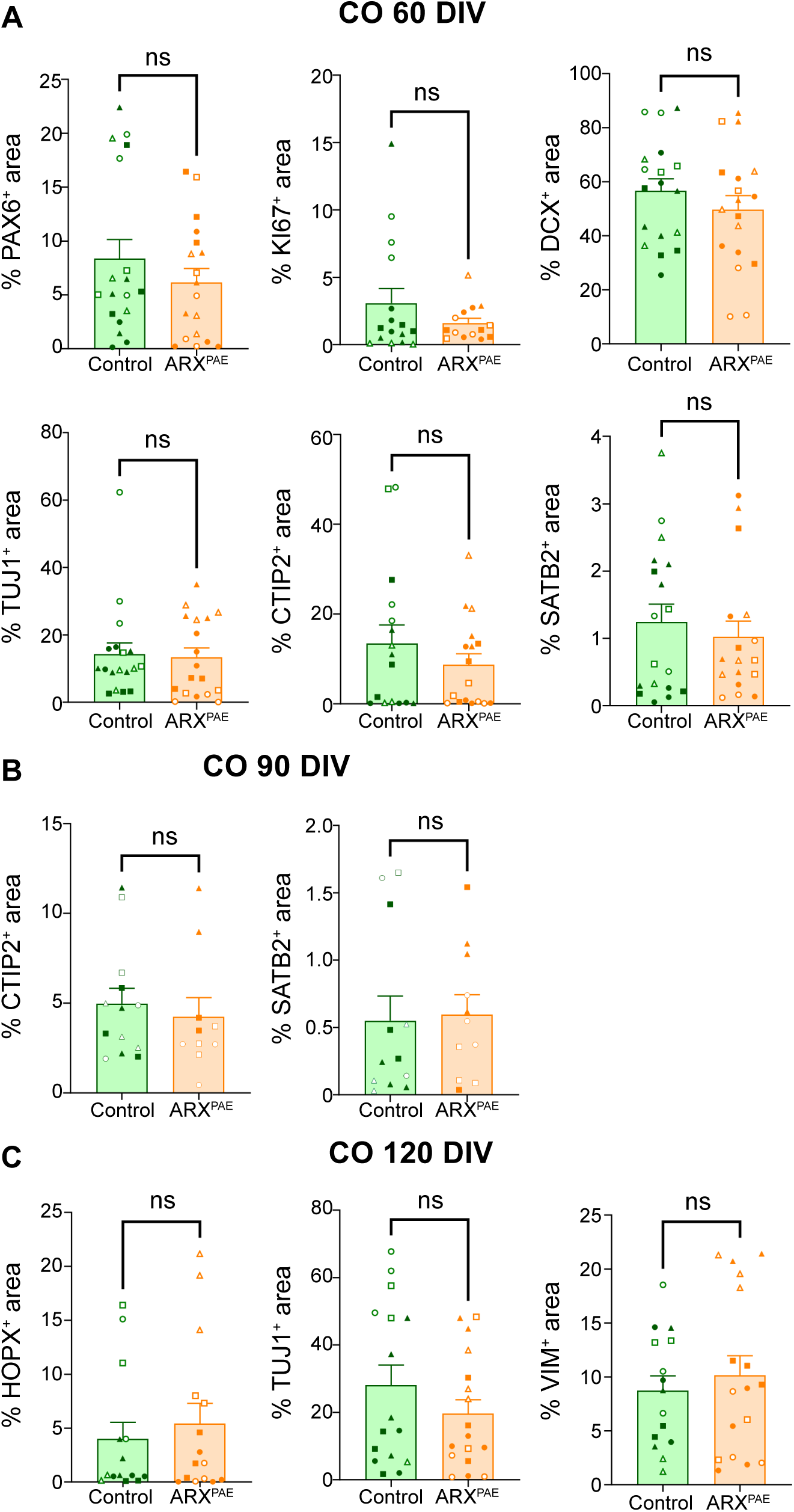
Characterization of control and ARX^PAE^ Cos. **(A)** Graphs show the percentage of PAX6^+^, KI67^+^, DCX^+^, TUJ1^+^, CTIP2^+^ and SATB2^+^ area in control and ARX^PAE^ COs at 60 DIV. **(B)** Graphs show the percentage of CTIP2^+^ and SATB2^+^ area in control and ARX^PAE^ COs at 90 DIV. **(C)** Graphs show the percentage of HOPX^+^, TUJ1^+^and VIM^+^ area in control and ARX^PAE^ COs at 120 DIV. Two-tailed Student’s *t*-test, ns = not significant. The results are the mean ± SEM from N= 18 organoids from 2 clones x 3 lines per condition. Individual points represented data from individual organoid and each symbol represented the data from each clone/line (Line 1: Clone A ● Clone B ○; Line 2: Clone A ◼ Clone B ◻; Line 3: Clone A ▲ Clone B △).

**Figure S25.**
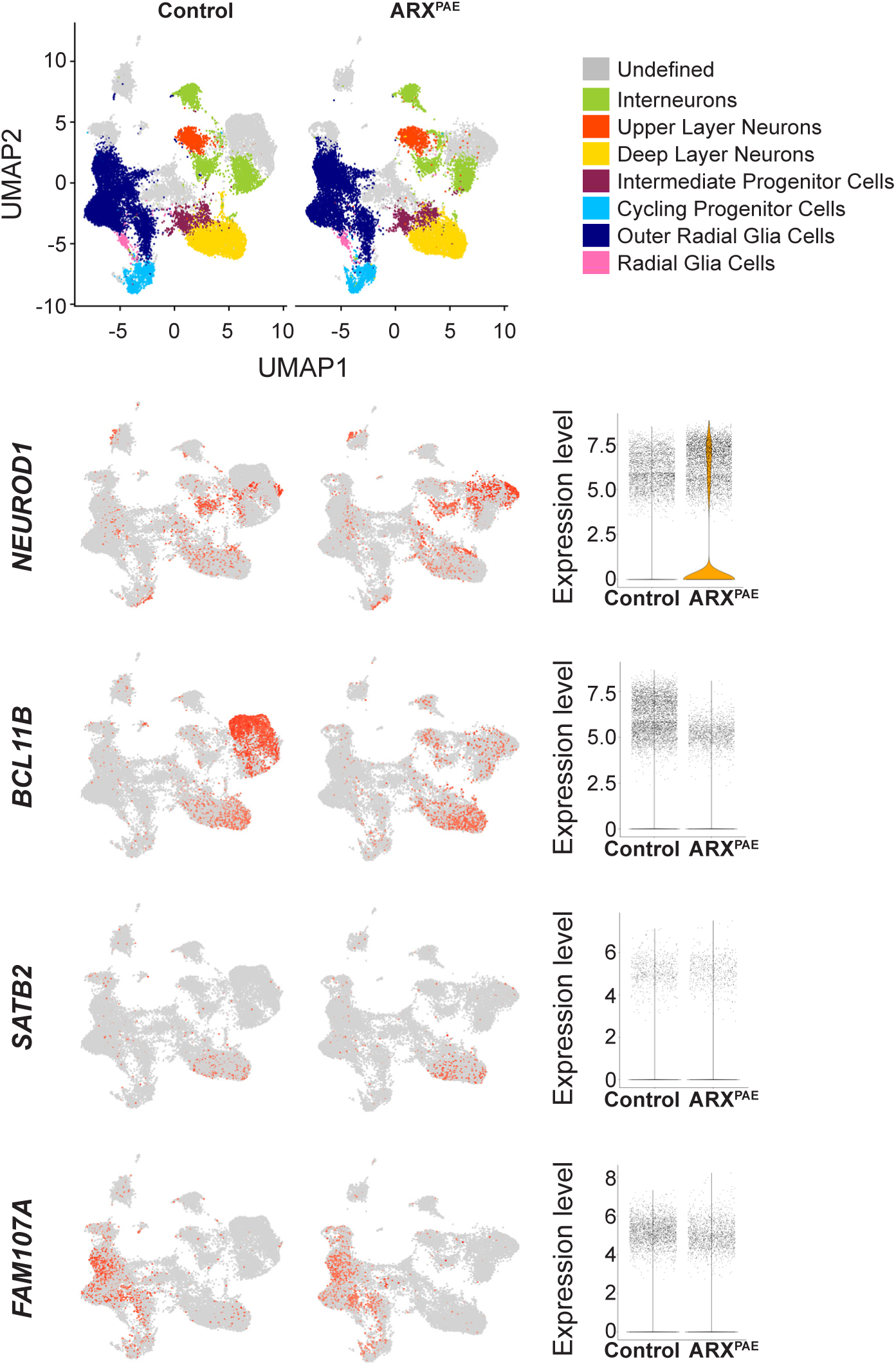
Gene expression in 120 DIV COs by scRNA-seq. UMAP plots from control and ARX^PAE^ 120 DIV COs showing the proportion of cells in each cluster and feature plots showing *NEUROD1*, *BCL11B*, *SATB2* and *FAM107A* expression per condition. Violin plots show the expression levels of *NEUROD1*, *BCL11B*, *SATB2* and *FAM107A.* N= 3-4 organoids from 1 clone x 3 lines per condition.

**Figure S26.**
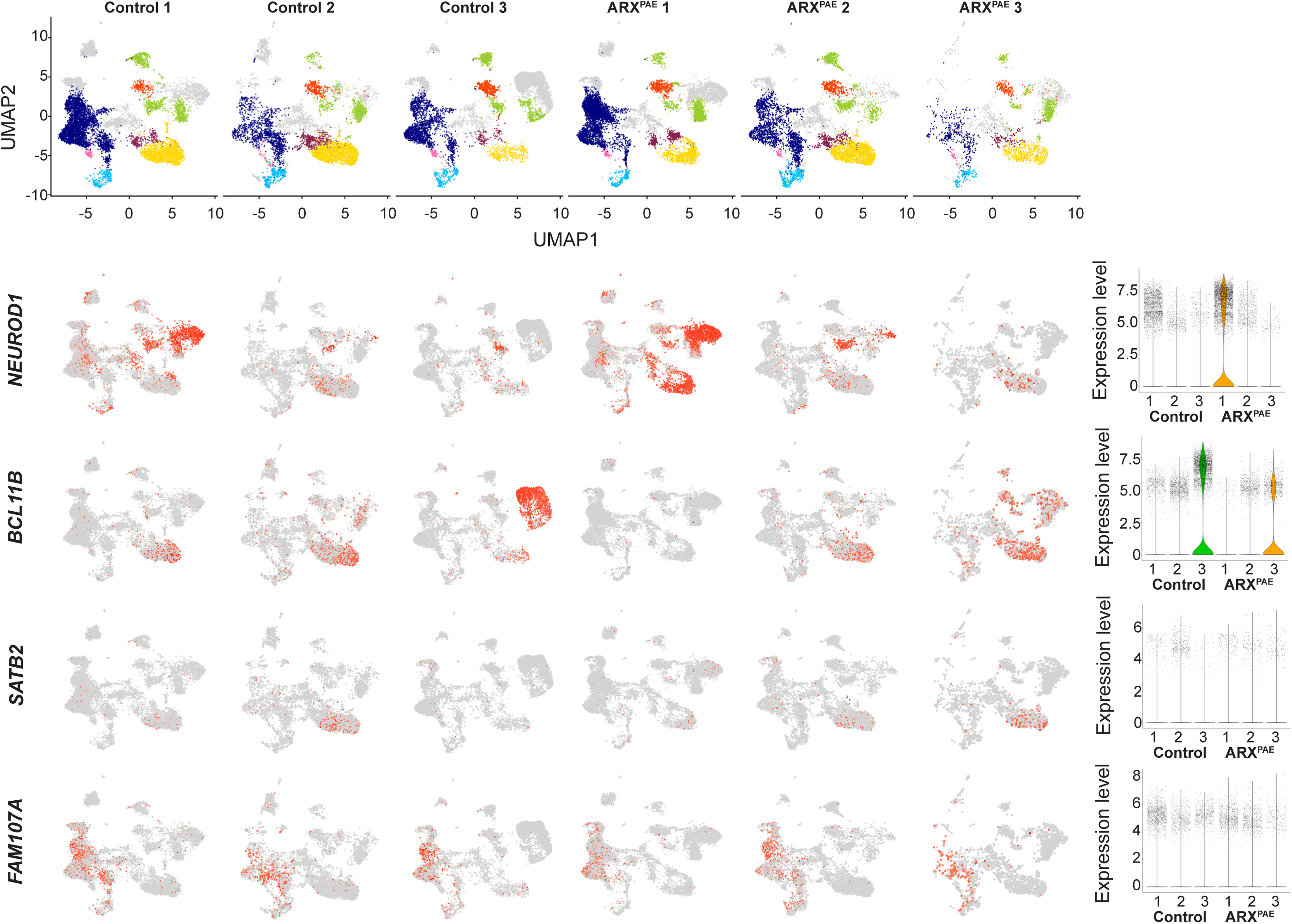
Gene expression in 120 DIV COs by scRNA-seq per line. UMAP plots from control and ARX^PAE^ 120 DIV COs showing the proportion of cells in each cluster and feature plots showing *NEUROD1*, *BCL11B*, *SATB2* and *FAM107A* expression per line. Violin plots show the expression levels of *NEUROD1*, *BCL11B*, *SATB2* and *FAM107A* per line. N= 3-4 organoids from 1 clone x 3 lines per condition.

**Figure S27.**
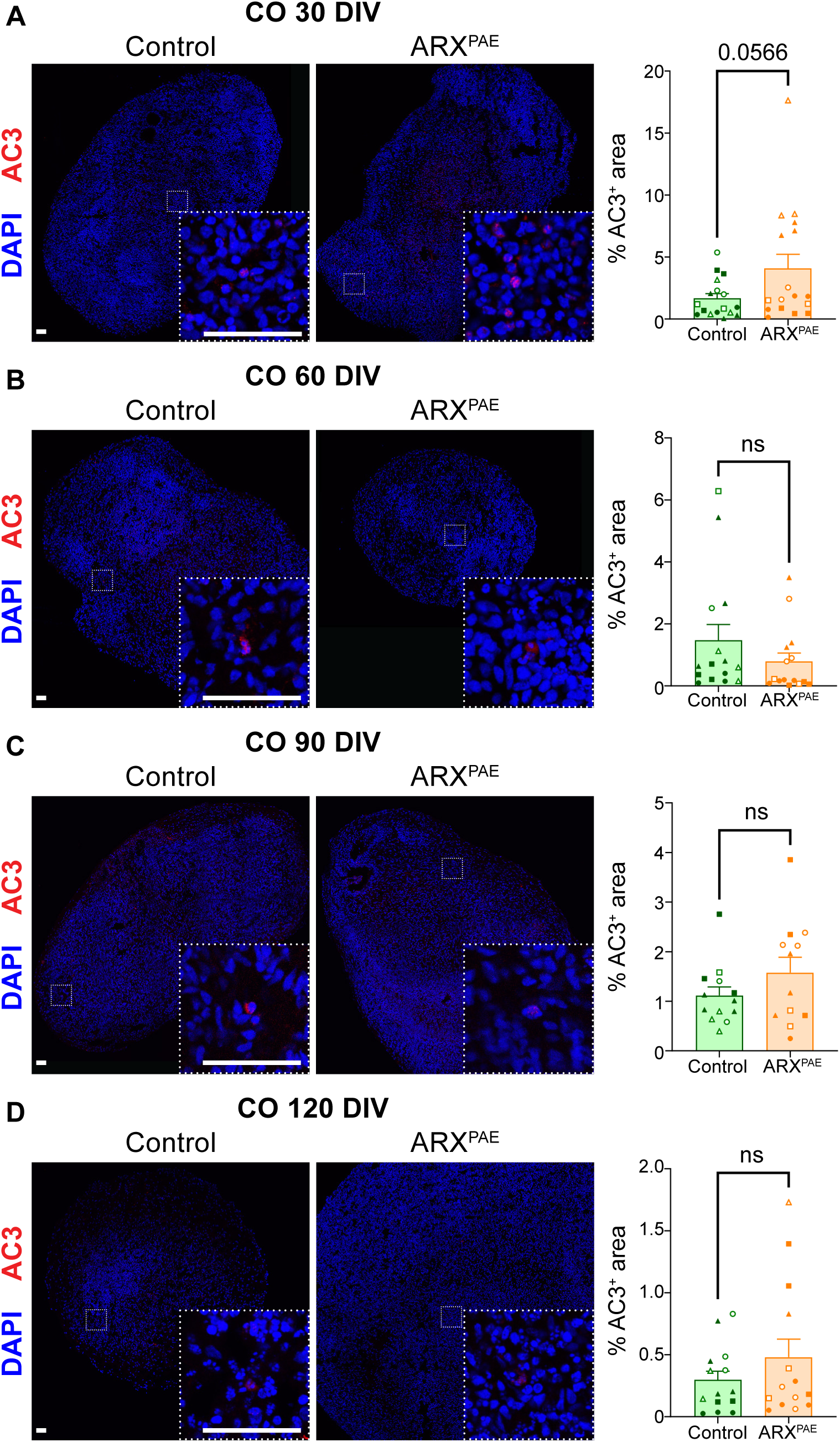
ARX^PAE^ does not promote cell death in COs. **(A)** The images show control and ARX^PAE^ COs at 30 DIV immunostained against AC3 and stained with DAPI. Graph shows the percentage of AC3^+^ area in control and ARX^PAE^ COs at 30 DIV. **(B)** The images show control and ARX^PAE^ COs at 60 DIV immunostained against AC3 and stained with DAPI. Graph shows the percentage of AC3^+^ area in control and ARX^PAE^ COs at 60 DIV. **(C)** The images show control and ARX^PAE^ COs at 90 DIV immunostained against AC3 and stained with DAPI. Graph shows the percentage of AC3^+^ area in control and ARX^PAE^ COs at 90 DIV. **(D)** The images show control and ARX^PAE^ COs at 120 DIV immunostained against AC3 and stained with DAPI. Graph shows the percentage of AC3^+^ area in control and ARX^PAE^ COs at 120 DIV. Scale bar = 50 µm. DIV = days *in vitro*, COs = cortical organoids. Two-tailed Student’s *t*-test, ns = not significant. The results are the mean ± SEM from N= 18 organoids from 2 clones x 3 lines per condition. Individual points represented data from individual organoid and each symbol represented the data from each clone/line (Line 1: Clone A ● Clone B ○; Line 2: Clone A ◼ Clone B ◻; Line 3: Clone A ▲ Clone B △).

**Table S1:** hiPSC lines used.

| Subject | Gender | Age | Number of Poly-Ala | Clinical summary |
| --- | --- | --- | --- | --- |
| **Control 1** | Male | 5.5 y | PA2: 12 | Healthy |
| **Control 2** | Male | 7.5 y | PA2: 12 | Healthy |
| **Control 3** | Male | 35 y | PA2: 12 | Healthy |
| **Patient 1** | Male | 5.5 y | PA2: 12 + 8 | Intellectual disability, medically refractory epilepsy |
| **Patient 2** | Male | 17 y | PA2: 12 + 8 | Intellectual disability |
| **Patient 3** | Male | 42 y | PA2: 12 + 8 | Intellectual disability, epilepsy |

**Table S2:** List of primary antibodies.

| **Antibody** | **Host Species** | **Company** | **Cat. Number** | **Dilution** |
| --- | --- | --- | --- | --- |
| **Anti-AC3** | Rabbit | Cell Signaling | 9661 | 1:400 |
| **Anti-ARX** | Rabbit | Gift from Drs. Morohashi and Kitamura |  | 1:500 |
| **Anti-CB** | Mouse | Swant | CB300 | 1:1000 |
| **Anti-CR** | Mouse | Swant | CR6B3 | 1:1000 |
| **Anti-CTIP2** | Rat | Abcam | AB18465 | 1:300 |
| **Anti-CXCR4** | Rabbit | Invitrogen | PA3-305 | 1:1000 |
| **Anti-DCX** | Guinea pig | Millipore | AB2253 | 1:1000 |
| **Anti-FAM107A** | Rabbit | Sigma | HPA055888 | 1:500 |
| **Anti-GABA** | Rabbit | Sigma | A2052 | 1:1000 |
| **Anti-GFP** | Chicken | AvesLab | GFP-120 | 1:500 |
| **Anti-HOPX** | Rabbit | Sigma | HPA030180 | 1:1000 |
| **Anti-Ki67** | Mouse | BD | 550609 | 1:500 |
| **Anti-LIN28** | Rabbit | Cell Signaling | 3978 | 1:1000 |
| **Anti-NANOG** | Mouse | Thermoscientific | MA1-017 | 1:500 |
| **Anti-NESTIN** | Mouse | Millipore | MAB353 | 1:250 |
| **Anti-NKX2-1** | Rabbit | Abcam | ab133737 | 1:500 |
| **Anti-NEUROD1** | Goat | Santa Cruz | sc-1084 | 1:500 |
| **Anti-mCherry** | Mouse | Invitrogen | M11217 | 1:1000 |
| **Anti-Oct3/4** | Mouse | Santa Cruz | sc-5279 | 1:1000 |
| **Anti-PAX6** | Rabbit | Biolegend | 901301 | 1:300 |
| **Anti-PV** | Mouse | Millipore | MAB1572 | 1:400 |
| **Anti-SATB2** | Rabbit | Abcam | AB34735 | 1:500 |
| **Anti-SOX2** | Rabbit | Millipore | AB5603 | 1:1000 |
| **Anti-SST** | Rat | Millipore | MAB354 | 1:250 |
| **Anti-TBR1** | Rabbit | Abcam | AB31940 | 1:1000 |
| **Anti-TUJ1** | Mouse | Sigma | T8660 | 1:400 |
| **Anti-Vimentin** | Chicken | Abcam | Ab24525 | 1:500 |

**Table S3:**
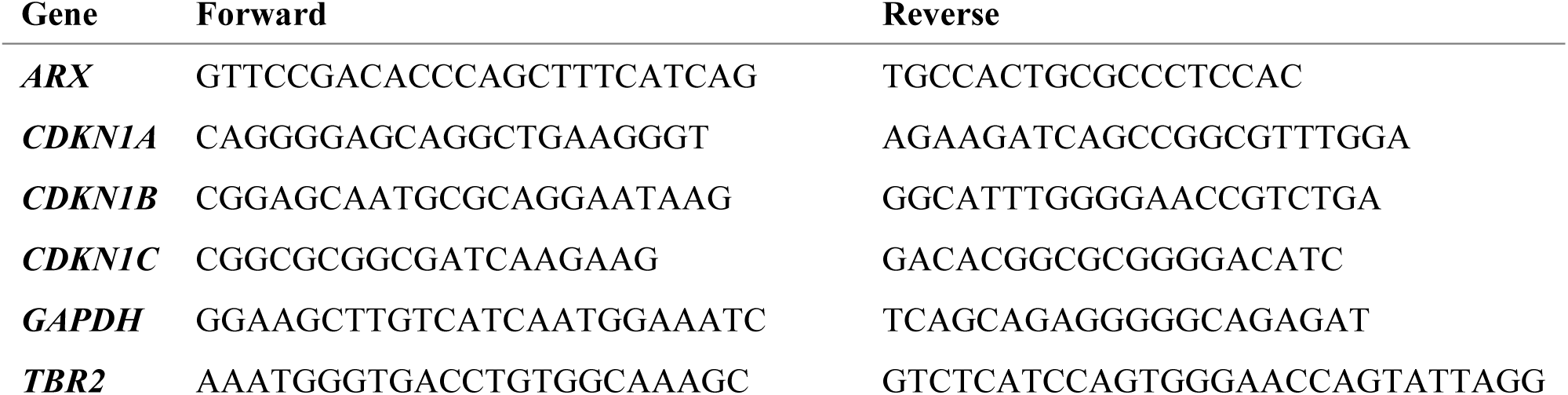
List of primers for RT-qPCR.

**Video 1: Migration of DLX1/2-GFP^+^ interneurons in control assembloids. Example 1**

The video show DLX1/2-GFP^+^ interneurons migrating in the CO side of a control assembloid.

**Video 2: Migration of DLX1/2-GFP^+^ interneurons in control assembloids. Example 2**

The video show DLX1/2-GFP^+^ interneurons migrating in the CO side of a control assembloid.

**Video 3: Migration of DLX1/2-GFP^+^ interneurons in ARX^PAE^ assembloids. Example 1**

The video show DLX1/2-GFP^+^ interneurons migrating in the CO side of a ARX^PAE^ assembloid.

**Video 4: Migration of DLX1/2-GFP^+^interneurons in ARX^PAE^ assembloids. Example 2**

The video show DLX1/2-GFP^+^ interneurons migrating in the CO side of a ARX^PAE^ assembloid.

